# Characterization of a marine bacteria through a novel metabologenomics approach

**DOI:** 10.1101/2024.08.11.607463

**Authors:** Gabriel Santos Arini, Tiago Cabral Borelli, Elthon Góis Ferreira, Rafael de Felício, Paula Rezende Teixeira, Matheus Pedrino, Franciene Rabiço, Guilherme Marcelino Viana de Siqueira, Luiz Gabriel Mencucini, Henrique Tsuji, Lucas Sousa Neves Andrade, Leandro Maza Garrido, Gabriel Padilla, Alberto Gil-de-la-Fuente, Mingxun Wang, Norberto Peporine Lopes, Daniela Barretto Barbosa Trivella, Leticia Veras Costa Lotufo, María-Eugenia Guazzaroni, Ricardo Roberto da Silva

## Abstract

Exploiting microbial natural products is a key pursuit of the bioactive compound discovery field. Recent advances in modern analytical techniques have increased the volume of microbial genomes and their encoded biosynthetic products measured by mass spectrometry-based metabolomics. However, connecting multi-omics data to uncover metabolic processes of interest is still challenging. This results in a large portion of genes and metabolites remaining unannotated. Further exacerbating the annotation challenge, databases and tools for annotation and omics integration are scattered, requiring complex computations to annotate and integrate omics datasets. Here we performed a two-way integrative analysis combining genomics and metabolomics data to describe a new approach to characterize the marine bacterial isolate BRA006 and to explore its biosynthetic gene cluster (BGC) content as well as the bioactive compounds detected by metabolomics. We described BRA006 genomic content and structure by comparing Illumina and Oxford Nanopore MinION sequencing approaches. Digital DNA:DNA hybridization (dDDH) taxonomically assigned BRA006 as a potential new species of the *Micromonospora* genus. Starting from LC-ESI(+)-HRMS/MS data, and mapping the annotated enzymes and metabolites belonging to the same pathways, our integrative analysis allowed us to correlate the compound Brevianamide F to a new BGC, previously assigned to other function.

## Introduction

The search for new bioactive compounds of natural origin from different organisms coming from different biomes is an arduous task. Since marine environments are poorly explored, they hold the promise of a formidable source of rich metabolic potential for the production of novel biosynthetic compounds, especially when you consider the microorganisms that reach a billion strains in a gram of marine sediment (1). Considering that more than 40% of pharmaceutical ingredients are derived directly or indirectly from natural products derived from plants or microorganisms, one can expect that thousands of unknown potential medicines are expected to be discovered in marine ecosystems (2–3). This feature places Brazil under the spotlight since its coast is especially large, ranging from tropical to temperate climate zones (4).

Within the phylum Actinomycetota, the genus *Micromonospora* is distributed in different regions of the world, mainly in soil and marine environments (5). The *Micromonospora* genus is composed of 177 Gram-positive, spore-forming aerobic species and is ubiquitous in marine environments. They belong to the *Actinomycetota*, a diverse phylum responsible for 70% of natural compounds under development or already in clinical use. The chemical diversity, in terms of natural products, that this genus is capable of producing is enormous. *Micromonospora* natural products are used as drugs against infections caused by fungi or bacteria. The genera started to receive attention after the discovery of gentamicin in 1963 and after that, more than 740 bioactive compounds have been reported from *Micromonospora* strains. Among this chemical diversity produced, as well as the different locations where this genus can be found, there are reports in the literature searching specifically for anticancer bioactive compounds on *Micromonospora* sp. BRA006 (6).

The evolution of bacterial genome organization clustered genes that encode enzymes of the same metabolic pathway, which are known as biosynthetic gene clusters (BGC) (7). For this reason, modern drug discovery from bacteria is based on BGC identification as the starting point followed by an experimental procedure that aims to detect, isolate, or produce the compound (8). However, searching for novel natural bioactive compounds from microorganisms can be a harsh task, mostly because the majority of the microbial life cannot be cultured under laboratory conditions. Also, it is difficult to obtain bioactive compounds in the desired concentrations (9). Thus, bacterial bioactive compound discovery requires a multidisciplinary approach, such as genomics and metabolomics (metabologenomics) (10). Genomics enables the analysis of the whole genome sequencing data and raises hypotheses about metabolic pathways and compound products based on the genetic content (11). Then, metabolomic assays based on mass spectrometry analysis, such as LC-MS/MS are performed to validate them. The integration of data in the description of new bacterial strains has been shown as a powerful resource, as it allows the study and characterization of new microorganisms in a holistic way (12).

In the present work, we used metabologenomics to describe the potential bioactive compounds of BRA006, a bacteria strain recovered from a marine environment collected on the coast of Brazil. We performed our analysis in a two-way direction, searching for metabolites by LC-MS/MS previously predicted by antiSMASH, using two genome sequencing platforms, as well as finding coding sequences (CDS) for enzymes that are part of metabolic pathways for syntheses of BRA006’s observed metabolome.

## 2. Results

### 2.1 Genomic Analysis

The isolate BRA006 exhibits very characteristic growth. The colonies are orange in color with apical growth and a rough appearance with the presence of individual dark-colored spores. In solid medium, it releases as yet undetermined compounds that diffuse into the agar, giving it a pink-purple color, as can be seen in Supplementary Figure 1.

We compared the Illumina and MinION whole genome sequence techniques results from BRA006. While Illumina assembly yielded 10 contigs and 6,734,372 base pairs (6.73 Mb) in total, sequencing by MinION showed higher genome completeness, with an assembly resulting in 4 contigs with 6,762,267 base pairs (6.76 Mb) in total. Functional analysis (Table 1) performed by Prokka showed a significant increase in CDS predicted from MinION assembly data. The complete result is shown in Supplementary Tables 1 and 2. The CDS prediction followed the number of hypothetical proteins, with more predictions from MinION than from Illumina.

**Table 1.**
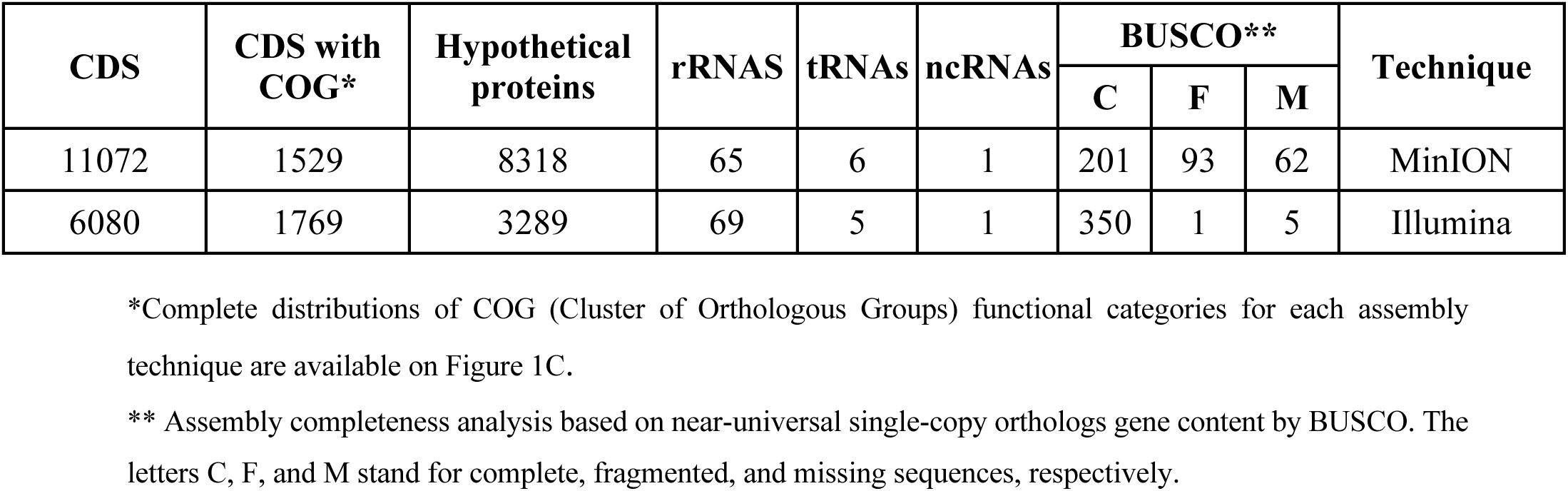
Functional genomic features.

| CDS | CDS with COG* | Hypothetical proteins | rRNAs | tRNAs | ncRNAs | BUSCO** |  |  | Technique |
| --- | --- | --- | --- | --- | --- | --- | --- | --- | --- |
|  |  |  |  |  |  | C | F | M |  |
| 11072 | 1529 | 8318 | 65 | 6 | 1 | 201 | 93 | 62 | MinION |
| 6080 | 1769 | 3289 | 69 | 5 | 1 | 350 | 1 | 5 | Illumina |

\*Complete distributions of COG (Cluster of Orthologous Groups) functional categories for each assembly technique are available on Figure 1C.
\*\* Assembly completeness analysis based on near-universal single-copy orthologs gene content by BUSCO. The letters C, F, and M stand for complete, fragmented, and missing sequences, respectively.

### 2.2. Metabolic potential

The *Micromonospora* is the largest genera of Actinomycetota and possesses a large repertoire of bioactive secondary metabolites (SM) with a broad spectrum of therapeutic effects (13), for instance, aminoglycosides and macrolactam antibiotics. Through antiSMASH, we annotated the BGC content from Illumina and MinION assemblies of BRA006. MinION assembly resulted in a total of 15 BGCs (Table 2) that vary in similarity with antiSMASH database. Among them, there are those with reported antimicrobial, antifungal, and antitumor activities. For instance, quinolidomicin A is a macrolide with antibiotic and anticancer effects isolated from *Micromonospora sp* JY16 (7) and BRA006 presents a quinolidomicin BGC with 219 Mb in length and 67% similarity with the most similar known cluster.

**Table 2.**
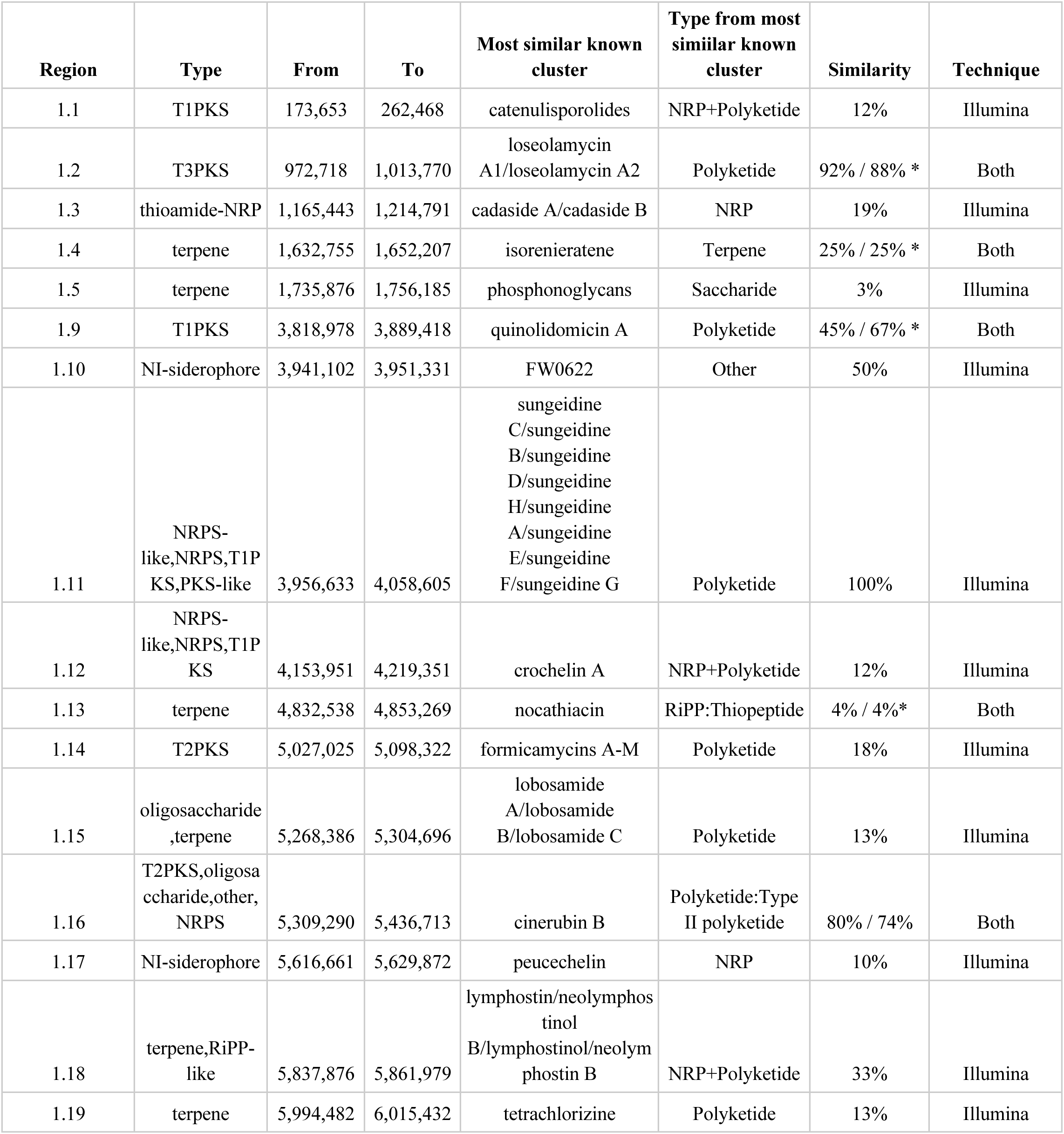
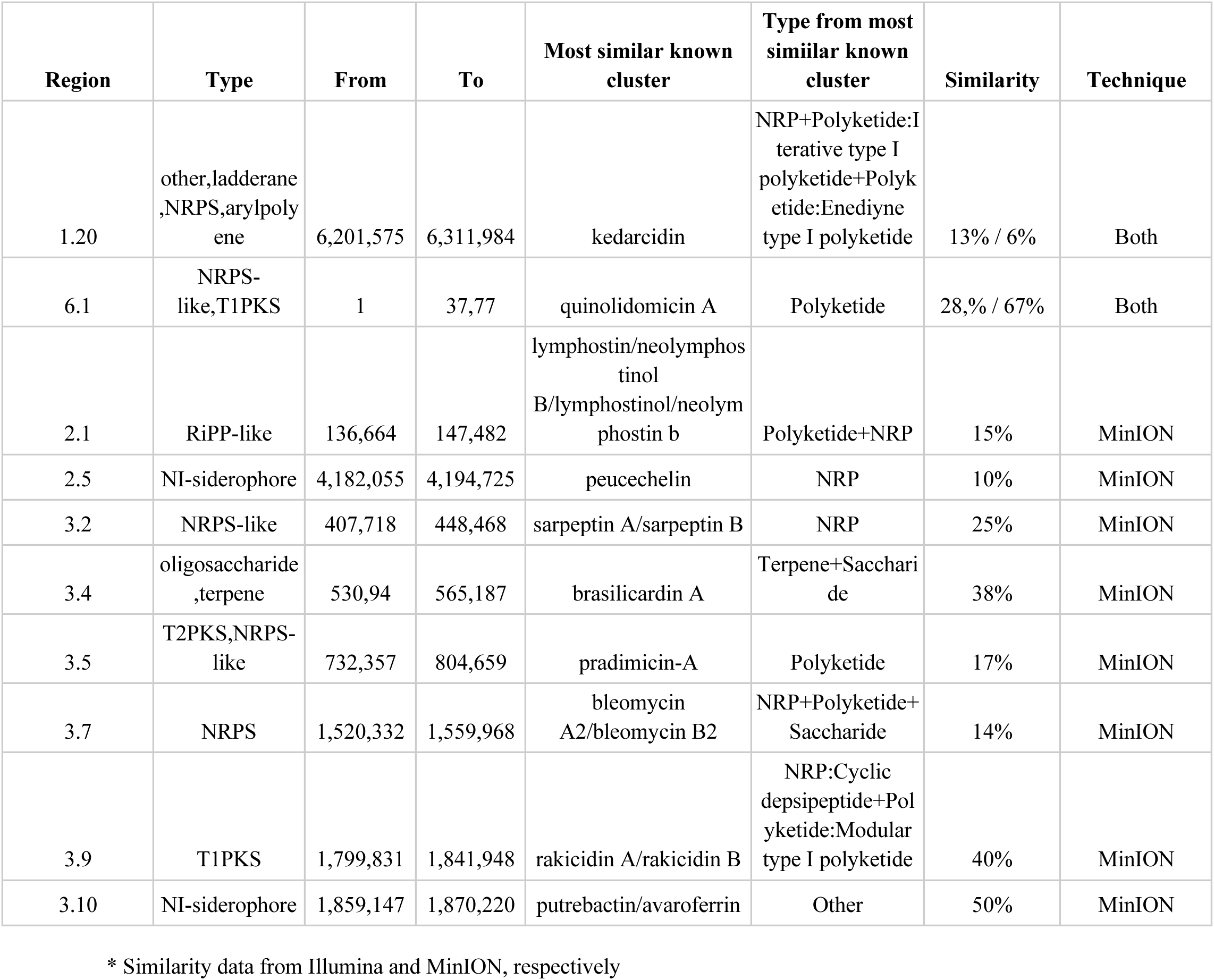
Biosynthetic Gene Cluster from antiSMASH.

The type-III polyketide Loseolamycin was identified from *Micromonospora endolithica* and inhibited the growth of the Gram-positive *Bacillus subtilis* and also showed herbicidal activity against the weed *Agrostis stolonifera* (14). Cinerubins are anthracycline antimicrobials produced by actinomycetota which also present antitumor activity (15–16). BRA006 possesses highly similar clusters to cinerubin B (74% by MinION and 80% by Illumina) and loseolamycin (88% by MinION and 92% by Illumina) A1/A2. BRA006 also has less similar BGCs for the biosynthesis of other natural compounds with antitumoral activity such as bleomycins and kedarcidin, isolated from *Streptomyces verticillus* and an unclassified Actinomycetales strain (ATCC 53650) *Actinomycetes,* respectively (17–18).

AntiSMASH results from Illumina assembly yielded 18 BGCs (Table 2) in total, including clusters for the production of Quinolidomicyn, Cinerubin B, and Loseolamycin A1/A2 found in both Illumina and MinION data. However, only by sequencing with Illumina, it was possible to find a BGC 100% similar to the production of Sungeidines (19), a group of metabolites produced by pathways with close evolutionary relations with the antitumor Dynemicins (20).

### 2.3. Evolutionary relationships

Since both genome sequencing methods yielded identical clusters with enzymes from pathways for synthesizing compounds with medical applications, we decided to explore the evolutionary relationship. According to digital DNA:DNA hybridization used by Type Genome Server (TYGS) (21) for phylogenetic inferencing, these two assemblies were classified as *Micromonospora spp* and pointed out as possible novel species due to their relatively high genomic distance to its closest related group: *Micromonospora aurantiaca* ATCC 27029 (Figure 2). However, the BGC with higher similarity to the antiSMASH database is the one for the production of Loseolamycin A1/A2 in both MinION and Illumina assemblies. Therefore we used BLASTp to compare the proteins within those BGCs to the reference: BGC0002362 from *Micromonospora endolithica*.

Figure 1B shows the protein similarity between loseolamycin-producing BGC from BRA006 and *M. endolithica.* Both assembly methods detected the complete inversion of this *M. endolithica* BGC followed by a series of indels in the upstream region majorly composed of CDSs that encode proteins with no functional annotation. Other BGCs with less similarity to the antiSMASH database were also compared with the reference sequences. For the cinebubin B and quinalidomicin A BGCs there were differences between antiSMASH results regarding the sequencing method. In the case of cinerubin B BGC, sequencing by Illumina resulted in a BGC with 172 Mb length while MinION’s with 71 Mb. Both with a few highly similar proteins to the reference BGS (BGC0000212) (Supplementary Figure 2) original from *Streptomyces sp. SPBO074*. On the other hand, quinolidomicin A BGCs showed 219 Mb in MinION and 70 Mb in Illumina assembly. Quinolidomicin A BGC from MinION data presented more similarity (67% against 45% from the counterpart) to reference BGC0002520 original from *Micromonospora sp* (Supplementary Figure 3).

**Figure 1.**
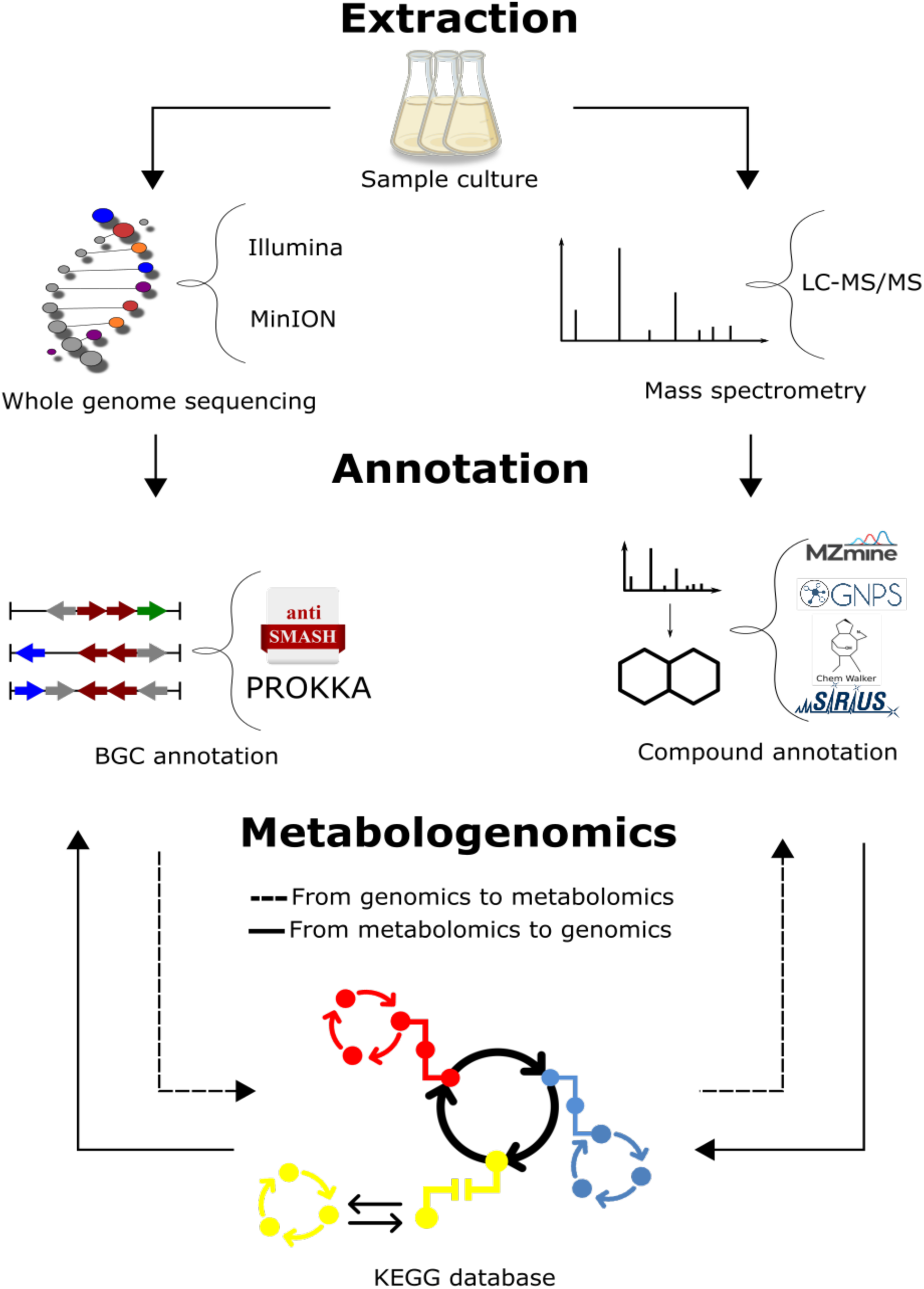
Metabologenomics workflow. LC-MS/MS raw data was pre-processed using MZmine3. Spectral pairing and molecular network construction with GNPS2. in silico annotations were performed with ChemWalker and SIRIUS. The annotated compounds were used to search KEGG pathway database. KEGG pathway matches had the enzymes serarched in the genome. The genome assemblies for MInION and Illumina had gene annotation performed by Prokka and AntiSMASH. From the prediction of metabolites, we searched the annotated metabolome.

**Figure 2.**
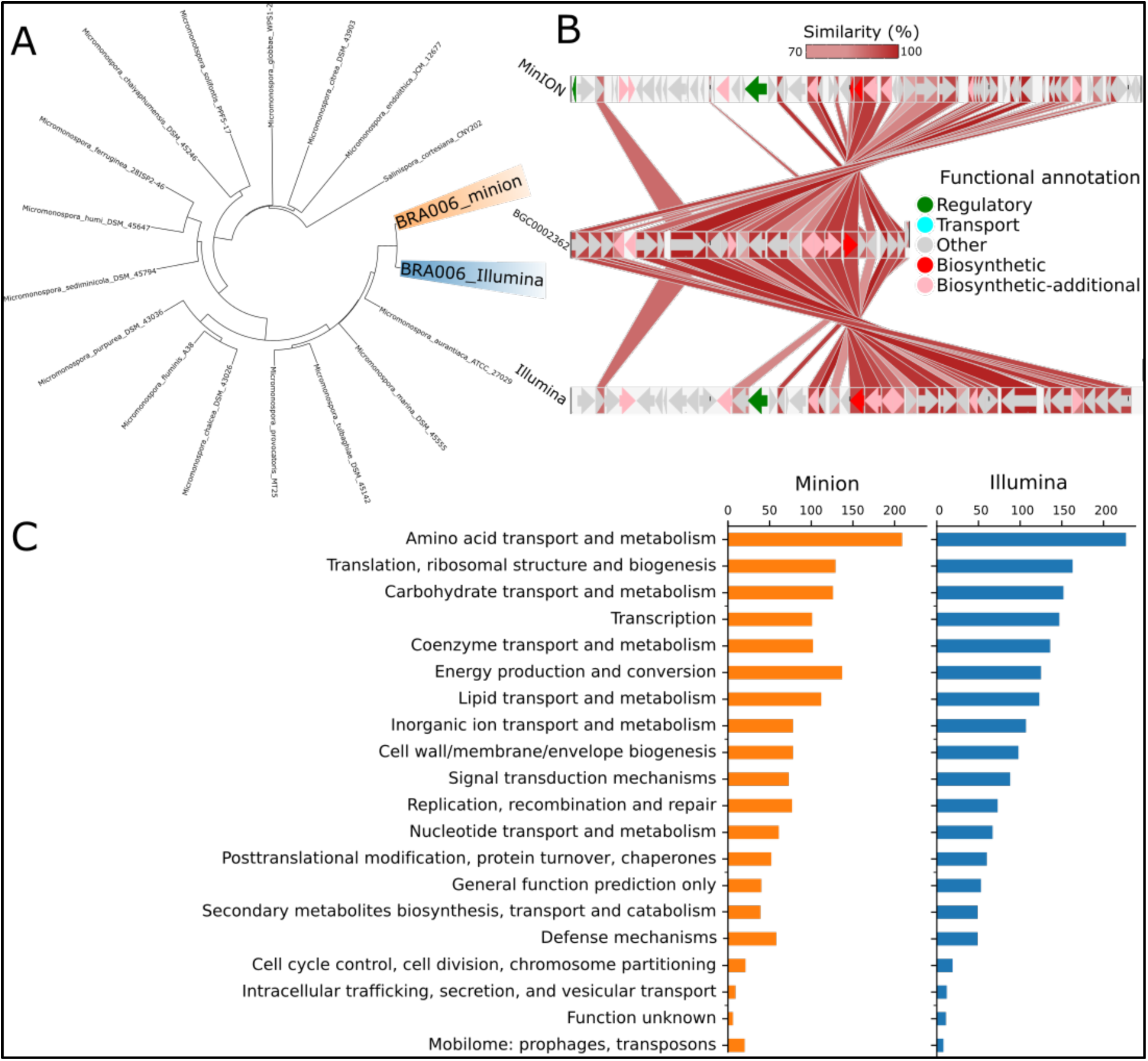
(A) Digital DNA:DNA hybridization phylogenetic tree of the Illumina and MinION data from the isolate BRA006. The Type Genome Server displays the 16 closest related genomes present in its database based on the genomic distance of the whole genome sequencing data. (B) Protein BLASTp similarity between BRA006 Loseolamycin BGC and reference BGC0002362 point by antiSMASH. All possible matches for every BRA006 match were filtered by the smallest e-value (See complete BLAST data in Supplementary Tables 1, 2 and 3). (C) The distribution of CDS annotated by Prokka according to Cluster of Orthologous Groups.

### 2.4. Metabolomic Analysis

From the analysis of potential BGCs found in BRA006, where their potential to biosynthesize compounds with antibacterial (loseolamycins A, quinolidomicin A, cinerubin B, and brasilicardin A), antifungal (pradimicin A) and anticancer (bleomycin A2 and kedarcidin) activity was identified, we investigated whether these compounds were present in the metabolome of this Actinomycetota. To do so, and to extend the analysis to other compounds that BRA006 might be able to produce, we performed a metabolomics analysis in which the BRA006 extract was analyzed by LC-ESI(+)-HRMS/MS. After the LC-MS/MS analysis, the obtained raw data were converted into .mzXML format using Proteowizard (22), and feature finding was performed using MZmine (23). We then performed the annotations sequentially, in three steps. In the first, the annotations were made by spectral pairing and molecular network construction through GNPS2 (https://gnps2.org/). As a result, we obtained 527 nodes with at least one connection, of which 83 were annotated by GNPS2. Based on this result, we propagated the annotations to the nodes that did not present an annotation by spectral pairing using ChemWalker (24), increasing the annotations to 373 nodes. Finally, for the nodes that could not be annotated by this tool, we used a third *in silico* spectral annotation tool, SIRIUS (25), increasing the annotations for an additional 67 nodes. Thus, a total of 527 nodes were represented by the molecular network, of which 523 were annotated. With these results in hand, we performed an automated chemical classification of each annotated compound using ClassyFire (26). The molecular network was colored based on the superclasses to which each annotated compound belonged (Fig. 3).

**Figure 3.**
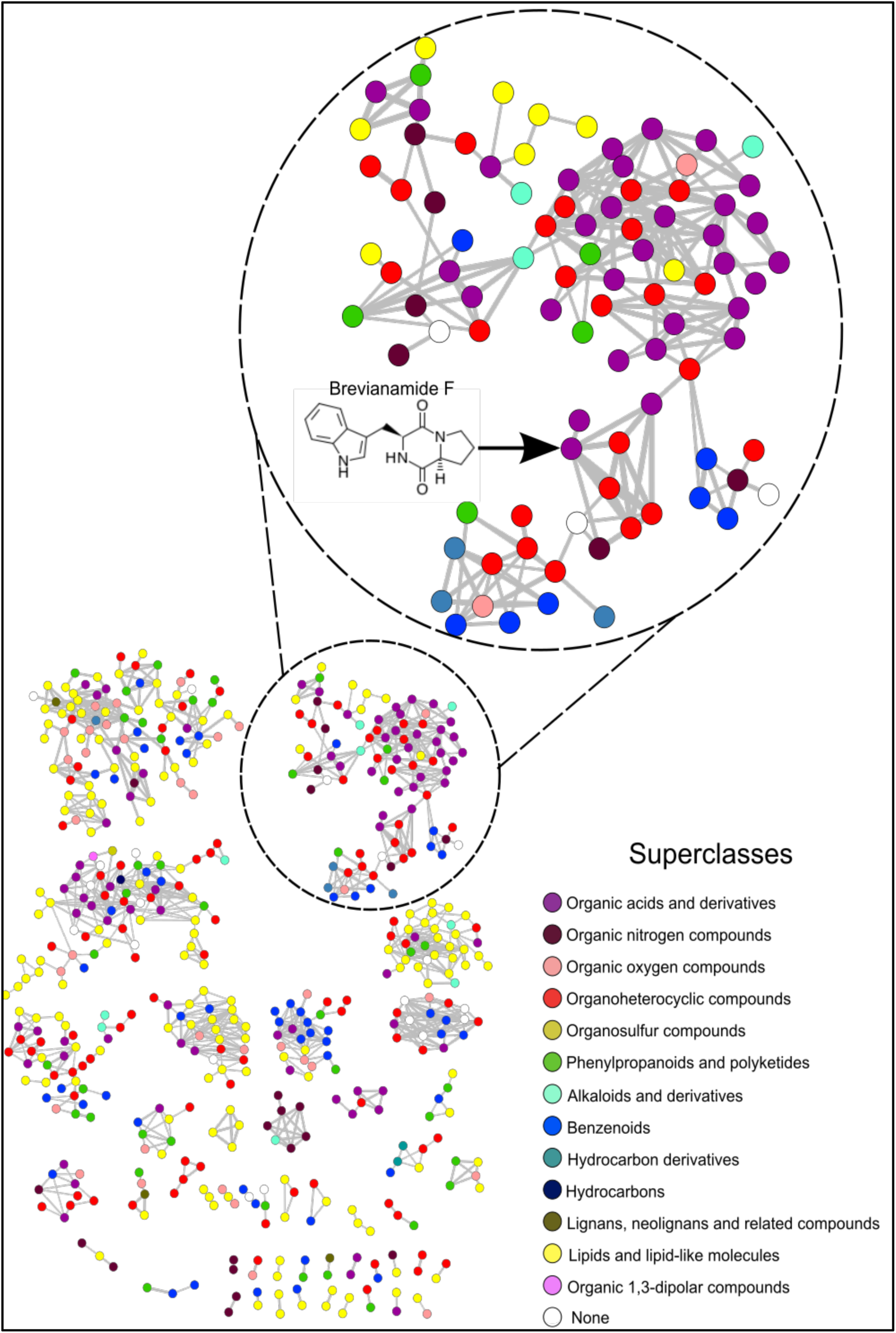
Molecular network constructed from BRA006 metabolomic data. Nodes were colored based on the superclass classification performed on the molecules annotated by ClassyFire. The full description of the annotation set is presented in Supplementary Table 4. The cluster showing the node with the annotation for brevianamide F is highlighted.

The compounds could be grouped into 13 different chemical superclasses, where 30.2% of the annotated compounds belong to the superclass of lipids and lipid-like molecules, 19.5% to organoheterocyclic compounds, 15.1% to organic acids and derivatives, 9.9% to benzenoids, 7.6% to phenylpropanoids and polyketides, 6.5% to organic oxygen compounds, 4.0% to organic nitrogen compounds, 1.7% to alkaloids and derivatives, 0.6% to lignans, neolignans, and related compounds, 0.4% to hydrocarbon derivatives, 0.2% to organic 1, 3-dipolar compounds, 0.2% to hydrocarbons, 0.2% to organosulfur compounds, and 3.8% could not be classified (None). The complete relationship between all annotated compounds and their chemical hierarchical classification into kingdom, superclass, and class is shown in Supplementary Table 4. From the entire set of annotations and classifications, we searched for the seven compounds predicted by antiSMASH in the BRA006 metabolome. None of the seven compounds were found in the annotation hall. Therefore, we took the molecular structure of these seven compounds and performed their chemical classification using ClassyFire; once we had their chemical class, we would search for compounds annotated in the BRA006 metabolome that had the same chemical class. From a pharmacological point of view, compounds belonging to the same chemical class might belong to the same biosynthetic pathway and or have similar effects, as is the case, for example, with the class of peptidomimetics in the treatment of cancer (27) and steroids in the treatment of pain (28). The seven compounds were grouped into six different classes of molecules, where loseolamycin A1 belongs to the phenol class, cinerubin B belongs to the anthracycline class, brasilicardin A belongs to the steroid and steroid derivative class, pradimicin A belongs to the naphthacene class, bleomycin A2 belongs to the peptidomimetic class, and quinolidomycin A and kedarcidin are organooxygen compounds.

Of these six chemical classes to which the compounds of interest belong, the Phenols class presented 10 annotated molecules, while the Organooxygen Compounds and Steroids and Steroid Derivatives classes presented 34 compounds each and Peptidomimetics presented two compounds in our analysis. Among the 10 compounds belonging to the Phenol class, three of them had bioactivity previously reported, but with a different action from the antibiotic loseolamycins A1. In the case of the compounds found, lumizinone A was described as having inhibitory activity on proteases (29), rubrolide R showed cytotoxic activity against tumor cells (30), while hierridin C showed antimalarial activity (31). In turn, of the 34 molecules belonging to the organooxygen compounds, six showed bioactivity reported in the literature. Pyrenocine B was described for its immunosuppressive potential (32), while subamolide A and surinone C were described for their cytotoxic activity (33–34). Talaroenamine B showed evidence of antiplasmodial activity (35), while moniliferanone B was described for its antibiotic potential (36), the same activity described for quinolidomycin A as predicted by antiSMASH. For the 34 compounds belonging to the class of steroids and steroid derivatives, seven showed biological activity described in the literature. Of these, veramiline has been described for its potential in the therapy of dengue infection (37). Certonardosterol D3 has been described for its anticancer potential (38), while vaganine D has been described for its cholinesterase inhibitory activity (39). Among the others, 6-ketolithocholic acid was described for its action in suppressing bile acid production (40), cholic acid for its action in cholesterol reduction (41), and gitoxigenin showed therapeutic effects in congestive heart failure (42). Finally, of the two compounds detected in the metabolome belonging to the class of peptidomimetics, pseudodestruxin A has been described for its antibacterial activity (43). The targeting offered by antiSMASH, by searching for molecules belonging to the same class as those with bioactive activity predicted by the tool, allowed us to find a larger and more diverse range of compounds.

Using a reverse flow of integrative analysis, where we start from what was annotated in the metabolome, we set out to evaluate whether it would be possible to identify, from a given metabolite, the enzymes that lead to its production in BRA006. To this end, we first crossed the annotated metabolome with the KEGG database (44) to search within the metabolome of this Actinomycetota for molecules with known biosynthetic pathways. As a result, we found a molecule belonging to the staurosporin biosynthetic pathway, brevianamide F. It should be noted that of the three methods used in the annotation process, brevianamide F was annotated by spectral pairing with the GNPS spectral library (45), showing a MQScore of 0.97469 and a m/z error of 2.47028 ppm. Since the topology of the molecular network is given by the similarity between nodes, the node with an annotation for brevianamide F is connected to seven other nodes (Figure 3B). All seven nodes have been annotated by ChemWalker. Looking at the predicted structures, six of the seven share the same indole-like nucleus as brevianamide F, suggesting that other molecules may ultimately be produced, either by brevianamide F BGC or by other intermediates belonging to the same pathway as brevianamide F. Once identified, the enzymes that make up the staurosporin biosynthetic pathway, we searched the BRA006 genome for which of these enzymes would be encoded. From this search, 16 staurosporin pathway enzymes were identified in the genome of this Actinomycetota (Supplementary Figure 4), and 11 overlap the region of the BGC 20 from Illumina (Figure 4). Among them, we found an NRPK 2,3-dihydroxybenzoate-AMP ligase functionally classified as a biosynthetic-additional enzyme by antiSMASH. These results show that a dynamic integrative approach, i.e., first combining spectral and *in silico* annotation, assigning chemical classes, and then searching for metabolites from the genome to match encoded proteins in the genome to the pathway producing the putative metabolite annotated, is an efficient approach to characterize new species with the potential to produce bioactive compounds.

**Figure 4.**
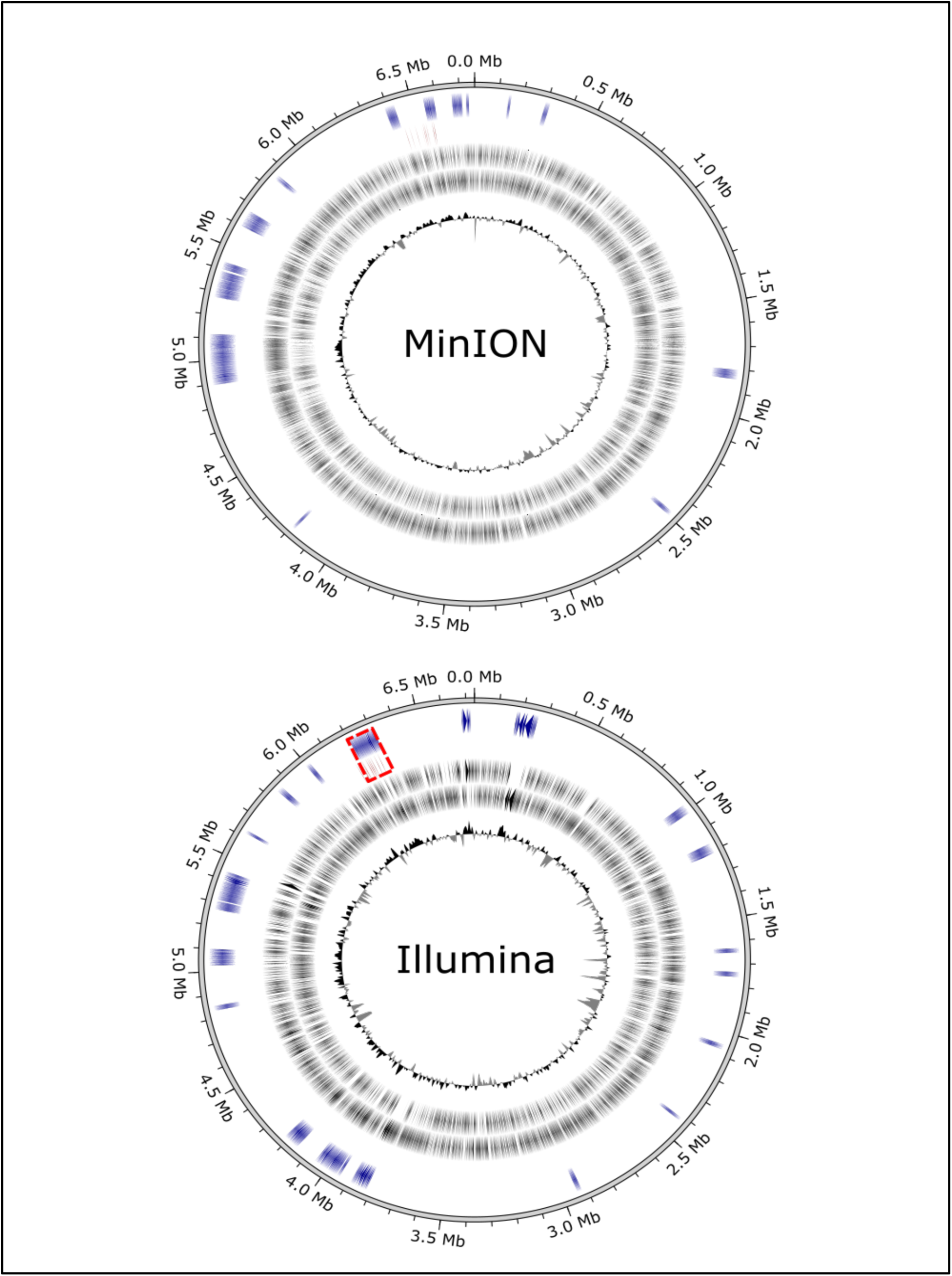
Circular representation of BRA006 genome according to the sequencing technique. The first circle (from inside to outside) GC content, the gray circles represent forward and reverse CDS positions, respectively, the red circles represent CDSs that encode proteins present in the KO000404 pathway (https://www.genome.jp/pathway/ko00404), and the blue circles represent the CDSs from BGCs. The highlighted area (in red) shows the overlap between BGC 20 and proteins present in the KO00404.

## 3. Discussion

The metabologenomic characterization of new *Micromonospora* strains, especially those found in the marine environment, is a highly relevant task given the potential for the discovery of new bioactive compounds. In this sense, the use of dynamic integrative analytical approaches seems to be a promising resource, both in the process of characterizing new species and in the evaluation of the potential for the production of natural products from a new microorganism. In the present work, we characterized BRA006, a potential new species of the genus *Micromonospora* in terms of secondary metabolites production. We used both genomic and metabolomic points of view, comparing two whole genome sequencing approaches and a multiple-step metabolite annotation workflow. This strategy introduces an innovative dynamic approach to metabologenomic analysis in which the annotated BGCs types predicted by antiSMASH guided the search for bioactive compounds in the metabolome content, as well as the compounds identified in the metabolome (which match specific pathways) led to the search for CDSs that encode enzymes from these pathways. Among the various microbial genera described to date that stand out for their ability to produce bioactive compounds, the genus *Micromonospora* is an important model in natural products research and a milestone in the discovery of new biocompounds (5). The potential to produce bioactive natural products from bacterial isolates from the Brazilian coast is already known, especially those with antitumor activity (6,46). Among these, the activity observed in crude extracts of the genus *Micromonospora* was attributed to a group of anthracyclinones (6).

According to (13), species from the *Micromonospora* genus encode from 4200 to 8017 proteins and have genome sizes from 5.07 to 9.24 Mb. BRA006 possesses a 6.7 Mb genome, confirmed by two independent sequencing methods, although MinION assembly annotated by Prokka showed up to 11.000 CDS against 6080 from Illumina. This gap between them is mostly due to higher error rates from MinION assembly, which probably causes artificial stop codons that could explain the higher number of CDSs. A proper way to solve this issue would be to perform a hybrid assembly (47–48). However, this question needs further investigation. Besides the difference in CDS number, antiSMASH found the same BGCs in both assemblies, such as Cinerubin B and Loseolamycins, with high similarity to antiSMASH database. Also, the whole genome sequence phylogeny placed both assemblies as a monophyletic group dissimilar enough from *Micromonispora aurantiaca* ATCC27029 to TYGS to point BRA006 as a possible novel *Micromonospora* species. As an example of genetic divergence of BRA006 from other *Micromonospora,* we can cite the Loseolamycin A1 BGC, where it is possible to observe a complete inversion of the cluster and several indels. According to Medema et al., (2014) (49) NRPK clusters evolved from gene duplication followed by differentiation, which could explain the difference between BRA006 and *M. endolithica.* Unfortunately, even with Prokka, annotating most of the proteins in that BGC was not yet possible.

The approach “from genomics to metabolomics” yielded a BGC that encodes pathways to compounds with pharmaceutical applications, which confirms the importance of the *Micromonospora* genus, although the compounds produced by these BGCs were not found in metabolomic data. Their absence can be explained by differences between laboratory culture media and the original ecological niche, as well as the need for improvements in the acquisition parameters. Genomic-guided works often require heterologous expression of parts or the entire BGC to obtain the active compound in the laboratory (50). For instance, Lasch et al. (14) obtained Loseolamycins from *M. endolithica* by heterologously expressing type III polyketide synthase, Domingues Vieira et al. increased the production of Eponemycin and related epoxyketone peptides by cloning the whole *epn-tmc* BGC from *Streptomyces sp.* BRA346 (8), and Yamanaka produced Taraomycin A by editing regulators of this BGC from *Saccharomonospora sp.* CNQ490 (51).

Starting from metabolomics, the analysis of the BRA006 metabolome allowed us to identify annotated compounds belonging to already well-established chemical classes whose biosynthesis is reported in the literature for this genus, as in the case of macrolides (5). Of these, we were able to annotate nine compounds belonging to this class (Supplementary Table 4), two of which previously reported bioactivity: Tricholide A, with antibacterial activity (52) and 11,12-dihydroxy-6,14-dimethyl-1,7-dioxacyclotetradeca-3,9-diene-2,8-dione, with immunosuppressive activity (53). It should be noted that two compounds were recorded as Tricholide A. Both presented the same *m/z* value but with very different retention times, indicating the presence of isomers of this compound, as both appear as neighboring nodes in the molecular network. The annotation procedures and KEGG pathway search identified Brevianamide F, a compound with activity against *Staphylococcus aureus* (54) and an intermediate of well-established inducer of apoptosis, Staurosporin (55). Among the three annotation methods for a given molecule we used, spectral matching against a reference library is the best available resource (56). In addition, we used two other *in silico* annotation resources: ChemWalker and SIRIUS. Of these two tools, SIRIUS has the best accuracy, but it is very difficult to use when dealing with large sets of spectra. This limitation is overcome by ChemWalker, which allows greater annotation coverage, taking into account the topology of the molecular network. We reached brevianamide F annotation through two different annotation routes, which brings robustness to the interpretation of the result obtained and highlights the potential for a reverse flow in elucidating the biosynthetic potential of a new non-model organism. It is worth highlighting that we also carried out an *in silico* prediction analysis using SIRIUS for the seven nodes related to that of Brevianimide F. However, the best candidates predicted by this tool had little structural similarity with the compound itself, unlike the candidates provided by ChemWalker. The advantage of ChemWalker is that it uses the sample context given by the molecular network topology on compound re-ranking. Eventually, this new proposed BGC could synthesize the other molecules annotated and linked to the Brevianamide F’s node or, maybe, they are intermediates from different pathways that had not been described yet. In both cases, the knowledge of these analogues opens the possibilities to improve the known bioactivity of Brevianamide F, which could be tested by isolation or even by (bio)synthesizing the compounds.

Inspecting KEGG’s metabolic pathways, we found that Brevianamide F, a product of fungi metabolism (57), is part of Staurosporin biosynthesis (KO00404). Therefore, we retrieved all EC numbers from the KO00404 pathway, connected them to our Prokka data, and found three enzymes that can catalyze the Brevianamide F synthesis reaction. In the KO00404 pathway, brevianamide F biosynthesis requires tryptophan and proline as substrates, being its core assembled by a non-ribosomal peptide synthetase (COG1020), named Brevianamide F synthase (EC: 6.3.2.-), which is encoded by the gene FtmA (NCBI ID *Aspergillus fumigatus* (AFUA_8G00170) (58), and is also reported in *Streptomyces sp* (59). The isolate BRA006 has an NRPK mbtB_1 (MinION data) that matches with COG and EC number of FtmA, but is not a component of any BGC found by antiSMASH. However, examining the downstream and upstream regions of 2,3-dihydroxybenzoate-AMP ligase CDS (Illumina data) we found a genomic region that has the potential to be part of the brevianemide F synthesis pathway BGC.

AntiSMASH identifies BGCs based on profiles of Hidden Markov Models (pHMM) from PFAM (60), TIGRFAMs (61), SMART (62), BAGEL (63), Yadav et al. 2019 (64) and custom models that recognize signature sequences of such conserved domains in genomic query sequences (65). However, there are BGCs that lack universal class-specific signature sequences and therefore are partially identified. To overcome this limitation, deep-learning-based tools such as DeepBGC (66) have been applied in genomic mining research to uncover new BGCs.

It is interesting to emphasize the innovative approach used in the present work with a two-way analysis of the genome and metabolome. We started with the metabolome to see if the biosynthetic gene clusters involved in the production of a given metabolite could be identified in the genome. We then analyzed the genome sequencing data, and from there, by searching specific databases such as antiSMASH, we went to the metabolome to check whether the compounds predicted by antiSMASH were being produced (8). Traditionally, the search for potential new compounds with bioactivity follows the latter linear flow of analysis, which in our case did not result in the identification of 7 of the metabolites predicted by genome mining. However, by using the chemical classes of these compounds, we could find analogues in our metabolomic data.

In addition, we present a new approach that integrates the classical approach with a reverse analysis, starting from the metabolome to the genome. For example, in neither short-read (Illumina) nor long-read (minION) sequencing data, it was possible to automatically detect the biosynthetic gene cluster for brevianamide F or staurosporin production. By using the two-way approach we could identify brevianamide F (reported as an intermediate in the Staurosporin biosynthesis) in BRA006 metabolome and, from there, we could identify some of brevianamide F’ putative analogs and a BGC in the BRA006 genome, initially annotated with other function, that could represent Brevianamide F biosynthetic pathway in BRA006. Therefore, the approach presented in the present work allowed us to extend the characterization of the potential of bioactive natural products produced by BRA006.

## 4. Materials and Methods

### 4.1 Collection, DNA Extraction and Sequencing

The BRA006, from MicroMarin collection (https://www.labbmar.ufc.br/micromarinbr) was cultured in A1 medium [Starch (10 g/L); Yeast extract (4 g/L); Peptone (2 g/L); Sea Water 75%] in a volume of 100 mL in 250 mL Erlenmeyer flasks. The cultures were centrifuged at 12.000 x g for 10 min and the cell pellets were resuspended in 10 uL of lysozyme 0.05 g/mL for 500 ul of SET buffer. The mixture was incubated at 37°C for 30 minutes. Then 14 ul Proteinase K (20 mg/ml) and 60 uL SDS 10% were added to the cell lysate and incubated for a further 1h at 55°C. Then 200 uL NaCl 5M was added and the temperature was raised to 37°C. 500uL of chloroform was added and the system was centrifuged at 4,500 x g for 10min at 20°C. 500uL was collected in a new tube and 300uL of isopropanol was added and incubated overnight (16 hours). The system was centrifuged at 14,000 x g for 10 minutes at 4°C. The supernatant was discarded and the DNA pellet was resuspended in TE buffer. DNA libraries were prepared using the Oxford Nanopore Ligation Sequencing Kit (SQK-LSK110) and library loading and sequencing were performed according to the manufacturer’s instructions and protocol for Flongle.

### 4.2 Genome Sequencing and Assembly Pipeline

For MinION data in use an in house pipeline with tools suggested by Oxford Nanopore. The acquisition of the raw data, a series of processing steps were followed until these genomes were assembled. The first stage of data processing consisted of converting the raw signals into DNA base sequences (base calling) using the Guppy tool (67). With this data in hand, the quality of the readings taken in the previous step was assessed using the NanoStat software (68). With these results, the processing continued with the removal of adapters from the base called reads to prepare them for subsequent steps; this was done using the Porechop tool (https://github.com/rrwick/Porechop), which also compressed the resulting reads. Once the adapters had been removed, a new quality analysis was carried out using the NanoStat tool. Once this step was completed, the low-quality bases and short reads were trimmed using the Chopper tool (69), which not only trimmed but also compressed the resulting reads. The next step was to analyze the quality of the resulting sequence, again using the NanoStat tool. With the resulting data, we then used the Flye (70) tool to assemble the genomes using filtered, high-quality reads. The genome assembly was then refined using information from reads mapped using the Racon tool (71). Finally, with the final result in hand, the quality of the final genome assembly was assessed using the Quast tool (72) and BUSCO (73). For Illumina procedures, BRA006 genome samples were sequenced using MiSeq technology at Macrogen facility (Seul, South Korea), all the processing steps such as read mapping, trimming low quality reads and *de novo* genome assembly were performed using the proprietary software Geneious (Version 11) (74).

### 4.3 Phylogenetic analysis

For phylogenetic inference, we choose a comprehensive method called digital DNA:DNA hybridization (dDDH) available in the Type Genome Server (TYGS) web tool (21). We used both MinION and Illumina data as query sequences with a standard parameter set.

### 4.4 BGCs analysis and functional annotation

To reveal biosynthetic gene clusters (BGC) from the BRA006 genome, we used the antiSMASH tool (version 7.1.0) (75) with relaxed detection strictness and all extra features selection. Coding sequences (CDS) prediction of BRA006 assemblies were made using Prokka (1.14.6) (76), which is based on prodigal (77) HMM models to identify proteins by their family motifs. Finally, we combine Prokka and antiSMASH results to obtain a better resolution of protein functions within BGCs. We manually filtered antiSMASH extra features and retrieved the most similar known cluster genbank format of our BGCs of interest from MiBiG (version 3.0) (78). Thus, python scripts were used to convert CDS of GenBank files from MiBiG and antiSMASH into fasta format in which the MiBiG sequences were used to construct reference databases and antiSMASH’s as query sequences for BLASTp. With the TSV files from BLASTp, we grouped all possible matches by each query sequence and selected the one with the lowest e-value. Finally, we used BioPython to plot the comparisons with more than 70% of similarity.

### 4.5 Metabolomic analysis

The BRA006 isolates were cultivated in 100 mL of the sterile A1 culture medium, in 250 mL Erlenmeyer flasks. The liquid cultures were extracted with ethyl acetate, and the organic phase was dried under pressure and kept at 4°C. For the LC-MS/MS analyses, organic extracts were diluted in methanol at a ratio of 1.0 mg/mL (79). The LC-MS/MS analysis itself was conducted in the Acquity UPLC H-Class (Waters, Milford, MA - US) hyphenated with Impact II mass spectrometer (Bruker Daltonics, Billerica - US). The mobile phase (flow 0.3 mL.min-1) consisted of water (A) and methanol (B) in the following gradient: 0.0 - 15.0 min (5 - 20% B, curve 6); 15.0 - 30.0 min (20-95% B, curve 6); 30.0 - 33.0 min - (100 B, curve 1); 33.0 - 40.0 min (5% B, curve 1). C18 - Luna (Phenomonex® - 100 mm x 2.1 mm x 2.6 μm) and the temperature adjusted to 35°C. The parameters adjusted for the spectrometer were: end plate offset of 500V; capillary voltage of 4.5kV; nitrogen (N2) was used as gas; drying gas flow at 5.0L.min-1; drying gas temperature at 180°C; 4 bar nebulizer gas pressure; positive ESI mode. Spectra (*m/z* 30–2000) were recorded at a rate of 8 Hz. Accurate masse s were obtained using sodium formic acid [HCOO-Na+] as an internal standard.

### 4.6 GNPS2 Molecular Networking and *in silico* annotation

The .csv and .mgf files generated in MZmine3 (23) from the raw data of the metabolomic analysis were imported into the GNPS2 platform (https://gnps2.org/), where the molecular network and spectral pairing were performed using the Feature Based Molecular Network (FBMN) (80) (https://gnps2.org/workflowinput?workflowname=feature_based_molecular_networking_workflow). We used the standard parameters for FBMN. Once the network was built, the annotations were propagated using the ChemWalker (24) tool through GNPS2 interface (https://gnps2.org/workflowinput?workflowname=chemwalker_nextflow_workflow). We used the standard parameters for ChemWalker, including COCONUT (81) as the reference database and 0 for the component index to propagate information for the whole network. For the nodes that could not be annotated, the MS/MS mass spectra were analyzed using the SIRIUS tool (25). For *in silico* annotation, both ChemWalker and SIRIUS were used to annotate a structure with the best ranked candidate. The raw data set, as well as the parameters used for preprocessing in MZmine3 and access to the results with GNPS2 are available from Zenodo https://doi.org/10.5281/zenodo.10366840.

## Author Contributions

Conceptualization, G.S.A., T.C.B., and R.R.d.S.; methodology, G.S.A., T.C.B., and R.R.d.S.; software, T.C.B., A.G.F., M.W., and R.d.S.; resources, R.R.d.S., M.E.G., L.C.L., G.P., D.B.B.T., and N.P.L.; data acquisition, G.S.A., M.P., G.M.V.S., E.G.F., R.d.F., P.R.T., and F.R.O.; data curation, G.S.A., T.C.B., L.G.M., H.T., L.G.; writing—original draft preparation, G.S.A., T.C.B., and R.R.d.S.; writing—review and editing, all authors; supervision, R.R.d.S.;

## Acknowledgments

MW was supported by the UCR startup.

## Data availability

All data, Python scripts and Jupyter notebooks used during metabologenomics data analysis are available on this project GitHub page: https://github.com/computational-chemical-biology/metabologenomics. The FBMN results can be found here https://gnps2.org/status?task=7b134da60f0f4a80aec790d2a294aedd and ChemWalker results here https://gnps2.org/status?task=9141a5cdabf842d39387e514e5305398. The assemblies are available on NBCI database. MiION: https://www.ncbi.nlm.nih.gov/biosample/SAMN39609461/ eIllumina: https://www.ncbi.nlm.nih.gov/biosample/?term=SAMN29586427.

## References

1. Fenical, W.; Jensen, P.R. Developing a new resource for drug discovery: marine actinomycete bacteria. Nat Chem Biol. 2006, 12, 666–73.

2. Newman, D.J.; Cragg, G.M. Natural Products as Sources of New Drugs over the Nearly Four Decades from 01/1981 to 09/2019. J Nat Prod. 2020, 83, 770–803.

3. Kim, L.J.; Ohashi, M.; Zhang, Z.; Tan, D.; Asay, M.; Cascio, D.; Rodriguez, J.A.; Tang, Y.; Nelson, H.M. Prospecting for natural products by genome mining and microcrystal electron diffraction. Nat Chem Biol. 2021, 17, 872–877.

4. Wilke, D.V.; Jimenez, P.C.; Branco, P.C.; Rezende-Teixeira, P.; Trindade-Silva, A.E.; Bauermeister, A.; Lopes, N.P.; Costa-Lotufo, L.V. Anticancer Potential of Compounds from the Brazilian Blue Amazon. Planta Med. 2021, 87, 49–70.

5. Hifnawy, M.S.; Fouda, M.M.; Sayed, A.M.; Mohammed, R.; Hassan, H.M.; AbouZid, S.F.; Rateb, M.E.; Keller, A.; Adamek, M.; Ziemert, N.; Abdelmohsen, U.R. The genus Micromonospora as a model microorganism for bioactive natural product discovery. RSC Adv. 2020, 10, 20939–20959.

6. Sousa, T. da. S.; Jimenez, P.C.; Ferreira, E.G.; Silveira, E.R.; Braz-Filho, R.; Pessoa, O.D.; Costa-Lotufo, L.V. Anthracyclinones from Micromonospora sp. J Nat Prod. 2012, 75, 489–93.

7. Ruzzini, A.C.; Clardy, J Gene Flow and Molecular Innovation in Bacteria. Curr Biol. 2016, 26, R859–R864.

8. Domingues Vieira, B.; Niero, H.; de Felício, R.; Giolo Alves, L.F.; Freitas Bazzano, C.; Sigrist, R.; Costa Furtado, L.; Felix Persinoti, G.; Veras Costa-Lotufo, L.; Barretto Barbosa Trivella, D. Production of Epoxyketone Peptide-Based Proteasome Inhibitors by Streptomyces sp. BRA-346: Regulation and Biosynthesis. Front Microbiol. 2022, 13, 786008.

9. Sun, W.; Wu, W.; Liu, X.; Zaleta-Pinet, D.A.; Clark, B.R. Bioactive Compounds Isolated from Marine-Derived Microbes in China: 2009-2018. Mar Drugs. 2019, 17, 339.

10. Behsaz, B.; Bode, E.; Gurevich, A.; Shi, Y.N.; Grundmann, F.; Acharya, D.; Caraballo-Rodríguez, A.M.; Bouslimani, A.; Panitchpakdi, M.; Linck, A.; Guan, C.; Oh, J.; Dorrestein, P.C.; Bode, H.B.; Pevzner, P.A.; Mohimani, H. Integrating genomics and metabolomics for scalable non-ribosomal peptide discovery. Nat Commun. 2021, 12, 3225.

11. Andrade, L.S.N. Identification and characterization of biosynthetic clusters from Micromonospora sp. M.Sc dissertation, University of São Paulo, São Paulo, Brazil, 2020.

12. Kato, N.N.; Arini, G.S.; Silva, R.R.; Bichuette, M.E.; Bitencourt, J.A.P.; Lopes, N.P. The World of Cave Microbiomes: Biodiversity, Ecological Interactions, Chemistry, and the Multi-Omics Integration. J Braz Chem Soc. 2023, 00, 1–16.

13. Yan, S.; Zeng, M.; Wang, H.; Zhang, H. Micromonospora: A Prolific Source of Bioactive Secondary Metabolites with Therapeutic Potential. J Med Chem. 2022, 13, 8735–8771.

14. Lasch, C.; Gummerlich, N.; Myronovskyi, M.; Palusczak, A.; Zapp, J.; Luzhetskyy, A. Loseolamycins: A Group of New Bioactive Alkylresorcinols Produced after Heterologous Expression of a Type III PKS from Micromonospora endolithica. Molecules. 2020, 20, 4594.

15. Paderog, M.J.V.; Suarez, A.F.L.; Sabido, E.M.; Low, Z.J.; Saludes, J.P.; Dalisay, D.S. Anthracycline Shunt Metabolites From Philippine Marine Sediment-Derived Streptomyces Destroy Cell Membrane Integrity of Multidrug-Resistant Staphylococcus aureus. Front Microbiol. 2020, 11, 743.

16. Silva, L.J.; Crevelin, E.J.; Souza, D.T.; Lacerda-Júnior, G.V.; de Oliveira, V.M.; Ruiz, A.L.T.G.; Rosa, L.H.; Moraes, L.A.B.; Melo, I.S. Actinobacteria from Antarctica as a source for anticancer discovery. Sci Rep. 2020, 1, 13870.

17. Hecht, S.M. Bleomycin: new perspectives on the mechanism of action. J Nat Prod. 2000, 1, 158–68.

18. Hofstead, S.J.; Matson, J.A.; Malacko, A.R.; Marquardt, H. Kedarcidin, a new chromoprotein antitumor antibiotic. II. Isolation, purification and physico-chemical properties. J Antibiot (Tokyo), 1992, 8, 1250–4.

19. Low, Z.J.; Ma, G.L.; Tran, H.T.; Zou, Y.; Xiong, J.; Pang, L.; Nuryyeva, S.; Ye, H.; Hu, J.F.; Houk, K.N.; Liang, Z.X. Sungeidines from a Non-canonical Enediyne Biosynthetic Pathway. J Am Chem Soc. 2020, 4, 1673–1679.

20. Unno, R.; Michishita, H.; Inagaki, H.; Suzuki, Y.; Baba, Y.; Jomori, T.; Nishikawa, T.; Isobe, M. Synthesis and antitumor activity of water-soluble enediyne compounds related to dynemicin A. Bioorg Med Chem. 1997, 5, 987–99.

21. Meier-Kolthoff, J.P.; Göker, M. TYGS is an automated high-throughput platform for state-of-the-art genome-based taxonomy. Nat Commun. 2019, 1, 2182.

22. Chambers, M.C.; Maclean, B.; Burke, R.; Amodei, D.; Ruderman, D.L.; Neumann, S.; Gatto, L.; Fischer, B.; Pratt, B.; Egertson, J.; Hoff, K.; Kessner, D.; Tasman, N.; Shulman, N.; Frewen, B.; Baker, T.A.; Brusniak, M.Y.; Paulse, C.; Creasy, D.; Flashner, L.; Kani, K.; Moulding, C.; Seymour, S.L.; Nuwaysir, L.M.; Lefebvre, B.; Kuhlmann, F.; Roark, J.; Rainer, P.; Detlev, S.; Hemenway, T.; Huhmer, A.; Langridge, J.; Connolly, B.; Chadick, T.; Holly, K.; Eckels, J.; Deutsch, E.W.; Moritz, R.L.; Katz, J.E.; Agus, D.B.; MacCoss, M.; Tabb, D.L.; Mallick, P. A cross-platform toolkit for mass spectrometry and proteomics. Nat Biotechnol. 2012, 10, 918–20.

23. Schmid, R.; Heuckeroth, S.; Korf, A.; Smirnov, A.; Myers, O.; Dyrlund, T.S.; Bushuiev, R.; Murray, KJ.; Hoffmann, N.; Lu, M.; Sarvepalli, A.; Zhang, Z.; Fleischauer, M.; Dührkop, K.; Wesner, M.; Hoogstra, SJ.; Rudt, E.; Mokshyna, O.; Brungs, C.; Ponomarov, K.; Mutabdžija, L.; Damiani, T.; Pudney, CJ.; Earll, M.; Helmer, PO.; Fallon, TR.; Schulze, T.; Rivas-Ubach, A.; Bilbao, A.; Richter, H.; Nothias, LF.; Wang, M.; Orešič, M.; Weng, JK.; Böcker, S.; Jeibmann, A.; Hayen, H.; Karst, U.; Dorrestein, PC.; Petras, D.; Du, X.; Pluskal, T. Integrative analysis of multimodal mass spectrometry data in MZmine 3. Nat Biotechnol. 2023, 41, 447–449.

24. Borelli, T.C.; Arini, G.S.; Feitosa, L.G.P.; Dorrestein, P.C.; Lopes, N.P.; da Silva, R.R. Improving annotation propagation on molecular networks through random walks: introducing ChemWalker. Bioinformatics. 2023, 3, btad078.

25. Dührkop, K.; Fleischauer, M.; Ludwig, M.; Aksenov, A.A.; Melnik, A.V.; Meusel, M.; Dorrestein, P.C.; Rousu, J.; Böcker, S. SIRIUS 4: a rapid tool for turning tandem mass spectra into metabolite structure information. Nat Methods. 2019, 4, 299–302.

26. Djoumbou Feunang, Y.; Eisner, R.; Knox, C.; Chepelev, L.; Hastings, J.; Owen, G.; Fahy, E.; Steinbeck, C.; Subramanian, S.; Bolton, E.; Greiner, R.; Wishart DS. ClassyFire: automated chemical classification with a comprehensive, computable taxonomy. J Cheminform. 2016, 8, 61.

27. de Valk, K.S.; Deken, M.M.; Handgraaf, H.J.M.; Bhairosingh, S.S.; Bijlstra, O.D.; van Esdonk, M.J.; Terwisscha van Scheltinga, A.G.T.; Valentijn, A.R.P.M.; March, T.L.; Vuijk, J.; Peeters, K.C.M.J.; Holman, F.A.; Hilling, D.E.; Mieog, J.S.D.; Frangioni, J.V.; Burggraaf, J.; Vahrmeijer, A.L. First-in-Human Assessment of cRGD-ZW800-1, a Zwitterionic, Integrin-Targeted, Near-Infrared Fluorescent Peptide in Colon Carcinoma. Clin Cancer Res. 2020, 15, 3990–3998.

28. Paulsen, Ø.; Aass, N.; Kaasa, S.; Dale, O. Do corticosteroids provide analgesic effects in cancer patients? A systematic literature review. J Pain Symptom Manage. 2013, 1, 96–105.

29. Park, H.B.; Crawford, J.M. Pyrazinone protease inhibitor metabolites from Photorhabdus luminescens. J Antibiot (Tokyo). 2016, 8, 616–21.

30. Zhu, T.; Chen, Z.; Liu, P.; Wang, Y.; Xin, Z.; Zhu, W. New rubrolides from the marine-derived fungus Aspergillus terreus OUCMDZ-1925. J Antibiot. 2014, 67, 315–318.

31. Costa, M.; Sampaio-Dias, IE.; Castelo-Branco, R.; Scharfenstein, H.; Rezende de Castro, R.; Silva, A.; Schneider, M.P.C.; Araújo, M.J.; Martins, R.; Domingues, V.F.; Nogueira, F.; Camões, V.; Vasconcelos, V.M.; Leão, P.N. Structure of Hierridin C, Synthesis of Hierridins B and C, and Evidence for Prevalent Alkylresorcinol Biosynthesis in Picocyanobacteria. J Nat Prod. 2019, 2, 393–402.

32. Shishido, T.; Hachisuka, M.; Ryuzaki, K.; Miura, Y.; Tanabe, A.; Tamura, Y.; Kusayanagi, T.; Takeuchi, T.; Kamisuki, S.; Sugawara, F.; Sahara, H. EpsinR, a target for pyrenocine B, role in endogenous MHC-II-restricted antigen presentation. Eur J Immunol. 2014, 11, 3220–31.

33. Liu, C.H.; Chen, C.Y.; Huang, A.M.; Li, J.H. Subamolide A, a component isolated from Cinnamomum subavenium, induces apoptosis mediated by mitochondria-dependent, p53 and ERK1/2 pathways in human urothelial carcinoma cell line NTUB1. J Ethnopharmacol. 2011, 1, 503–11.

34. Cheng, M.J.; Lee, S.J.; Chang, Y.Y.; Wu, S.H.; Tsai, I.L.; Jayaprakasam, B.; Chen, I.S. Chemical and cytotoxic constituents from Peperomia sui. Phytochemistry. 2003, 5, 603–8.

35. Zang, Y.; Genta-Jouve, G.; Sun, T.A.; Li, X.; Didier, B.; Mann, S.; Mouray, E.; Larsen, A.K.; Escargueil, A.E.; Nay, B.; Prado, S. Unexpected talaroenamine derivatives and an undescribed polyester from the fungus Talaromyces stipitatus ATCC10500. Phytochemistry. 2015, 119, 70–5.

36. Brkljača, R.; Urban, S. HPLC-NMR, and HPLC-MS investigation of antimicrobial constituents in Cystophora monilifera and Cystophora subfarcinata. Phytochemistry. 2015, 117, 200–208.

37. Cordero, A.M.F.; Gonzales III, A.A. Using multiscale molecular modeling to analyze possible NS2b-NS3 protease inhibitors from medicinal plants endemic to the Philippines. bioRxiv. 2023.

38. Wang, W.; Hong, J.; Lee, C.O.; Im, K.S.; Choi, J.S.; Jung, J.H. Cytotoxic sterols and saponins from the starfish Certonardoa semiregularis. J Nat Prod. 2004, 4, 584–91.

39. Kalauni, S.K.; Choudhary, M.I.; Shaheen, F.; Manandhar, M.D.; Atta-ur-Rahman.; Gewali, M.B.; Khalid, A. Steroidal alkaloids from the leaves of Sarcococca coriacea of Nepalese origin. J Nat Prod. 2001, 6, 842–4.

40. Salen, G.; Verga, D.; Batta, A.K.; Tint, G.S.; Shefer, S. Effect of 7-ketolithocholic acid on bile acid metabolism in humans. Gastroenterology. 1982, 2, 341–7.

41. Mandia, D.; Chaussenot, A.; Besson, G.; Lamari, F.; Castelnovo, G.; Curot, J.; Duval, F.; Giral, P.; Lecerf, J.M.; Roland, D.; Pierdet, H.; Douillard, C.; Nadjar, Y. Cholic acid as a treatment for cerebrotendinous xanthomatosis in adults. J Neurol. 2019, 266, 2043–2050.

42. Bedir, E.; Karakoyun, Ç.; Doğan, G.; Kuru, G.; Küçüksolak, M.; Yusufoğlu, H. New Cardenolides from Biotransformation of Gitoxigenin by the Endophytic Fungus Alternaria eureka 1E1BL1: Characterization and Cytotoxic Activities. Molecules. 2021, 10, 3030.

43. Che, Y.; Swenson, D.C.; Gloer, J.B.; Koster, B.; Malloch, D. Pseudodestruxins A and B: new cyclic depsipeptides from the coprophilous fungus Nigrosabulum globosum. J Nat Prod. 2001, 5, 555–8.

44. Kanehisa, M.; Goto, S. KEGG: kyoto encyclopedia of genes and genomes. Nucleic Acids Res. 2000, 1, 27–30.

45. Wang, M.; Carver, J.J.; Phelan, V.V.; Sanchez, L.M.; Garg, N.; Peng, Y.; Nguyen, D.D.; Watrous, J.; Kapono, C.A.; Luzzatto-Knaan, T.; Porto, C.; Bouslimani, A.; Melnik, A.V.; Meehan, M.J.; Liu, W.T.; Crüsemann, M.; Boudreau, P.D.; Esquenazi, E.; Sandoval-Calderón, M.; Kersten, R.D.; Pace, L.A.; Quinn, R.A.; Duncan, K.R.; Hsu, C.C.; Floros, D.J.; Gavilan, R.G.; Kleigrewe, K.; Northen, T.; Dutton, R.J.; Parrot, D.; Carlson, E.E.; Aigle, B.; Michelsen, C.F.; Jelsbak, L.; Sohlenkamp, C.; Pevzner, P.; Edlund, A.; McLean, J.; Piel, J.; Murphy, B.T.; Gerwick, L.; Liaw, C.C.; Yang, Y.L.; Humpf, H.U.; Maansson, M.; Keyzers, R.A.; Sims, A.C.; Johnson, A.R.; Sidebottom, A.M.; Sedio, B.E.; Klitgaard, A.; Larson, C.B. P. CAB.; Torres-Mendoza, D.; Gonzalez, D.J.; Silva, D.B.; Marques, L.M.; Demarque, D.P.; Pociute, E.; O’Neill, E.C.; Briand, E.; Helfrich, E.J.N.; Granatosky, E.A.; Glukhov, E.; Ryffel, F.; Houson, H.; Mohimani, H.; Kharbush, J.J.; Zeng, Y.; Vorholt, J.A.; Kurita, K.L.; Charusanti, P.; McPhail, K.L.; Nielsen, K.F.; Vuong, L.; Elfeki, M.; Traxler, M.F.; Engene, N.; Koyama, N.; Vining, O.B.; Baric, R.; Silva, R.R.; Mascuch, S.J.; Tomasi, S.; Jenkins, S.; Macherla, V.; Hoffman, T.; Agarwal, V.; Williams, P.G.; Dai, J.; Neupane, R.; Gurr, J.; Rodríguez, A.M.C.; Lamsa, A.; Zhang, C.; Dorrestein, K.; Duggan, B.M.; Almaliti, J.; Allard, P.M.; Phapale, P.; Nothias, L.F.; Alexandrov, T.; Litaudon, M.; Wolfender, J.L.; Kyle, J.E.; Metz, T.O.; Peryea, T.; Nguyen, D.T.; VanLeer, D.; Shinn, P.; Jadhav, A.; Müller, R.; Waters, K.M.; Shi, W.; Liu, X.; Zhang, L.; Knight, R.; Jensen, P.R.; Palsson, B.O.; Pogliano, K.; Linington, R.G.; Gutiérrez, M.; Lopes, N.P.; Gerwick, W.H.; Moore, B.S.; Dorrestein, P.C.; Bandeira, N. Sharing and community curation of mass spectrometry data with Global Natural Products Social Molecular Networking. Nat Biotechnol. 2016, 34, 828–837.

46. Silva, A.E.T.; Guimarães, L.A.; Ferreira, E.G.; Torres, M.da.C.M.; da Silva, A.B.; Branco, P.C.; Oliveira, F.A.S.; Silva, G.G.Z.; Wilke, D.V.; Silveira, E.R.; Pessoa, O.D.L.; Jimenez, P.C.; Costa-Lotufo, L.V. Bioprospecting Anticancer Compounds from the Marine-Derived Actinobacteria Actinomadura sp. Collected at the Saint Peter and Saint Paul Archipelago (Brazil). J Braz Chem Soc. 2017, 28, 465 – 474.

47. Wick, R.R.; Judd, L.M.; Gorrie, C.L.; Holt, K.E. Completing bacterial genome assemblies with multiplex MinION sequencing. Microb Genom. 2017, 10, e000132.

48. Laver, T.; Harrison, J.; O’Neill, P.A.; Moore, K.; Farbos, A.; Paszkiewicz, K.; Studholme, D.J. Assessing the performance of the Oxford Nanopore Technologies MinION. Biomol Detect Quantif. 2015, 3, 1–8.

49. Medema, M.H.; Cimermancic, P.; Sali, A.; Takano, E.; Fischbach, M.A. A systematic computational analysis of biosynthetic gene cluster evolution: lessons for engineering biosynthesis. PLoS Comput Biol. 2014, 12, e1004016.

50. Xu, M.; Wright, G.D. Heterologous expression-facilitated natural products’ discovery in actinomycetes. J Ind Microbiol Biotechnol. 2019, 3–4, 415-431.

51. Yamanaka, K.; Reynolds, K.A.; Kersten, R.D.; Ryan, K.S.; Gonzalez, D.J.; Nizet, V.; Dorrestein, P.C.; Moore, B.S. Direct cloning and refactoring of a silent lipopeptide biosynthetic gene cluster yields the antibiotic taromycin A. Proc Natl Acad Sci U S A. 2014, 5, 1957–62.

52. Bertin, M.J.; Roduit, A.F.; Sun, J.; Alves, G.E.; Via, C.W.; Gonzalez, M.A.; Zimba, P.V.; Moeller, P.D.R. Tricholides A and B and Unnarmicin D: New Hybrid PKS-NRPS Macrocycles Isolated from an Environmental Collection of Trichodesmium thiebautii. Mar Drugs. 2017, 7, 206.

53. Fujimoto, H.; Nagano, J.; Yamaguchi, K.; Yamazaki, M. Immunosuppressive components from an Ascomycete, Diplogelasinospora grovesii. Chem Pharm Bull (Tokyo). 1998, 3, 423–9.

54. Ben Ameur Mehdi, R.; Shaaban, K.A.; Rebai, I.K.; Smaoui, S.; Bejar, S.; Mellouli, L. Five naturally bioactive molecules including two rhamnopyranoside derivatives isolated from the Streptomyces sp. strain TN58. Nat Prod Res. 2009, 12, 1095–107.

55. Belmokhtar, C.A.; Hillion, J.; Ségal-Bendirdjian, E. Staurosporine induces apoptosis through both caspase-dependent and caspase-independent mechanisms. Oncogene. 2001, 26, 3354–62.

56. Ausloos, P.; Clifton, C.L.; Lias, S.G.; Mikaya, A.I.; Stein, S.E.; Tchekhovskoi, D.V.; Sparkman, O.D.; Zaikin, V.; Zhu, D. The critical evaluation of a comprehensive mass spectral library. J Am Soc Mass Spectrom. 1999, 4, 287–99.

57. Mehetre, G.T.J.S.V.; Burkul, B.B.; Desai, D.B.S.; Dharne, M.S.; Dastager, S.G. Bioactivities and molecular networking-based elucidation of metabolites of potent actinobacterial strains isolated from the Unkeshwar geothermal springs in India. RSC Adv. 2019, 17, 9850–9859.

58. Wang, J.-T.; Shi, T.-T.; Ding, L.; Xie, J.; Zhao, P.-J. Multifunctional Enzymes in Microbial Secondary Metabolic Processes. Catalysts. 2023, 13, 581.

59. Maiya, S.; Grundmann, A.; Li, S.M.; Turner, G. The fumitremorgin gene cluster of Aspergillus fumigatus: identification of a gene encoding brevianamide F synthetase. Chembiochem. 2006, 7, 1062–9.

60. Mistry, J.; Chuguransky, S.; Williams, L.; Qureshi, M.; Salazar, G.A.; Sonnhammer, E.L.L.; Tosatto, S.C.E.; Paladin, L.; Raj, S.; Richardson, L.J.; Finn, R.D.; Bateman, A. Pfam: the protein families database in 2021. Nucleic Acids Res. 2021, 49, D412–D419.

61. Haft, D.H.; Selengut, J.D.; Richter, R.A.; Harkins, D.; Basu, M.K.; Beck, E. TIGRFAMs and genome properties in 2013. Nucleic Acids Res. 2013, 41, D387–D395.

62. Letunic, I.; Khedkar, S.; Bork, P. SMART: recent updates, new developments and status in 2020. Nucleic Acids Res. 2021, 49, D458–D460.

63. van Heel, A.J.; de Jong, A.; Song, C.; Viel, J.H.; Kok, J.; Kuipers, O.P. BAGEL4: a user-friendly web server to thoroughly mine RiPPs and bacteriocins. Nucleic Acids Res. 2018, 46, W278–W281.

64. Yadav, G.; Gokhale, R.S.; Mohanty, D. Towards prediction of metabolic products of polyketide synthases: an In silico analysis. PLOS Comput. Biol. 2009, 5, e1000351.

65. Biermann, F.; Wenski, S. L.; Helfrich, E. J. N. Navigating and expanding the roadmap of natural product genome mining tools. Beilstein J Org Chem. 2022, 18, 1656–1671.

66. Hannigan, G.D.; Prihoda, D.; Palicka, A.; Soukup, J.; Klempir, O.; Rampula, L.; Durcak, J.; Wurst, M.; Kotowski, J.; Chang, D.; Wang, R.; Piizzi, G.; Temesi, G.; Hazuda, D.J.; Woelk, C.H.; Bitton, D.A. A deep learning genome-mining strategy for biosynthetic gene cluster prediction. Nucleic Acids Res. 2019, 18, e110.

67. Wick, R.R.; Judd, L.M.; Holt, K.E. Performance of neural network basecalling tools for Oxford Nanopore sequencing. Genome Biol. 2019, 1, 129.

68. De Coster, W.; D’Hert, S.; Schultz, D.T.; Cruts, M.; Van Broeckhoven, C. NanoPack: visualizing and processing long-read sequencing data. Bioinformatics. 2018, 15, 2666–2669.

69. De Coster, W.; Rademakers, R. NanoPack2: population-scale evaluation of long-read sequencing data. Bioinformatics. 2023, 5, btad311.

70. Kolmogorov, M.; Yuan, J.; Lin, Y.; Pevzner, P.A. Assembly of long, error-prone reads using repeat graphs. Nat Biotechnol. 2019, 5, 540–546.

71. Vaser, R.; Sović, I.; Nagarajan, N.; Šikić, M. Fast and accurate de novo genome assembly from long uncorrected reads. Genome Res. 2017, 5, 737–746.

72. Gurevich, A.; Saveliev, V.; Vyahhi, N.; Tesler, G. QUAST: quality assessment tool for genome assemblies. Bioinformatics. 2013, 8, 1072–5.

73. Manni, M.; Berkeley, M. R.; Seppey, M.; Zdobnov, E. M. BUSCO: Assessing genomic data quality and beyond. Curr Protoc. 2021, 12, e323

74. Kearse, M.; Moir, R.; Wilson, A.; Stones-Havas, S.; Cheung, M.; Sturrock, S.; Buxton, S.; Cooper, A.; Markowitz, S.; Duran, C.; Thierer, T.; Ashton, B.; Meintjes, P.; Drummond, A. Geneious Basic: an integrated and extendable desktop software platform for the organization and analysis of sequence data. Bioinformatics. 2012, 12, 1647–9.

75. Blin, K.; Shaw, S.; Augustijn, H.E.; Reitz, Z.L.; Biermann, F.; Alanjary, M.; Fetter, A.; Terlouw, B.R.; Metcalf, W.W.; Helfrich, E.J.N.; van Wezel, G.P.; Medema, M.H.; Weber, T. antiSMASH 7.0: new and improved predictions for detection, regulation, chemical structures, and visualization. Nucleic Acids Res. 2023, 51, W46–W50.

76. Seemann, T. Prokka: rapid prokaryotic genome annotation. Bioinformatics. 2014, 14, 2068–9.

77. Hyatt, D.; Chen, G.L.; LoCascio, P.F.; Land, M.L.; Larimer, F.W.; Hauser, L.J. Prodigal: prokaryotic gene recognition and translation initiation site identification. BMC Bioinformatics. 2010, 1, 119.

78. Terlouw, B.R.; Blin, K.; Navarro-Muñoz, J.C.; Avalon, N.E.; Chevrette, M.G.; Egbert, S.; Lee, S.; Meijer, D.; Recchia, M.J.J.; Reitz, Z.L.; van Santen, J.A.; Selem-Mojica, N.; Tørring, T.; Zaroubi, L.; Alanjary, M.; Aleti, G.; Aguilar, C.; Al-Salihi, S.A.A.; Augustijn, H.E.; Avelar-Rivas, J.A.; Avitia-Domínguez, LA.; Barona-Gómez, F.; Bernaldo-Agüero, J.; Bielinski, V.A.; Biermann, F.; Booth, T.J.; Carrion Bravo, V.J.; Castelo-Branco, R.; Chagas, F.O.; Cruz-Morales, P.; Du, C.; Duncan, K.R.; Gavriilidou, A.; Gayrard, D.; Gutiérrez-García, K.; Haslinger, K.; Helfrich, E.J.N.; van der Hooft, J.J.J.; Jati, A.P.; Kalkreuter, E.; Kalyvas, N.; Kang, K.B.; Kautsar, S.; Kim, W.; Kunjapur, A.M.; Li, Y.X.; Lin, G.M.; Loureiro, C.; Louwen, J.J.R.; Louwen, N.L.L.; Lund, G.; Parra, J.; Philmus, B.; Pourmohsenin, B.; Pronk, L.J.U.; Rego, A.; Rex, D.A.B.; Robinson, S.; Rosas-Becerra, L.R.; Roxborough, E.T.; Schorn, M.A.; Scobie, D.J.; Singh, K.S.; Sokolova, N.; Tang, X.; Udwary, D.; Vigneshwari, A.; Vind, K.; Vromans, S.P.J.M.; Waschulin, V.; Williams, S.E.; Winter, J.M.; Witte, T.E.; Xie, H.; Yang, D.; Yu, J.; Zdouc, M.; Zhong, Z.; Collemare, J.; Linington, R.G.; Weber, T.; Medema, M.H. MIBiG 3.0: a community-driven effort to annotate experimentally validated biosynthetic gene clusters. Nucleic Acids Res. 2023, 51, D603–D610.

79. Bauermeister, A.; Zucchi, T.D.; Moraes, L.A. Mass spectrometric approaches for the identification of anthracycline analogs produced by actinobacteria. J Mass Spectrom. 2016, 6, 437–45.

80. Nothias, L.F.; Petras, D.; Schmid, R.; Dührkop, K.; Rainer, J.; Sarvepalli, A.; Protsyuk, I.; Ernst, M.; Tsugawa, H.; Fleischauer, M.; Aicheler, F.; Aksenov, A.A.; Alka, O.; Allard, P.M.; Barsch, A.; Cachet, X.; Caraballo-Rodriguez, A.M.; Da Silva, R.R.; Dang, T.; Garg, N.; Gauglitz, J.M.; Gurevich, A.; Isaac, G.; Jarmusch, A.K.; Kameník, Z.; Kang, K.B.; Kessler, N.; Koester, I.; Korf, A.; Le Gouellec, A.; Ludwig, M.; Martin, H C.; McCall, L.I.; McSayles, J.; Meyer, S.W.; Mohimani, H.; Morsy, M.; Moyne, O.; Neumann, S.; Neuweger, H.; Nguyen, N.H.; Nothias-Esposito, M.; Paolini, J.; Phelan, V.V.; Pluskal, T.; Quinn, R.A.; Rogers, S.; Shrestha, B.; Tripathi, A.; van der Hooft, J.J.J.; Vargas, F.; Weldon, K.C.; Witting, M.; Yang, H.; Zhang, Z.; Zubeil, F.; Kohlbacher, O.; Böcker, S.; Alexandrov, T.; Bandeira, N.; Wang, M.; Dorrestein, P.C. Feature-based molecular networking in the GNPS analysis environment. Nat Methods. 2020, 9, 905–908.

81. Sorokina, M.; Merseburger, P.; Rajan, K.; Yirik, M.A.; Steinbeck, C. COCONUT online: Collection of Open Natural Products database. J Cheminform. 2021, 1, 2.

