## Supplementary material for "Characterization of a marine bacteria through a novel metabologenomics approach": Supp_figures.docx

^g^Laboratory of Bioproducts, Department of Microbiology. Institute of Biomedical Sciences, University of São Paulo, São Paulo, 05508-900, Brazil

^h^Centro de Metabolómica y Bioanálisis (CEMBIO), Facultad de Farmacia, Universidad San Pablo-CEU, CEU Universities, Urbanización Montepríncipe, Boadilla del Monte, Spain

^i^Departamento de Tecnologías de la Información, Escuela Politécnica Superior, Universidad San Pablo-CEU, CEU Universities, Urbanización Montepríncipe, Boadilla del Monte, Spain

^j^Department of Computer Science and Engineering, University of California Riverside, Riverside, CA 92521, USA

^*^G.S.A. and T.C.B. contributed equally to this work.

^#^Author to whom correspondence should be addressed.

| 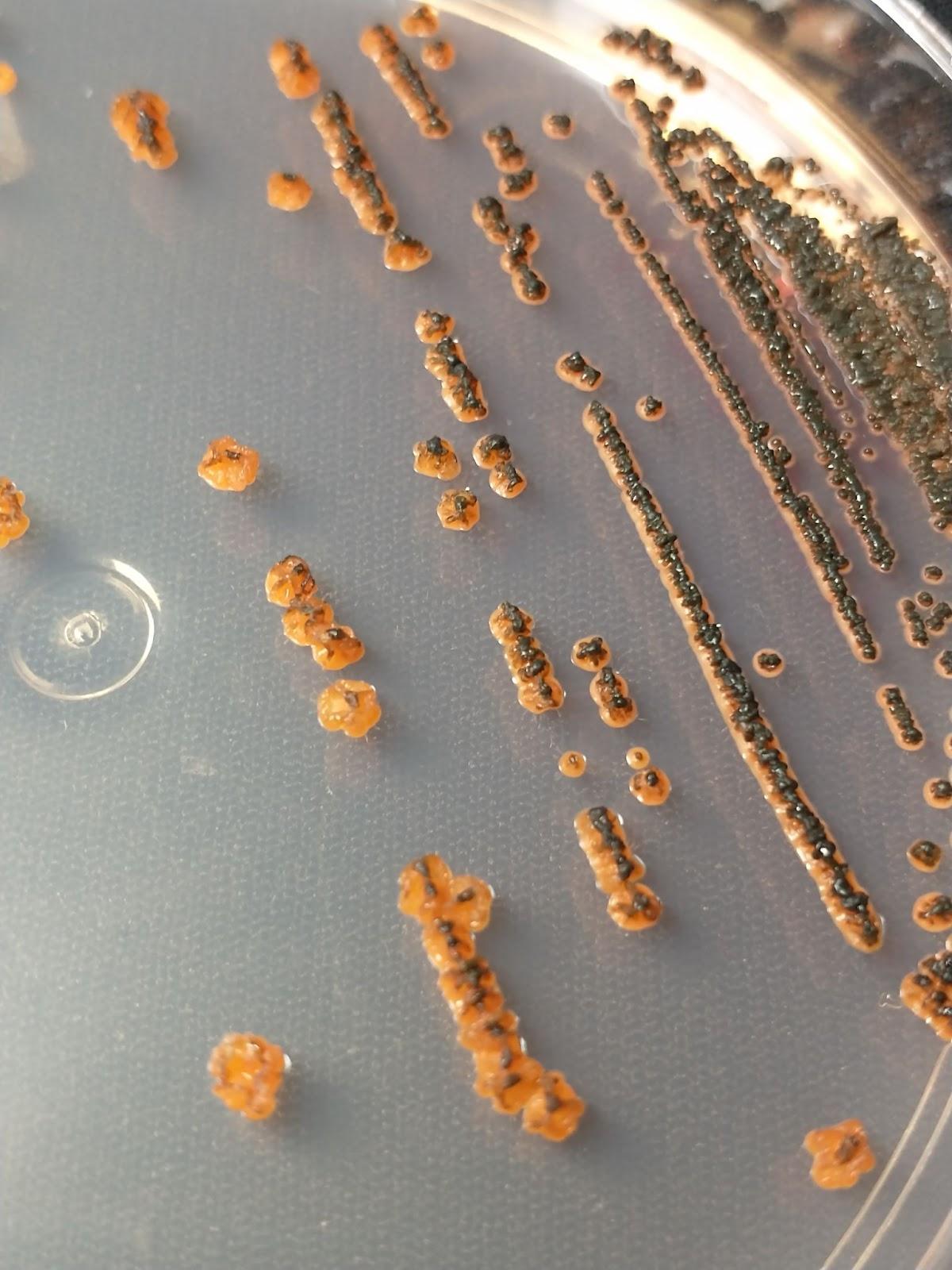 | 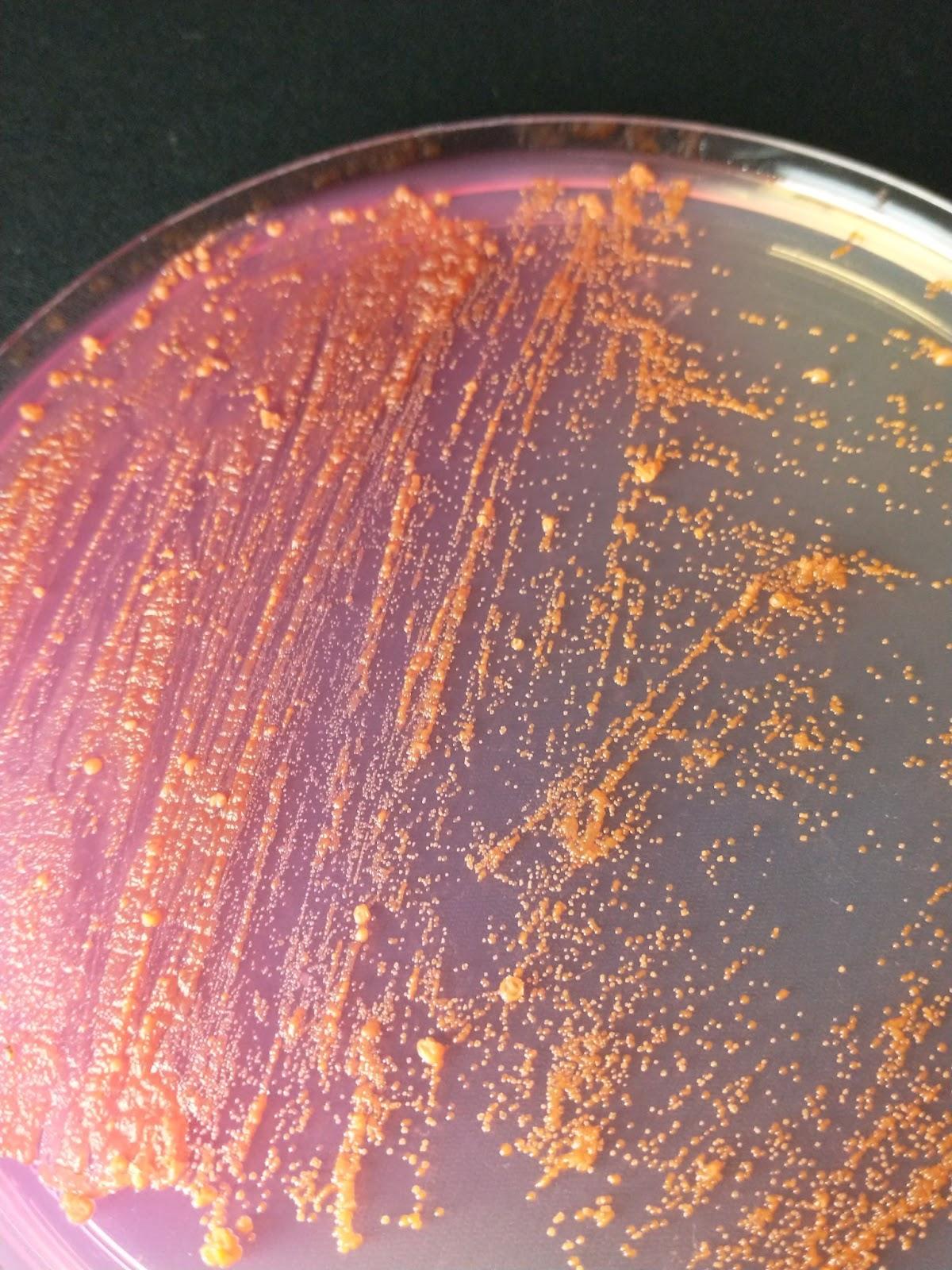 |
| --- | --- |

Supplementary Figure 1. Macroscopic aspects of BRA006 colonies. On the left, the colonies are highlighted in orange with their apical growth and their respective spores in black. On the right, the color of the water is highlighted by the growth of this bacterium.

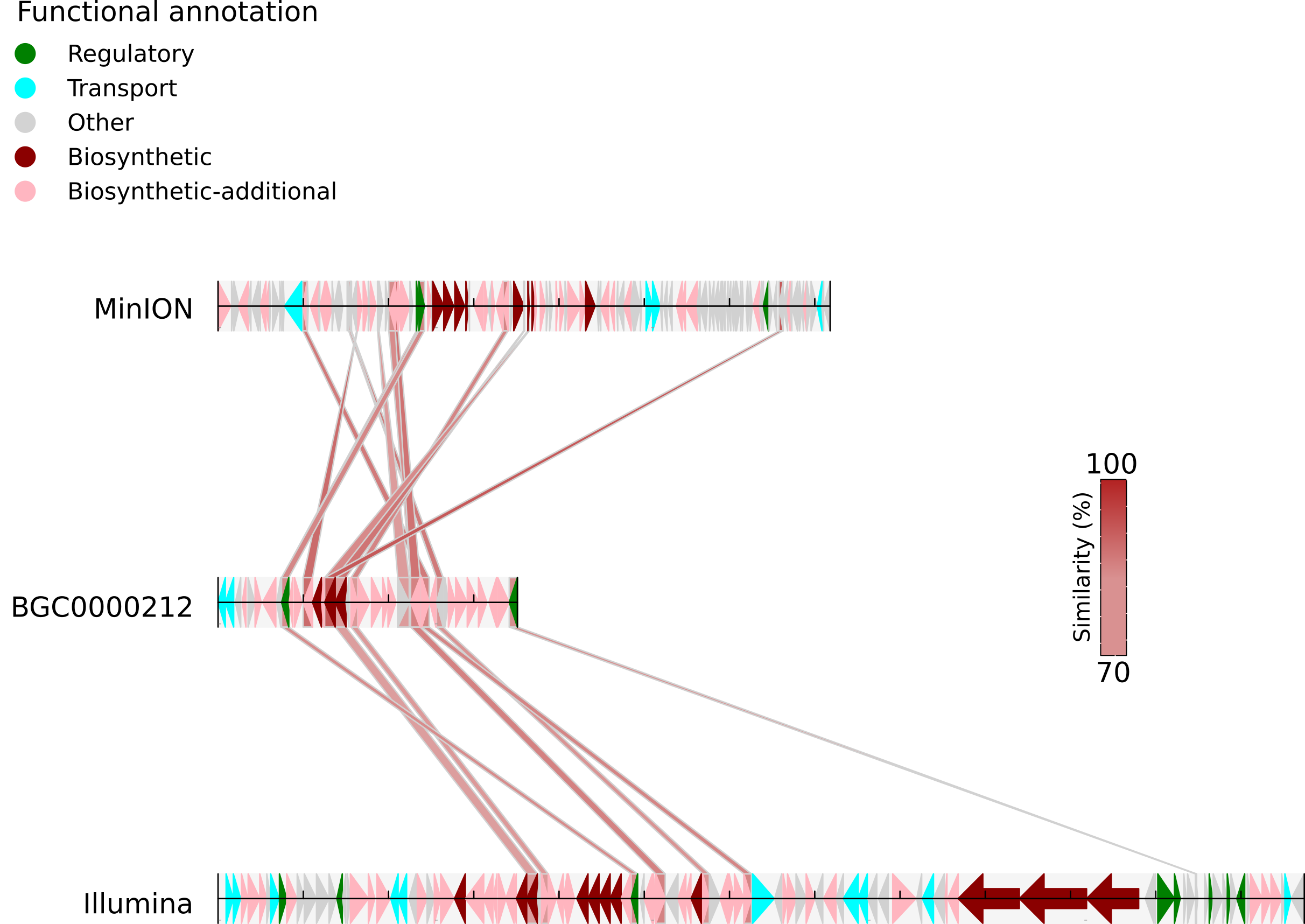

Supplementary Figure 2. Cenerubin B’s BGC and its similarities between MinION and Illumina sequencing methods against its reference BGC BGC0000212.

Supplementary Figure 3. Quinolidomicin A’s BGC and its similarity similarities between MinION and Illumina sequencing methods against reference BGC0002520 original from Micromonospora sp .
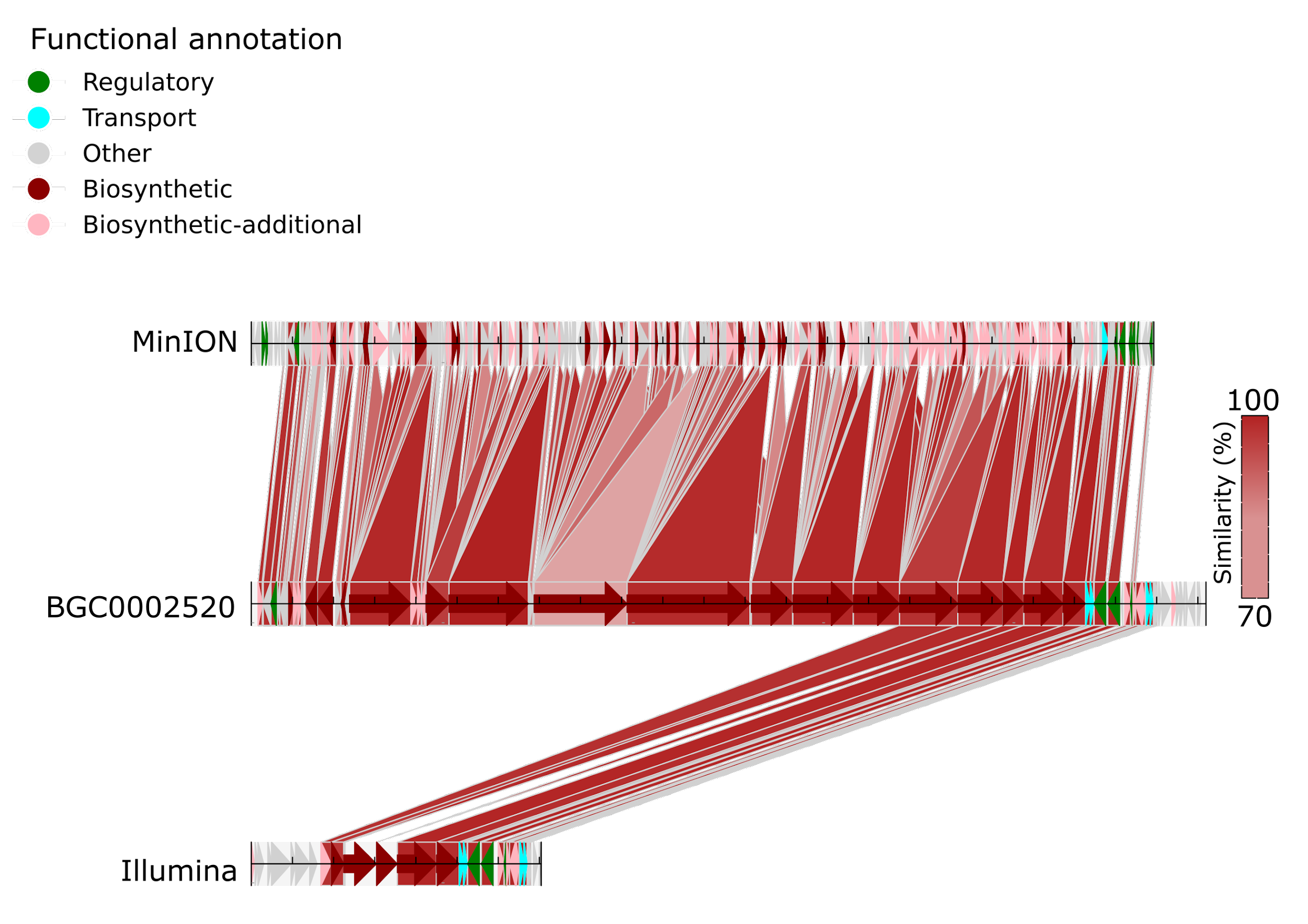

Supplementary Figure 4. Staurosporin biosynthetic pathway with its respective annotated enzymes, highlighted in red boxes. Highlighted in the green box, brevianamide F.
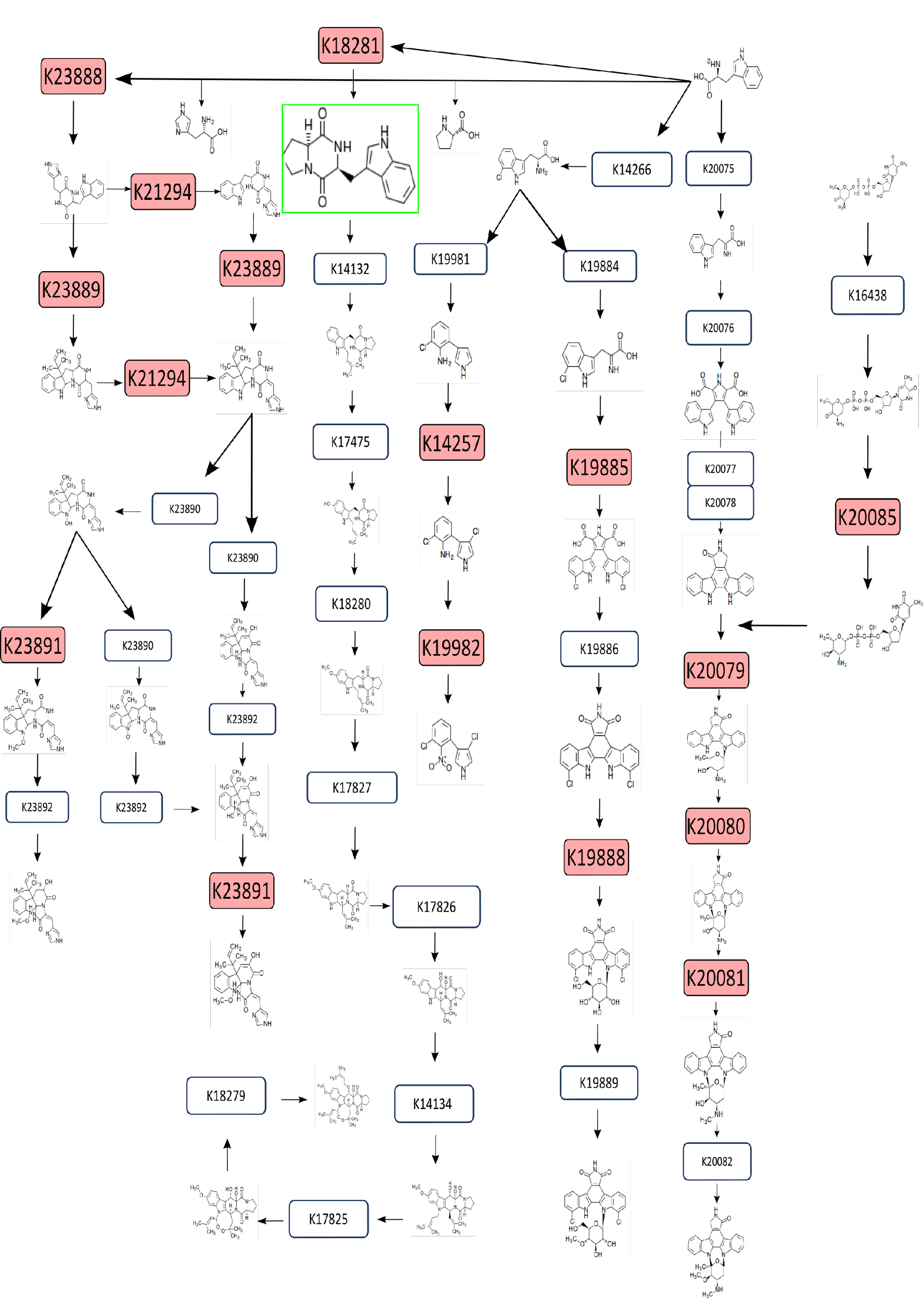
