## Supplementary material for "Characterization of a marine bacteria through a novel metabologenomics approach": SuppTable4-annotated_compounds.docx

Supplementary Table 1. Relationship between retention time (RT), m/z ratio for precursor ions (precursor mass - PM), annotation method, InChIKey, kingdom, superclass and chemical class for the annotated compounds.

| **PM (m/z)** | **RT (min)** | **Annotated Method** | **InChIKey** | **Kingdom** | **Superclass** | **Class** |
| --- | --- | --- | --- | --- | --- | --- |
| 221,0921 | 3,46 | ChemWalker | BVKMJNLLWFFTEI-UHFFFAOYSA-N | Organic compounds | Organoheterocyclic compounds | Azolidines |
| 388,2122 | 4,28 | ChemWalker | HRHBRMVWRHJCLW-UHFFFAOYSA-N | Organic compounds | Phenylpropanoids and polyketides | Coumarins and derivatives |
| 287,1282 | 4,27 | ChemWalker | UJUNVNYVYMHQIS-UHFFFAOYSA-N | Organic compounds | Benzenoids | Benzene and substituted derivatives |
| 203,1755 | 4,28 | ChemWalker | VZDWHAQZFONNQU-UHFFFAOYSA-N | Organic compounds | Organoheterocyclic compounds | Tetrahydrofurans |
| 235,1080 | 4,57 | ChemWalker | MFUNIDMQFPXVGU-UHFFFAOYSA-N | Organic compounds | Organic acids and derivatives | Carboxylic acids and derivatives |
| 203,1755 | 4,59 | ChemWalker | VZDWHAQZFONNQU-UHFFFAOYSA-N | Organic compounds | Organoheterocyclic compounds | Tetrahydrofurans |
| 284,0920 | 5,49 | ChemWalker | NKZNKBLXNLBATF-UHFFFAOYSA-N | Organic compounds | Organoheterocyclic compounds | Quinolines and derivatives |
| 285,1347 | 5,64 | ChemWalker | SHQHOHRUGBYJBS-UHFFFAOYSA-N | Organic compounds | Phenylpropanoids and polyketides | Macrolides and analogues |
| 277,1186 | 6,25 | GNPS2 | GENSLUDVKWKQMX-UHFFFAOYSA-N | Organic compounds | Organic acids and derivatives | Carboxylic acids and derivatives |
| 235,1080 | 7,04 | ChemWalker | NZWOEKJFEMNONQ-UHFFFAOYSA-N | Organic compounds | Organic acids and derivatives | Carboxylic acids and derivatives |
| 293,1266 | 8,28 | ChemWalker | ZVJCAZYCLGOEGI-UHFFFAOYSA-N | Organic compounds | Benzenoids | Benzene and substituted derivatives |
| 290,1184 | 8,28 | ChemWalker | GLMXCYNQAMJMGO-UHFFFAOYSA-N | Organic compounds | Organoheterocyclic compounds | Quinolines and derivatives |
| 298,1045 | 8,28 | ChemWalker | MOHAAJWDZWJRHY-UHFFFAOYSA-N | Organic compounds | Organoheterocyclic compounds | Azoles |
| 271,1447 | 8,29 | ChemWalker | YUUJSGRMPNOCBZ-UHFFFAOYSA-N | Organic compounds | Benzenoids | Benzene and substituted derivatives |
| 355,0975 | 8,31 | ChemWalker | NYQKSLDPYVPTRT-UHFFFAOYSA-N | Organic compounds | Organic oxygen compounds | Organooxygen compounds |
| 220,0285 | 8,32 | SIRIUS | FAVZJKGWZPIQTI-MRVPVSSYSA-N | Organic compounds | Benzenoids | Benzene and substituted derivatives |
| 361,0413 | 8,32 | ChemWalker | CBJMSBDVNIRJJW-UHFFFAOYSA-N | Organic compounds | Phenylpropanoids and polyketides | Linear 1,3-diarylpropanoids |
| 296,1361 | 8,36 | ChemWalker | DHAFSUUVLRLSFO-UHFFFAOYSA-N | Organic compounds | Organoheterocyclic compounds | Tetrahydroisoquinolines |
| 214,0900 | 8,34 | ChemWalker | KMEDDQQKAPHCLF-UHFFFAOYSA-N | Organic compounds | Organoheterocyclic compounds | Azoles |
| 205,0974 | 8,73 | ChemWalker | RAGSBUDIXSABRI-UHFFFAOYSA-N | Organic compounds | Organoheterocyclic compounds | Diazines |
| 245,1286 | 8,98 | ChemWalker | NUXHSAAEUDFPJT-UHFFFAOYSA-N | Organic compounds | Organoheterocyclic compounds | Indoles and derivatives |
| 261,1239 | 9,27 | GNPS2 | LSGOTAXPWMCUCK-UHFFFAOYSA-N | Organic compounds | Organic acids and derivatives | Carboxylic acids and derivatives |
| 244,1085 | 9,49 | ChemWalker | IFQZEERDQXQTLJ-UHFFFAOYSA-N | Organic compounds | Organic acids and derivatives | Carboxylic acids and derivatives |
| 359,1973 | 9,75 | GNPS2 | UOUFYEQUUCSLPP-UHFFFAOYSA-N | Organic compounds | Phenylpropanoids and polyketides | Stilbenes |
| 258,1164 | 9,76 | GNPS2 | RYJWSNUDETVRFF-FARCUNLSSA-N | Organic compounds | Benzenoids | Benzene and substituted derivatives |
| 227,1394 | 10,64 | ChemWalker | PAAZAXUMIQKCEE-UHFFFAOYSA-N | Organic compounds | Organoheterocyclic compounds | Piperidines |
| 453,2715 | 11,16 | ChemWalker | GFPWUVSZMLDPAF-UHFFFAOYSA-N | Organic compounds | Organic acids and derivatives | Carboxylic acids and derivatives |
| 227,1395 | 11,16 | ChemWalker | RJUBCHXDUMKQQH-UHFFFAOYSA-N | Organic compounds | Organic acids and derivatives | Carboxylic acids and derivatives |
| 258,1243 | 11,74 | ChemWalker | VDMMFAOUINDEGC-UHFFFAOYSA-N | Organic compounds | Organic acids and derivatives | Carboxylic acids and derivatives |
| 453,2714 | 11,89 | ChemWalker | GFPWUVSZMLDPAF-UHFFFAOYSA-N | Organic compounds | Organic acids and derivatives | Carboxylic acids and derivatives |
| 227,1395 | 11,90 | GNPS2 | NXDGLHJHHJJTSZ-UHFFFAOYSA-N | Organic compounds | Organic oxygen compounds | Organooxygen compounds |
| 218,1406 | 13,90 | ChemWalker | SLLDHXKXDFLVPF-UHFFFAOYSA-N | Organic compounds | Alkaloids and derivatives | None |
| 261,1240 | 13,98 | ChemWalker | MLIUTNHOSMIVTE-UHFFFAOYSA-N | Organic compounds | Organoheterocyclic compounds | Azolidines |
| 284,1401 | 14,72 | ChemWalker | YNOVWLONBDRSSK-UHFFFAOYSA-N | Organic compounds | Organoheterocyclic compounds | Indoles and derivatives |
| 283,1060 | 14,74 | ChemWalker | WOQRPBXJCRJRBC-UHFFFAOYSA-M | Organic compounds | Organic oxygen compounds | Organooxygen compounds |
| 521,2405 | 14,75 | ChemWalker | ZGIGRYSYJMUJMP-UHFFFAOYSA-N | Organic compounds | Organic acids and derivatives | Carboxylic acids and derivatives |
| 261,1240 | 14,75 | ChemWalker | PYQJYHACQOBZLF-UHFFFAOYSA-N | Organic compounds | Organic acids and derivatives | Carboxylic acids and derivatives |
| 277,1553 | 14,98 | GNPS2 | GENSLUDVKWKQMX-UHFFFAOYSA-N | Organic compounds | Organic acids and derivatives | Carboxylic acids and derivatives |
| 263,1395 | 15,96 | GNPS2 | LMDVFSHGYANGRP-UHFFFAOYSA-N | Organic compounds | Organic acids and derivatives | Carboxylic acids and derivatives |
| 211,1443 | 16,21 | GNPS2 | SZJNCZMRZAUNQT-UHFFFAOYSA-N | Organic compounds | Organic acids and derivatives | Carboxylic acids and derivatives |
| 211,1443 | 16,46 | GNPS2 | SZJNCZMRZAUNQT-UHFFFAOYSA-N | Organic compounds | Organic acids and derivatives | Carboxylic acids and derivatives |
| 211,1443 | 17,10 | GNPS2 | SZJNCZMRZAUNQT-UHFFFAOYSA-N | Organic compounds | Organic acids and derivatives | Carboxylic acids and derivatives |
| 421,2815 | 17,22 | ChemWalker | RZIIIIQWCVBQRA-UHFFFAOYSA-N | Organic compounds | Organic acids and derivatives | Carboxylic acids and derivatives |
| 292,0972 | 18,06 | ChemWalker | HKBMKYVWVUSDSV-UHFFFAOYSA-M | Organic compounds | Phenylpropanoids and polyketides | Coumarins and derivatives |
| 280,1231 | 18,44 | ChemWalker | XBSGOYCPMWEADN-UHFFFAOYSA-N | Organic compounds | Organoheterocyclic compounds | Diazanaphthalenes |
| 277,1551 | 18,54 | ChemWalker | BHSJMKOSSSPKGH-UHFFFAOYSA-N | Organic compounds | Organoheterocyclic compounds | Azolidines |
| 284,1398 | 18,89 | ChemWalker | FQLXKOCHDBPFLL-UHFFFAOYSA-N | Organic compounds | Organoheterocyclic compounds | Indoles and derivatives |
| 245,1288 | 18,93 | GNPS2 | IWIANZLCJVYEFX-UHFFFAOYSA-N | Organic compounds | Organic acids and derivatives | Carboxylic acids and derivatives |
| 489,2499 | 18,93 | ChemWalker | MMPRKMGNJFBEBT-UHFFFAOYSA-N | Organic compounds | Organic acids and derivatives | Carboxylic acids and derivatives |
| 284,1397 | 19,32 | GNPS2 | RYFZBPVMVYTEKZ-UHFFFAOYSA-N | Organic compounds | Organic acids and derivatives | Carboxylic acids and derivatives |
| 245,1289 | 19,43 | GNPS2 | QZBUWPVZSXDWSB-RYUDHWBXSA-N | Organic compounds | Organic acids and derivatives | Carboxylic acids and derivatives |
| 259,2385 | 19,82 | ChemWalker | XLTXPTGAYGVGIQ-UHFFFAOYSA-N | Organic compounds | Organic nitrogen compounds | Organonitrogen compounds |
| 208,1368 | 19,82 | SIRIUS | MXBKJMHNGFHBMV-UHFFFAOYSA-N | None | None | None |
| 222,1128 | 19,95 | GNPS2 | SDKQRNRRDYRQKY-UHFFFAOYSA-N | Organic compounds | Benzenoids | Benzene and substituted derivatives |
| 211,1235 | 20,22 | ChemWalker | YJDFIRLAWMCPDP-UHFFFAOYSA-N | Organic compounds | Alkaloids and derivatives | Harmala alkaloids |
| 311,1395 | 20,40 | ChemWalker | GXEHMSZCMANLPC-UHFFFAOYSA-N | Organic compounds | Organoheterocyclic compounds | Diazanaphthalenes |
| 294,1549 | 20,58 | ChemWalker | DKDAWRFLUBCYLM-UHFFFAOYSA-M | Organic compounds | Organic acids and derivatives | Carboxylic acids and derivatives |
| 277,1284 | 20,59 | ChemWalker | CWLWBMWELZSMPG-UHFFFAOYSA-N | Organic compounds | Organic acids and derivatives | Carboxylic acids and derivatives |
| 286,1554 | 20,81 | ChemWalker | SCVJAFKUKFRWDZ-UHFFFAOYSA-N | Organic compounds | Organoheterocyclic compounds | Azolidines |
| 213,1599 | 20,81 | ChemWalker | LPEJYDOQHAUJGY-UHFFFAOYSA-N | Organic compounds | Organic acids and derivatives | Carboxylic acids and derivatives |
| 213,1599 | 21,08 | ChemWalker | LPEJYDOQHAUJGY-UHFFFAOYSA-N | Organic compounds | Organic acids and derivatives | Carboxylic acids and derivatives |
| 241,0977 | 21,20 | ChemWalker | HKFLYRJFMVGMAU-UHFFFAOYSA-N | Organic compounds | Organoheterocyclic compounds | Azoles |
| 300,1709 | 21,53 | ChemWalker | CLPGDJVPQLMZTC-UHFFFAOYSA-N | Organic compounds | Organoheterocyclic compounds | Indoles and derivatives |
| 264,1280 | 21,65 | ChemWalker | NFXSUBLDOSKHAN-UHFFFAOYSA-N | Organic compounds | Alkaloids and derivatives | Harmala alkaloids |
| 306,1128 | 21,76 | ChemWalker | PPVQDNRCMRAFGT-UHFFFAOYSA-N | Organic compounds | Organic acids and derivatives | Carboxylic acids and derivatives |
| 227,0840 | 21,93 | GNPS2 | LTODWURWNDCSOY-UHFFFAOYSA-N | Organic compounds | Organoheterocyclic compounds | Indoles and derivatives |
| 247,1443 | 21,85 | ChemWalker | CWCMOJCXPIFZMG-UHFFFAOYSA-N | Organic compounds | Organoheterocyclic compounds | Benzazepines |
| 243,0878 | 21,96 | GNPS2 | ZJTJUVIJVLLGSP-UHFFFAOYSA-N | Organic compounds | Organoheterocyclic compounds | Pteridines and derivatives |
| 265,0696 | 21,95 | ChemWalker | CSFJGQPKIMHTFV-UHFFFAOYSA-N | Organic compounds | Organic acids and derivatives | Carboxylic acids and derivatives |
| 223,1439 | 22,00 | ChemWalker | PANQLXVBDRZFBB-UHFFFAOYSA-N | Organic compounds | Organoheterocyclic compounds | Diazines |
| 300,1710 | 21,98 | ChemWalker | MZURRFOUDOALSB-UHFFFAOYSA-N | Organic compounds | Organoheterocyclic compounds | Indoles and derivatives |
| 217,0836 | 22,01 | ChemWalker | QKPSYARWSBJEDY-UHFFFAOYSA-N | Organic compounds | Organoheterocyclic compounds | Furans |
| 306,1128 | 22,05 | ChemWalker | PPVQDNRCMRAFGT-UHFFFAOYSA-N | Organic compounds | Organic acids and derivatives | Carboxylic acids and derivatives |
| 227,1751 | 22,65 | ChemWalker | HERSSAVMHCMYSQ-UHFFFAOYSA-N | Organic compounds | Phenylpropanoids and polyketides | Macrolactams |
| 225,1595 | 22,73 | ChemWalker | IUZCDJYHMMWBBE-UHFFFAOYSA-N | Organic compounds | Organoheterocyclic compounds | Diazines |
| 259,1442 | 22,76 | ChemWalker | MVNBAVQHXXMXAO-UHFFFAOYSA-N | Organic compounds | Organic acids and derivatives | Carboxylic acids and derivatives |
| 211,0938 | 22,77 | ChemWalker | NWBBHJUOIAHCAN-UHFFFAOYSA-N | Organic compounds | Organoheterocyclic compounds | Pyrans |
| 227,1754 | 22,84 | ChemWalker | PXFBBACRARGBDY-UHFFFAOYSA-N | Organic compounds | Organoheterocyclic compounds | Oxazinanes |
| 225,1597 | 23,01 | ChemWalker | HNNUZZMCWXFRQM-UHFFFAOYSA-N | Organic compounds | Organoheterocyclic compounds | Diazines |
| 280,1229 | 23,06 | ChemWalker | XBSGOYCPMWEADN-UHFFFAOYSA-N | Organic compounds | Organoheterocyclic compounds | Diazanaphthalenes |
| 227,1753 | 23,10 | ChemWalker | HERSSAVMHCMYSQ-UHFFFAOYSA-N | Organic compounds | Phenylpropanoids and polyketides | Macrolactams |
| 261,1598 | 23,11 | GNPS2 | QPDMOMIYLJMOQJ-UHFFFAOYSA-N | Organic compounds | Organic acids and derivatives | Carboxylic acids and derivatives |
| 345,1124 | 23,11 | ChemWalker | SZSFOKCNKPUNLN-UHFFFAOYSA-N | Organic compounds | Organoheterocyclic compounds | Lactams |
| 521,3123 | 23,11 | ChemWalker | YJHCFDOTTLDFSM-UHFFFAOYSA-N | Organic compounds | Organic acids and derivatives | Carboxylic acids and derivatives |
| 245,1286 | 23,18 | ChemWalker | BMQAHUYSWMCDPJ-UHFFFAOYSA-N | Organic compounds | Organic acids and derivatives | Carboxylic acids and derivatives |
| 273,1024 | 23,23 | ChemWalker | NPYNQXKLKVFHIM-UHFFFAOYSA-M | Organic compounds | Organoheterocyclic compounds | Quinolines and derivatives |
| 257,1030 | 23,25 | ChemWalker | VKQQRZQKCNYGNQ-UHFFFAOYSA-N | Organic compounds | Organoheterocyclic compounds | Pteridines and derivatives |
| 261,1598 | 23,28 | ChemWalker | ALIBGDLGPQIBDM-UHFFFAOYSA-N | Organic compounds | Organic acids and derivatives | Carboxylic acids and derivatives |
| 262,1631 | 23,28 | ChemWalker | SHDVOJJCEZGPBX-UHFFFAOYSA-N | Organic compounds | Lipids and lipid-like molecules | Fatty Acyls |
| 209,1150 | 23,33 | ChemWalker | WQWQHCFZHDDBLY-UHFFFAOYSA-N | Organic compounds | Organoheterocyclic compounds | Pyrans |
| 396,3261 | 23,62 | ChemWalker | YIZAGIYKBZIOLR-UHFFFAOYSA-N | Organic compounds | Lipids and lipid-like molecules | Steroids and steroid derivatives |
| 227,1255 | 23,62 | SIRIUS | FYRIOAKWAWHYCM-UHFFFAOYSA-N | Organic compounds | Organoheterocyclic compounds | Azoles |
| 274,0863 | 23,74 | ChemWalker | WEHXAEGTVPWKDY-UHFFFAOYSA-O | Organic compounds | Organoheterocyclic compounds | Naphthopyrans |
| 295,1443 | 23,81 | ChemWalker | RVGGQPXRIAXKDV-UHFFFAOYSA-N | Organic compounds | Organoheterocyclic compounds | Benzodiazepines |
| 212,0820 | 23,87 | ChemWalker | PZRHRDRVRGEVNW-UHFFFAOYSA-N | Organic compounds | Organoheterocyclic compounds | Pyridines and derivatives |
| 398,3418 | 24,14 | ChemWalker | PAKZQLJDHCBPOR-UHFFFAOYSA-N | Organic compounds | Lipids and lipid-like molecules | Steroids and steroid derivatives |
| 449,2018 | 24,16 | ChemWalker | REGBMADJTTZONK-UHFFFAOYSA-N | Organic compounds | Organic acids and derivatives | Carboxylic acids and derivatives |
| 209,1150 | 24,21 | ChemWalker | WQWQHCFZHDDBLY-UHFFFAOYSA-N | Organic compounds | Organoheterocyclic compounds | Pyrans |
| 400,3574 | 24,41 | ChemWalker | SWTXHUUBYZNDAJ-UHFFFAOYSA-N | Organic compounds | Lipids and lipid-like molecules | Steroids and steroid derivatives |
| 259,1443 | 24,58 | ChemWalker | MVNBAVQHXXMXAO-UHFFFAOYSA-N | Organic compounds | Organic acids and derivatives | Carboxylic acids and derivatives |
| 203,1277 | 24,50 | ChemWalker | QNCYQSIQAGTLJX-UHFFFAOYSA-N | Organic compounds | Lipids and lipid-like molecules | Fatty Acyls |
| 211,0888 | 24,55 | ChemWalker | RZWQZVWQFRVQCI-UHFFFAOYSA-N | Organic compounds | Alkaloids and derivatives | Harmala alkaloids |
| 209,1150 | 24,52 | ChemWalker | WQWQHCFZHDDBLY-UHFFFAOYSA-N | Organic compounds | Organoheterocyclic compounds | Pyrans |
| 219,1743 | 24,63 | ChemWalker | GQUGDHVDNITJEZ-UHFFFAOYSA-N | Organic compounds | Lipids and lipid-like molecules | Prenol lipids |
| 223,0960 | 24,70 | GNPS2 | FLKPEMZONWLCSK-UHFFFAOYSA-N | Organic compounds | Benzenoids | Benzene and substituted derivatives |
| 258,1122 | 24,71 | ChemWalker | HRXAUDACLOQULD-UHFFFAOYSA-N | Organic compounds | Organic oxygen compounds | Organooxygen compounds |
| 259,1442 | 24,77 | ChemWalker | QHLKSESIIKJUHS-UHFFFAOYSA-N | Organic compounds | Benzenoids | Phenols |
| 223,1309 | 24,90 | GNPS2 | FLKPEMZONWLCSK-UHFFFAOYSA-N | Organic compounds | Benzenoids | Benzene and substituted derivatives |
| 400,3575 | 25,02 | ChemWalker | SWTXHUUBYZNDAJ-UHFFFAOYSA-N | Organic compounds | Lipids and lipid-like molecules | Steroids and steroid derivatives |
| 421,3063 | 25,12 | ChemWalker | PMKFHUDSBGNHST-UHFFFAOYSA-N | Organic compounds | Organoheterocyclic compounds | Dihydrofurans |
| 404,2797 | 25,13 | GNPS2 | FRIRHJVKXFYECW-ZTERCDMUSA-N | Organic compounds | Lipids and lipid-like molecules | Steroids and steroid derivatives |
| 386,2691 | 25,13 | ChemWalker | FCPKYEAPHJRRQX-UHFFFAOYSA-N | Organic compounds | Organoheterocyclic compounds | Oxanes |
| 535,2387 | 25,18 | ChemWalker | UDDLQPQKIJUXHP-UHFFFAOYSA-N | Organic compounds | Organic acids and derivatives | Peptidomimetics |
| 552,2652 | 25,18 | ChemWalker | WOYXZSPIKXPEHK-UHFFFAOYSA-N | Organic compounds | Organoheterocyclic compounds | Quinolines and derivatives |
| 557,2203 | 25,19 | ChemWalker | HYWNOKPRHXEGEO-UHFFFAOYSA-N | Organic compounds | Phenylpropanoids and polyketides | Diarylheptanoids |
| 223,1320 | 25,26 | GNPS2 | MXJWRABVEGLYDG-UHFFFAOYSA-N | Organic compounds | Benzenoids | Benzene and substituted derivatives |
| 261,0883 | 25,26 | ChemWalker | ACBYMHDZGRHKBL-UHFFFAOYSA-N | Organic compounds | Organic oxygen compounds | Organooxygen compounds |
| 240,1593 | 25,26 | ChemWalker | JDECRXOMHIGGFQ-UHFFFAOYSA-N | Organic compounds | Benzenoids | Benzene and substituted derivatives |
| 507,1718 | 25,26 | ChemWalker | IFYRJQZTJFYPFS-UHFFFAOYSA-N | Organic compounds | Phenylpropanoids and polyketides | Diarylheptanoids |
| 353,1694 | 25,26 | ChemWalker | IILDIIAZNITTLB-UHFFFAOYSA-N | Organic compounds | Benzenoids | Benzene and substituted derivatives |
| 705,3304 | 25,26 | ChemWalker | MEWNUFWOAOFSNB-UHFFFAOYSA-N | Organic compounds | Phenylpropanoids and polyketides | Macrolactams |
| 483,2054 | 25,27 | ChemWalker | LGPWEDBFIDYQAL-UHFFFAOYSA-N | Organic compounds | Phenylpropanoids and polyketides | Coumarins and derivatives |
| 331,0517 | 25,27 | ChemWalker | KCKLGAFEWGJTHI-UHFFFAOYSA-N | Organic compounds | Lipids and lipid-like molecules | Fatty Acyls |
| 307,0852 | 25,27 | ChemWalker | DUGLQGAPHNTGBR-UHFFFAOYSA-N | Organic compounds | Organoheterocyclic compounds | Benzodiazepines |
| 408,3110 | 25,32 | ChemWalker | BWNYFNVEJYUZSV-UHFFFAOYSA-N | Organic compounds | Phenylpropanoids and polyketides | Macrolide lactams |
| 354,2795 | 25,32 | ChemWalker | MNBYKMIRWHPDPD-UHFFFAOYSA-N | Organic compounds | Benzenoids | Benzene and substituted derivatives |
| 273,1600 | 25,46 | ChemWalker | OLTCJZPAHRXNGH-UHFFFAOYSA-N | Organic compounds | Organic nitrogen compounds | Organonitrogen compounds |
| 223,1320 | 25,57 | ChemWalker | GXWKLPJDISLIRF-UHFFFAOYSA-N | Organic compounds | Organoheterocyclic compounds | Pyrans |
| 209,1518 | 25,75 | ChemWalker | YWLLXHWMXNPEPY-UHFFFAOYSA-N | Organic compounds | Lipids and lipid-like molecules | Prenol lipids |
| 274,0862 | 25,76 | GNPS2 | MUMCCPUVOAUBAN-UHFFFAOYSA-N | Organic compounds | Alkaloids and derivatives | Aporphines |
| 225,1959 | 25,81 | ChemWalker | ADFXKUOMJKEIND-UHFFFAOYSA-N | Organic compounds | Organic acids and derivatives | Organic carbonic acids and derivatives |
| 203,1278 | 25,91 | ChemWalker | QNCYQSIQAGTLJX-UHFFFAOYSA-N | Organic compounds | Lipids and lipid-like molecules | Fatty Acyls |
| 217,1434 | 25,92 | ChemWalker | KLWFGMYQPMZELD-UHFFFAOYSA-N | Organic compounds | Organoheterocyclic compounds | Benzopyrans |
| 643,2568 | 26,00 | ChemWalker | IIXVPTUAJVHQCM-UHFFFAOYSA-N | Organic compounds | Lignans, neolignans and related compounds | None |
| 621,2752 | 26,00 | ChemWalker | AMASLVSVWKIJNI-UHFFFAOYSA-N | Organic compounds | Organic acids and derivatives | Carboxylic acids and derivatives |
| 235,1687 | 25,92 | ChemWalker | OCIQDJYHPBDJQX-UHFFFAOYSA-N | Organic compounds | Lipids and lipid-like molecules | Fatty Acyls |
| 209,1532 | 26,07 | ChemWalker | GIYINLRNLNHSBA-UHFFFAOYSA-N | Organic compounds | Lipids and lipid-like molecules | Fatty Acyls |
| 352,2635 | 26,14 | ChemWalker | DEBMTRRCNAXGIC-UHFFFAOYSA-N | Organic compounds | Organoheterocyclic compounds | Piperidines |
| 370,2741 | 26,14 | ChemWalker | SSBAWYYRBSCSAF-UHFFFAOYSA-N | Organic compounds | Organic acids and derivatives | Carboxylic acids and derivatives |
| 388,2847 | 26,14 | GNPS2 | COCMFMBNEAMQMA-KREOYVNCSA-N | Organic compounds | Lipids and lipid-like molecules | Steroids and steroid derivatives |
| 406,2951 | 26,14 | GNPS2 | RHCPKKNRWFXMAT-RRWYKFPJSA-N | Organic compounds | Lipids and lipid-like molecules | Steroids and steroid derivatives |
| 608,9394 | 26,14 | SIRIUS | HFFZUIBKLUIHNA-CVKSISIWSA-N | None | None | None |
| 609,4411 | 26,14 | ChemWalker | MBQUDSOKTNSZGX-UHFFFAOYSA-N | Organic compounds | Alkaloids and derivatives | None |
| 627,9129 | 26,15 | SIRIUS | RQWXYMCLLKFITD-FXMZOFOKSA-N | Organic compounds | Organic acids and derivatives | Carboxylic acids and derivatives |
| 811,5819 | 26,15 | ChemWalker | RWKDTQKIZDLDQT-UHFFFAOYSA-N | Organic compounds | Lipids and lipid-like molecules | Prenol lipids |
| 423,3217 | 26,15 | ChemWalker | GEQWUBLKSRVCLJ-UHFFFAOYSA-N | Organic compounds | Lipids and lipid-like molecules | Prenol lipids |
| 425,2690 | 26,16 | ChemWalker | VUTCKOBMFJDHLW-UHFFFAOYSA-N | Organic compounds | Lipids and lipid-like molecules | Prenol lipids |
| 490,2475 | 26,16 | ChemWalker | IIIFDCYKXMYQHI-UHFFFAOYSA-N | Organic compounds | Organoheterocyclic compounds | Indoles and derivatives |
| 317,1358 | 26,24 | ChemWalker | PUYKVAACYDFUCQ-UHFFFAOYSA-N | Organic compounds | Lipids and lipid-like molecules | Prenol lipids |
| 295,1536 | 26,23 | ChemWalker | RYPQSGURZSTFSX-UHFFFAOYSA-N | Organic compounds | Benzenoids | Benzene and substituted derivatives |
| 288,2897 | 26,26 | GNPS2 | AVFIYMSJDDGDBQ-UHFFFAOYSA-N | Organic compounds | Lipids and lipid-like molecules | Prenol lipids |
| 269,0920 | 26,30 | ChemWalker | ASMQHDFMJKDMQZ-UHFFFAOYSA-N | Organic compounds | Organoheterocyclic compounds | Diazanaphthalenes |
| 226,2166 | 26,39 | ChemWalker | VKAMZEDHHWTTNZ-UHFFFAOYSA-N | Organic compounds | Organoheterocyclic compounds | Lactams |
| 288,2896 | 26,48 | GNPS2 | AOMUHOFOVNGZAN-UHFFFAOYSA-N | Organic compounds | Lipids and lipid-like molecules | Fatty Acyls |
| 230,2477 | 26,51 | ChemWalker | WMUMHAZHWIUBPN-UHFFFAOYSA-N | Organic compounds | Organic nitrogen compounds | Organonitrogen compounds |
| 255,1223 | 26,52 | ChemWalker | ZJPNWNNXUHAYDP-UHFFFAOYSA-N | Organic compounds | Organoheterocyclic compounds | Pyrans |
| 349,1407 | 26,52 | ChemWalker | YOGCCOGHOCSMQE-UHFFFAOYSA-N | Organic compounds | Benzenoids | Phenols |
| 395,1376 | 26,53 | ChemWalker | LUESWCNVQRWLSN-UHFFFAOYSA-M | Organic compounds | Phenylpropanoids and polyketides | Coumarins and derivatives |
| 237,1118 | 26,53 | ChemWalker | NPEVDKHQEYSYTP-UHFFFAOYSA-N | Organic compounds | Organoheterocyclic compounds | Dihydrofurans |
| 328,2117 | 26,53 | ChemWalker | NKZJCHCKRDGVKG-UHFFFAOYSA-N | Organic compounds | Organoheterocyclic compounds | Pyrrolizidines |
| 333,1671 | 26,53 | ChemWalker | UZPHFSSGJMLBAY-UHFFFAOYSA-N | Organic compounds | Phenylpropanoids and polyketides | Coumarins and derivatives |
| 643,3441 | 26,53 | GNPS2 | CJGRGYBLAHPYOM-HOLMNUNMSA-N | Organic compounds | Phenylpropanoids and polyketides | Cinnamic acids and derivatives |
| 311,1852 | 26,53 | GNPS2 | ZKMLQDNHMSFULN-UHFFFAOYSA-N | Organic compounds | Phenylpropanoids and polyketides | Flavonoids |
| 201,1597 | 26,53 | GNPS2 | GIJHDGJRTUSBJR-UHFFFAOYSA-N | Organic compounds | Phenylpropanoids and polyketides | Coumarins and derivatives |
| 219,1742 | 26,53 | GNPS2 | LQZXAHKUYJVAAR-UHFFFAOYSA-N | Organic compounds | Organic acids and derivatives | Hydroxy acids and derivatives |
| 227,1639 | 26,59 | ChemWalker | ZKLGONMUSTXSKO-UHFFFAOYSA-N | Organic compounds | Organic oxygen compounds | Organooxygen compounds |
| 241,1734 | 26,57 | ChemWalker | OTFRJPUPCOSKKS-UHFFFAOYSA-N | Organic compounds | Alkaloids and derivatives | Harmala alkaloids |
| 389,2686 | 26,59 | ChemWalker | GXCUIVKBCLAWCS-UHFFFAOYSA-N | Organic compounds | Lipids and lipid-like molecules | Fatty Acyls |
| 201,1487 | 26,59 | ChemWalker | YFYDEZMIUOGZLE-UHFFFAOYSA-N | Organic compounds | Lipids and lipid-like molecules | Fatty Acyls |
| 239,1254 | 26,60 | GNPS2 | KZKWCKFDCPVDFJ-UHFFFAOYSA-N | Organic compounds | Phenylpropanoids and polyketides | Flavonoids |
| 209,1531 | 26,74 | ChemWalker | GIYINLRNLNHSBA-UHFFFAOYSA-N | Organic compounds | Lipids and lipid-like molecules | Fatty Acyls |
| 244,2634 | 26,75 | SIRIUS | BXNJVABOESKYKV-UHFFFAOYSA-O | Organic compounds | Organic nitrogen compounds | Organonitrogen compounds |
| 203,1277 | 26,46 | ChemWalker | ZJVMHPVIAUKERS-UHFFFAOYSA-N | Organic compounds | Lipids and lipid-like molecules | Fatty Acyls |
| 288,2897 | 26,86 | GNPS2 | KFQUQCFJDMSIJF-UHFFFAOYSA-N | Organic compounds | Organic nitrogen compounds | Organonitrogen compounds |
| 271,1442 | 26,87 | ChemWalker | WUTYHTAHLPCSRI-UHFFFAOYSA-N | Organic compounds | Organoheterocyclic compounds | Piperidines |
| 265,1796 | 26,89 | ChemWalker | QODRBRVKXLAPIS-UHFFFAOYSA-N | Organic compounds | Lipids and lipid-like molecules | Fatty Acyls |
| 339,1778 | 26,93 | ChemWalker | IIXIWBWRBCMJQB-UHFFFAOYSA-O | Organic compounds | Phenylpropanoids and polyketides | Aurone flavonoids |
| 447,3220 | 27,00 | ChemWalker | PJAPZIZSFGWFOQ-UHFFFAOYSA-N | Organic compounds | Lipids and lipid-like molecules | Prenol lipids |
| 396,1594 | 27,04 | ChemWalker | QJDBRFWHUXQNPC-UHFFFAOYSA-N | Organic compounds | Benzenoids | Benzene and substituted derivatives |
| 244,2634 | 27,07 | SIRIUS | JJRGVJZDAWBKMP-UHFFFAOYSA-N | Organic compounds | Organic nitrogen compounds | Organonitrogen compounds |
| 423,3219 | 27,10 | ChemWalker | GEQWUBLKSRVCLJ-UHFFFAOYSA-N | Organic compounds | Lipids and lipid-like molecules | Prenol lipids |
| 406,2952 | 27,10 | GNPS2 | RHCPKKNRWFXMAT-RRWYKFPJSA-N | Organic compounds | Lipids and lipid-like molecules | Steroids and steroid derivatives |
| 217,1434 | 27,11 | ChemWalker | MZSHZQAODGHFPQ-UHFFFAOYSA-N | Organic compounds | Lipids and lipid-like molecules | Fatty Acyls |
| 231,1591 | 27,11 | GNPS2 | ALOUNLDAKADEEB-UHFFFAOYSA-N | Organic compounds | Lipids and lipid-like molecules | Fatty Acyls |
| 304,2868 | 27,11 | SIRIUS | XYAAJTWJHPUIAG-UHFFFAOYSA-N | Organic compounds | Organic nitrogen compounds | Organonitrogen compounds |
| 229,1798 | 27,27 | ChemWalker | UEHUKETWTYBLMM-UHFFFAOYSA-N | Organic compounds | Lipids and lipid-like molecules | Fatty Acyls |
| 207,1357 | 27,30 | ChemWalker | XPMYTBIJWHMOIJ-UHFFFAOYSA-N | Organic compounds | Benzenoids | Benzene and substituted derivatives |
| 424,3062 | 27,39 | GNPS2 | IMMADCCLTPCOKH-HNWZYOJHSA-N | Organic compounds | Lipids and lipid-like molecules | Steroids and steroid derivatives |
| 448,3062 | 27,37 | ChemWalker | IVHIYFAYONFDBJ-UHFFFAOYSA-N | Organic compounds | Alkaloids and derivatives | Cytochalasans |
| 353,2477 | 27,38 | ChemWalker | LNMQCLOBYXKUCL-UHFFFAOYSA-N | Organic compounds | Lipids and lipid-like molecules | Fatty Acyls |
| 432,2382 | 27,39 | GNPS2 | WZYGIALDVOKLLL-UHFFFAOYSA-N | Organic compounds | Lipids and lipid-like molecules | Prenol lipids |
| 407,2793 | 27,39 | GNPS2 | RHCPKKNRWFXMAT-RRWYKFPJSA-N | Organic compounds | Lipids and lipid-like molecules | Steroids and steroid derivatives |
| 389,2689 | 27,39 | ChemWalker | GXCUIVKBCLAWCS-UHFFFAOYSA-N | Organic compounds | Lipids and lipid-like molecules | Fatty Acyls |
| 415,2115 | 27,39 | GNPS2 | OHNVJXDBOKZLFC-UHFFFAOYSA-N | Organic compounds | Lipids and lipid-like molecules | Prenol lipids |
| 221,1514 | 27,41 | ChemWalker | AGXXOOILRXZQTH-UHFFFAOYSA-N | Organic compounds | Lipids and lipid-like molecules | Fatty Acyls |
| 213,1849 | 27,66 | ChemWalker | GQVYBECSNBLQJV-UHFFFAOYSA-N | Organic compounds | Lipids and lipid-like molecules | Fatty Acyls |
| 265,1792 | 27,71 | ChemWalker | SZVNKXCDJUBPQO-UHFFFAOYSA-N | Organic compounds | Lipids and lipid-like molecules | Fatty Acyls |
| 316,3210 | 27,76 | GNPS2 | UPUIQOIQVMNQAP-UHFFFAOYSA-M | Organic compounds | Organic acids and derivatives | Organic sulfuric acids and derivatives |
| 219,1744 | 27,76 | GNPS2 | NCCWCZLEACWJIN-UHFFFAOYSA-N | Organic compounds | Benzenoids | Benzene and substituted derivatives |
| 259,1669 | 27,77 | ChemWalker | IGNIYJTXYTUVGD-UHFFFAOYSA-N | Organic compounds | Lipids and lipid-like molecules | Fatty Acyls |
| 495,3445 | 27,77 | ChemWalker | MQJYRDGWGIZCIV-UHFFFAOYSA-N | Organic compounds | Organoheterocyclic compounds | Benzopyrans |
| 201,1625 | 27,77 | GNPS2 | GIJHDGJRTUSBJR-UHFFFAOYSA-N | Organic compounds | Phenylpropanoids and polyketides | Coumarins and derivatives |
| 360,2439 | 27,88 | ChemWalker | ZUFGZIXTZJSHND-UHFFFAOYSA-O | Organic compounds | Organic oxygen compounds | Organooxygen compounds |
| 375,2144 | 27,88 | ChemWalker | UKRCOMRGFCFHQM-UHFFFAOYSA-N | Organic compounds | Lipids and lipid-like molecules | Fatty Acyls |
| 316,3211 | 27,94 | GNPS2 | UPUIQOIQVMNQAP-UHFFFAOYSA-M | Organic compounds | Organic acids and derivatives | Organic sulfuric acids and derivatives |
| 343,2956 | 27,98 | GNPS2 | MRUAUOIMASANKQ-UHFFFAOYSA-N | Organic compounds | Organic acids and derivatives | Carboxylic acids and derivatives |
| 240,2323 | 27,98 | ChemWalker | IPGLKLMWYXMYAB-UHFFFAOYSA-N | Organic compounds | Organic nitrogen compounds | Organonitrogen compounds |
| 323,0987 | 27,99 | ChemWalker | HGNCZWIESFNTCT-UHFFFAOYSA-N | Organic compounds | Organoheterocyclic compounds | Azolidines |
| 258,2793 | 27,99 | ChemWalker | DJCPHBCQLFTQBT-UHFFFAOYSA-O | Organic compounds | Organic nitrogen compounds | Organonitrogen compounds |
| 372,2897 | 28,04 | GNPS2 | JWZBXKZZDYMDCJ-IJPFKRJSSA-N | Organic compounds | Lipids and lipid-like molecules | Steroids and steroid derivatives |
| 390,3004 | 28,04 | GNPS2 | DKPMWHFRUGMUKF-HJNWHKLXSA-N | Organic compounds | Lipids and lipid-like molecules | Steroids and steroid derivatives |
| 408,3110 | 28,04 | ChemWalker | BWNYFNVEJYUZSV-UHFFFAOYSA-N | Organic compounds | Phenylpropanoids and polyketides | Macrolide lactams |
| 425,3375 | 28,04 | GNPS2 | BHQCQFFYRZLCQQ-OELDTZBJSA-N | Organic compounds | Lipids and lipid-like molecules | Steroids and steroid derivatives |
| 492,2633 | 28,04 | ChemWalker | BCGQJEGLKJUGLG-UHFFFAOYSA-N | Organic compounds | Phenylpropanoids and polyketides | Macrolides and analogues |
| 203,1269 | 28,05 | ChemWalker | OAKQXDBQNMCTKD-UHFFFAOYSA-N | Organic compounds | Organic acids and derivatives | Hydroxy acids and derivatives |
| 324,2171 | 28,09 | ChemWalker | JTRRPZVFSRBZIV-UHFFFAOYSA-N | Organic compounds | Lipids and lipid-like molecules | Prenol lipids |
| 231,1592 | 28,13 | ChemWalker | QHCWCHOUDLMCMJ-UHFFFAOYSA-N | Organic compounds | Organoheterocyclic compounds | Pyridines and derivatives |
| 239,1257 | 28,15 | ChemWalker | JWSADSZPLGCVPA-UHFFFAOYSA-N | Organic compounds | Benzenoids | Benzene and substituted derivatives |
| 245,1748 | 28,14 | ChemWalker | APOIUMVPTCBLQD-UHFFFAOYSA-N | Organic compounds | Lipids and lipid-like molecules | Prenol lipids |
| 235,1676 | 28,16 | ChemWalker | ZKNLFTOBEMGDRP-UHFFFAOYSA-N | Organic compounds | Organic oxygen compounds | Organooxygen compounds |
| 272,2948 | 28,17 | ChemWalker | UGVBFHUWZNNKIK-UHFFFAOYSA-N | Organic compounds | Organic nitrogen compounds | Organonitrogen compounds |
| 207,0177 | 28,13 | ChemWalker | NZQAQAUWFHMVEM-UHFFFAOYSA-M | Organic compounds | Phenylpropanoids and polyketides | Coumarins and derivatives |
| 316,3211 | 28,25 | GNPS2 | UPUIQOIQVMNQAP-UHFFFAOYSA-M | Organic compounds | Organic acids and derivatives | Organic sulfuric acids and derivatives |
| 243,1955 | 28,26 | ChemWalker | MRITWMKUVGBKRX-UHFFFAOYSA-N | Organic compounds | Lipids and lipid-like molecules | Fatty Acyls |
| 579,3625 | 28,31 | ChemWalker | QUJILRREMWWJDX-UHFFFAOYSA-O | Organic compounds | Benzenoids | Phenols |
| 389,1118 | 28,31 | ChemWalker | YHFVEOLVMJRQPS-UHFFFAOYSA-N | Organic compounds | Phenylpropanoids and polyketides | Depsides and depsidones |
| 223,1319 | 28,31 | GNPS2 | MXJWRABVEGLYDG-UHFFFAOYSA-N | Organic compounds | Benzenoids | Benzene and substituted derivatives |
| 296,2222 | 28,31 | ChemWalker | LITCXFPAGLBVLT-UHFFFAOYSA-O | Organic compounds | Organic nitrogen compounds | Organonitrogen compounds |
| 301,1775 | 28,31 | ChemWalker | JZAGMCTVWGDPFY-UHFFFAOYSA-N | Organic compounds | Organic oxygen compounds | Organooxygen compounds |
| 363,1481 | 28,31 | ChemWalker | SNYUMMZVPJTQOU-UHFFFAOYSA-N | Organic compounds | Lipids and lipid-like molecules | Prenol lipids |
| 387,1146 | 28,31 | ChemWalker | OOGIIAPQPDUUCX-UHFFFAOYSA-N | Organic compounds | Benzenoids | Benzene and substituted derivatives |
| 595,3310 | 28,31 | ChemWalker | LMTNTLPLGZVRFW-UHFFFAOYSA-N | Organic compounds | Benzenoids | Phenols |
| 387,3220 | 28,37 | ChemWalker | APKIEHGOXFERQX-UHFFFAOYSA-N | Organic compounds | Lipids and lipid-like molecules | Fatty Acyls |
| 249,1847 | 28,38 | ChemWalker | BGMHTWAIDRNJIY-UHFFFAOYSA-N | Organic compounds | Organic oxygen compounds | Organooxygen compounds |
| 408,3111 | 28,40 | ChemWalker | YPJUZEMVAYYXAV-UHFFFAOYSA-N | Organic compounds | Lipids and lipid-like molecules | Fatty Acyls |
| 430,2932 | 28,28 | SIRIUS | XEUKTMAQQWUUIX-GVRJEKJASA-N | None | None | None |
| 332,3188 | 28,45 | SIRIUS | XPEZYZKFQYNWAB-UHFFFAOYSA-N | Organic compounds | Organic nitrogen compounds | Organonitrogen compounds |
| 367,2094 | 28,53 | ChemWalker | WUNLLCOJYRGHCJ-UHFFFAOYSA-N | Organic compounds | Phenylpropanoids and polyketides | Macrolides and analogues |
| 448,3059 | 28,58 | GNPS2 | WVULKSPCQVQLCU-BUXLTGKBSA-N | Organic compounds | Lipids and lipid-like molecules | Steroids and steroid derivatives |
| 225,1098 | 28,84 | ChemWalker | YUODRAVGLILWNO-UHFFFAOYSA-N | Organic compounds | Organic oxygen compounds | Organooxygen compounds |
| 452,3218 | 28,73 | GNPS2 | INRCFVYVWPWZJS-IWQZZHSRSA-N | Organic compounds | Benzenoids | Phenols |
| 435,2955 | 28,73 | GNPS2 | SCABKEBYDRTODC-UHFFFAOYSA-N | Organic compounds | Lipids and lipid-like molecules | Fatty Acyls |
| 227,2006 | 28,75 | ChemWalker | ZVXDGKJSUPWREP-UHFFFAOYSA-N | Organic compounds | Lipids and lipid-like molecules | Fatty Acyls |
| 475,2305 | 28,81 | ChemWalker | VBYRVFSKLIRPLQ-UHFFFAOYSA-N | Organic compounds | Organoheterocyclic compounds | Indoles and derivatives |
| 223,0962 | 28,82 | GNPS2 | FLKPEMZONWLCSK-UHFFFAOYSA-N | Organic compounds | Benzenoids | Benzene and substituted derivatives |
| 205,0858 | 28,82 | ChemWalker | JTZDCGBXBVCDGU-UHFFFAOYSA-N | Organic compounds | Lipids and lipid-like molecules | Fatty Acyls |
| 279,1591 | 28,82 | GNPS2 | DOIRQSBPFJWKBE-UHFFFAOYSA-N | Organic compounds | Benzenoids | Benzene and substituted derivatives |
| 296,1854 | 28,82 | ChemWalker | ZDXGBZAVJULFIE-UHFFFAOYSA-N | Organic compounds | Organic acids and derivatives | Carboxylic acids and derivatives |
| 301,1410 | 28,82 | ChemWalker | PIKRTEPAODHCST-UHFFFAOYSA-N | Organic compounds | Organoheterocyclic compounds | Benzopyrans |
| 387,0780 | 28,82 | ChemWalker | GJUSBEFYAJEOLG-UHFFFAOYSA-N | Organic compounds | Benzenoids | Indanes |
| 579,2923 | 28,82 | GNPS2 | RPMNUQRUHXIGHK-UHFFFAOYSA-N | Organic compounds | Phenylpropanoids and polyketides | Flavonoids |
| 205,0858 | 28,86 | ChemWalker | JTZDCGBXBVCDGU-UHFFFAOYSA-N | Organic compounds | Lipids and lipid-like molecules | Fatty Acyls |
| 387,0780 | 28,86 | ChemWalker | QDLWVCARPPNRNL-UHFFFAOYSA-N | Organic compounds | Organoheterocyclic compounds | Lactones |
| 245,1748 | 29,03 | ChemWalker | REGGDLIBDASKGE-UHFFFAOYSA-N | Organic compounds | Lipids and lipid-like molecules | Fatty Acyls |
| 259,1905 | 29,03 | ChemWalker | ODDJEGGQRCHIDQ-UHFFFAOYSA-N | Organic compounds | Lipids and lipid-like molecules | Prenol lipids |
| 449,3375 | 29,06 | GNPS2 | GHCZAUBVMUEKKP-GYPHWSFCSA-N | Organic compounds | Lipids and lipid-like molecules | Steroids and steroid derivatives |
| 471,3196 | 29,06 | ChemWalker | JDSTWRDZSUZODG-UHFFFAOYSA-N | Organic compounds | Phenylpropanoids and polyketides | Coumarins and derivatives |
| 245,1151 | 29,04 | ChemWalker | LRDFOPNJXHWGGG-UHFFFAOYSA-N | Organic compounds | Hydrocarbon derivatives | Tropones |
| 341,3527 | 29,12 | SIRIUS | BDHJUCZXTYXGCZ-UHFFFAOYSA-N | Organic compounds | Lipids and lipid-like molecules | Fatty Acyls |
| 402,3579 | 29,21 | GNPS2 | PEUUVVGQIVMSAW-RZTYQLBFSA-N | Organic compounds | Lignans, neolignans and related compounds | Furanoid lignans |
| 295,1667 | 29,25 | ChemWalker | LLXQOSKXNCKPPE-UHFFFAOYSA-N | Organic compounds | Organic acids and derivatives | Organic sulfonic acids and derivatives |
| 391,2847 | 29,26 | ChemWalker | SVMBBDHQQNUOEU-UHFFFAOYSA-N | Organic compounds | Lipids and lipid-like molecules | Prenol lipids |
| 383,2043 | 29,27 | ChemWalker | BBAISYCWVQINOR-UHFFFAOYSA-N | Organic compounds | Lipids and lipid-like molecules | Prenol lipids |
| 414,3004 | 29,30 | ChemWalker | WDQJXJLCNREZMK-UHFFFAOYSA-O | Organic compounds | Organic acids and derivatives | Carboxylic acids and derivatives |
| 432,3111 | 29,31 | ChemWalker | KJDLCCSAVYUQIT-UHFFFAOYSA-N | Organic compounds | Organoheterocyclic compounds | Lactones |
| 450,3215 | 29,31 | GNPS2 | WVULKSPCQVQLCU-XMPMLJJQSA-N | Organic compounds | Lipids and lipid-like molecules | Steroids and steroid derivatives |
| 472,3021 | 29,31 | ChemWalker | SWKIBYOFRNTIGC-UHFFFAOYSA-N | Organic compounds | Organoheterocyclic compounds | Azepines |
| 355,2633 | 29,35 | ChemWalker | HAJOVCDDHUOYQO-UHFFFAOYSA-N | Organic compounds | Lipids and lipid-like molecules | Steroids and steroid derivatives |
| 271,2269 | 29,40 | ChemWalker | RPQJFAIEPUKQHK-UHFFFAOYSA-N | Organic compounds | Lipids and lipid-like molecules | Fatty Acyls |
| 295,1881 | 29,64 | ChemWalker | JPFXYNDBKFIYLX-UHFFFAOYSA-N | Organic compounds | Lipids and lipid-like molecules | Prenol lipids |
| 203,1794 | 29,45 | GNPS2 | QJAPFSSVKIZTMR-UHFFFAOYSA-N | Organic compounds | Organic oxygen compounds | Organooxygen compounds |
| 428,3736 | 29,49 | ChemWalker | GSYGVFBYRIDTHY-UHFFFAOYSA-N | Organic compounds | Lipids and lipid-like molecules | Fatty Acyls |
| 310,2379 | 29,55 | ChemWalker | CELZEZYUAVUZRU-UHFFFAOYSA-N | Organic compounds | Organoheterocyclic compounds | Pyrrolidines |
| 293,1747 | 29,57 | GNPS2 | YUTPMTLNRLENJI-UHFFFAOYSA-N | Organic compounds | Organic acids and derivatives | Carboxylic acids and derivatives |
| 315,1568 | 29,57 | ChemWalker | SBEJNBDOWAVCMR-UHFFFAOYSA-N | Organic compounds | Phenylpropanoids and polyketides | Coumarins and derivatives |
| 313,2351 | 29,57 | ChemWalker | HIEAMEODUHHHLB-UHFFFAOYSA-N | Organic compounds | Organic oxygen compounds | Organooxygen compounds |
| 291,2531 | 29,58 | GNPS2 | MMMVWBXLRFTTSV-UHFFFAOYSA-N | Organic compounds | Organic oxygen compounds | Organooxygen compounds |
| 371,3270 | 29,60 | GNPS2 | QGCUAFIULMNFPJ-UHFFFAOYSA-O | Organic compounds | Organic acids and derivatives | Carboxylic acids and derivatives |
| 273,2063 | 29,71 | ChemWalker | BYJHCUIKVIAEAC-UHFFFAOYSA-N | Organic compounds | Lipids and lipid-like molecules | Fatty Acyls |
| 295,1880 | 29,71 | ChemWalker | JPFXYNDBKFIYLX-UHFFFAOYSA-N | Organic compounds | Lipids and lipid-like molecules | Prenol lipids |
| 255,2320 | 29,72 | ChemWalker | KVXIRQZWCOAYRD-UHFFFAOYSA-N | Organic compounds | Lipids and lipid-like molecules | Fatty Acyls |
| 287,2219 | 29,76 | SIRIUS | VSQSFRDZIRKBEF-UHFFFAOYSA-N | Organic compounds | Lipids and lipid-like molecules | Fatty Acyls |
| 304,2484 | 29,76 | SIRIUS | LCBKXMNAPNKXEQ-UHFFFAOYSA-N | Organic compounds | Organic nitrogen compounds | Organonitrogen compounds |
| 293,2095 | 29,83 | ChemWalker | LRYVDYNQYBMNJP-UHFFFAOYSA-N | Organic compounds | Organic oxygen compounds | Organooxygen compounds |
| 245,1150 | 29,97 | ChemWalker | RHJDGWCFESNSMW-UHFFFAOYSA-N | Organic compounds | Benzenoids | Benzene and substituted derivatives |
| 425,2148 | 30,04 | ChemWalker | PVKOJQHHDYMUEI-UHFFFAOYSA-N | Organic compounds | Lipids and lipid-like molecules | Prenol lipids |
| 295,2246 | 30,04 | ChemWalker | YVWMHFYOIJMUMN-UHFFFAOYSA-N | Organic compounds | Lipids and lipid-like molecules | Fatty Acyls |
| 255,2320 | 30,05 | ChemWalker | LLPIORNGAIETEI-UHFFFAOYSA-N | Organic compounds | Lipids and lipid-like molecules | Fatty Acyls |
| 369,3841 | 30,06 | GNPS2 | FNLMCNKPGGHKQL-FPLPWBNLSA-N | Organic compounds | Benzenoids | Phenols |
| 385,3791 | 30,12 | ChemWalker | QRFANPLIZIVXBF-UHFFFAOYSA-N | Organic compounds | Lipids and lipid-like molecules | Steroids and steroid derivatives |
| 430,3894 | 30,13 | ChemWalker | UCROSHPNLXUYKT-UHFFFAOYSA-N | Organic compounds | Organoheterocyclic compounds | Diazines |
| 228,2325 | 30,21 | ChemWalker | NDWXIYYHLSYDIT-UHFFFAOYSA-N | Organic compounds | Organoheterocyclic compounds | Oxanes |
| 307,1907 | 30,22 | ChemWalker | ZICLWBMRDQUIDO-UHFFFAOYSA-N | Organic compounds | Benzenoids | Benzene and substituted derivatives |
| 329,1725 | 30,22 | ChemWalker | HUMPYOCQOJFKOK-UHFFFAOYSA-N | Organic compounds | Lipids and lipid-like molecules | Prenol lipids |
| 219,1016 | 30,22 | ChemWalker | IZOCZZMTQMWKLQ-UHFFFAOYSA-N | Organic compounds | Phenylpropanoids and polyketides | Coumarins and derivatives |
| 336,3110 | 30,30 | ChemWalker | YFSDFXHVCMRUBE-UHFFFAOYSA-O | Organic compounds | Lipids and lipid-like molecules | Prenol lipids |
| 355,2456 | 30,30 | ChemWalker | KSKGQTBFDMXQNE-UHFFFAOYSA-N | Organic compounds | Organic acids and derivatives | Carboxylic acids and derivatives |
| 341,2665 | 30,31 | ChemWalker | YLCXJNOKPGBEPM-UHFFFAOYSA-N | Organic compounds | Organic oxygen compounds | Organooxygen compounds |
| 245,1150 | 30,42 | ChemWalker | VICDYVFJMQPAHD-UHFFFAOYSA-N | Organic compounds | Organoheterocyclic compounds | Naphthofurans |
| 399,2720 | 30,38 | ChemWalker | LOYIDEKETTZLFY-UHFFFAOYSA-N | Organic compounds | Benzenoids | Phenols |
| 207,0178 | 30,60 | ChemWalker | NZQAQAUWFHMVEM-UHFFFAOYSA-M | Organic compounds | Phenylpropanoids and polyketides | Coumarins and derivatives |
| 326,2694 | 30,51 | ChemWalker | PPJMSWYZFNJSSW-UHFFFAOYSA-O | Organic compounds | Organoheterocyclic compounds | Quinolines and derivatives |
| 397,2951 | 30,66 | ChemWalker | NNPPLRJHBQPBAQ-UHFFFAOYSA-N | Organic compounds | Lipids and lipid-like molecules | Prenol lipids |
| 419,2770 | 30,67 | ChemWalker | GQGSZVZNLTULMP-UHFFFAOYSA-N | Organic compounds | Organic oxygen compounds | Organooxygen compounds |
| 414,3218 | 30,68 | ChemWalker | WAGYLURELCUJPG-UHFFFAOYSA-N | Organic compounds | Organic acids and derivatives | Hydroxy acids and derivatives |
| 441,3212 | 30,72 | ChemWalker | JTTPSRQRBBOCDL-UHFFFAOYSA-N | Organic compounds | Lipids and lipid-like molecules | Prenol lipids |
| 458,3476 | 30,73 | ChemWalker | XRODLBXBYOICFZ-UHFFFAOYSA-N | Organic compounds | Lipids and lipid-like molecules | Fatty Acyls |
| 463,3031 | 30,74 | ChemWalker | SYNJDZQPOAGZQA-UHFFFAOYSA-N | Organic compounds | Lipids and lipid-like molecules | Glycerolipids |
| 502,3738 | 30,77 | ChemWalker | ZEVICWRBRNCZED-UHFFFAOYSA-N | Organic compounds | Benzenoids | Phenols |
| 507,3292 | 30,77 | ChemWalker | YSRUKWIOXRIYRC-UHFFFAOYSA-N | Organic compounds | Phenylpropanoids and polyketides | Macrolides and analogues |
| 485,3474 | 30,77 | ChemWalker | SZISKKHIWHABQC-UHFFFAOYSA-N | Organic compounds | Lipids and lipid-like molecules | Prenol lipids |
| 523,3032 | 30,78 | ChemWalker | VKCJIMYQKDCKLH-UHFFFAOYSA-N | Organic compounds | Organic oxygen compounds | Organooxygen compounds |
| 546,4000 | 30,80 | ChemWalker | CZPDMRFCRFSYIT-UHFFFAOYSA-O | Organic compounds | Organic acids and derivatives | Hydroxy acids and derivatives |
| 552,3587 | 30,79 | ChemWalker | AJVSXHLKGPXWJH-UHFFFAOYSA-O | Organic compounds | Lipids and lipid-like molecules | Fatty Acyls |
| 567,3290 | 30,79 | ChemWalker | ASQRSYUZMDSTKU-UHFFFAOYSA-N | Organic compounds | Phenylpropanoids and polyketides | Macrolides and analogues |
| 529,3735 | 30,80 | ChemWalker | YUPHCPRFPDDUPD-UHFFFAOYSA-N | Organic compounds | Benzenoids | Benzene and substituted derivatives |
| 551,3552 | 30,81 | ChemWalker | VEKKIFQYJCIPMU-UHFFFAOYSA-N | Organic compounds | Lipids and lipid-like molecules | Prenol lipids |
| 590,4259 | 30,83 | ChemWalker | DUWJWUDMHJCAJI-UHFFFAOYSA-N | Organic compounds | Lipids and lipid-like molecules | Steroids and steroid derivatives |
| 596,3847 | 30,83 | ChemWalker | TZTMBPHEOFBDPY-UHFFFAOYSA-O | Organic compounds | Lipids and lipid-like molecules | Prenol lipids |
| 399,3584 | 30,83 | ChemWalker | DGFDSMDLLKKPHX-UHFFFAOYSA-N | Organic compounds | Lipids and lipid-like molecules | Steroids and steroid derivatives |
| 611,3551 | 30,84 | ChemWalker | KNJVBEGUJATRJI-UHFFFAOYSA-N | Organic compounds | Phenylpropanoids and polyketides | Diarylheptanoids |
| 634,4520 | 30,86 | ChemWalker | LTFHJSMCJZTLTP-UHFFFAOYSA-N | Organic compounds | Organic oxygen compounds | Organooxygen compounds |
| 640,4109 | 30,85 | ChemWalker | LZTUXOHRBPRKLZ-UHFFFAOYSA-N | Organic compounds | Phenylpropanoids and polyketides | Cinnamic acids and derivatives |
| 655,3812 | 30,86 | ChemWalker | OQWOKDQAPBSVGH-UHFFFAOYSA-N | Organic compounds | Lipids and lipid-like molecules | Glycerolipids |
| 678,4779 | 30,89 | ChemWalker | GRRAWGHVLYYPBU-UHFFFAOYSA-O | Organic compounds | Lipids and lipid-like molecules | Fatty Acyls |
| 684,4367 | 30,87 | ChemWalker | MDOMEJMFUHIERW-UHFFFAOYSA-N | Organic compounds | Organic acids and derivatives | Peptidomimetics |
| 722,5042 | 30,91 | ChemWalker | ZKHZGLULXDLLMK-UHFFFAOYSA-N | Organic compounds | Lipids and lipid-like molecules | Glycerophospholipids |
| 433,2926 | 31,05 | ChemWalker | CVLZBOJHINAXHY-UHFFFAOYSA-N | Organic compounds | Lipids and lipid-like molecules | Prenol lipids |
| 297,2401 | 31,05 | ChemWalker | BEBQWZBODPCAIK-UHFFFAOYSA-N | Organic compounds | Lipids and lipid-like molecules | Fatty Acyls |
| 245,1150 | 31,16 | ChemWalker | LRDFOPNJXHWGGG-UHFFFAOYSA-N | Organic compounds | Hydrocarbon derivatives | Tropones |
| 313,1712 | 31,41 | ChemWalker | OKWMTFYIQTVMIC-UHFFFAOYSA-O | Organic compounds | Organoheterocyclic compounds | Imidazopyrimidines |
| 477,3187 | 31,08 | ChemWalker | DXJLTHUHHKKFIH-UHFFFAOYSA-N | Organic compounds | Benzenoids | Benzene and substituted derivatives |
| 521,3451 | 31,10 | ChemWalker | KCXXIPPRZXVLRU-UHFFFAOYSA-N | Organic compounds | Organoheterocyclic compounds | Naphthofurans |
| 319,2845 | 31,11 | ChemWalker | VTIMCYVWFFHKIG-UHFFFAOYSA-N | Organic compounds | Lipids and lipid-like molecules | Fatty Acyls |
| 336,3110 | 31,11 | ChemWalker | YFSDFXHVCMRUBE-UHFFFAOYSA-O | Organic compounds | Lipids and lipid-like molecules | Prenol lipids |
| 341,2664 | 31,11 | ChemWalker | YLCXJNOKPGBEPM-UHFFFAOYSA-N | Organic compounds | Organic oxygen compounds | Organooxygen compounds |
| 380,3372 | 31,14 | SIRIUS | UYTRJTLZIHJHSV-HWOWSKLDSA-N | None | None | None |
| 385,2927 | 31,14 | ChemWalker | ADSNWNLTMACUJU-UHFFFAOYSA-N | Organic compounds | Organic acids and derivatives | Hydroxy acids and derivatives |
| 424,3633 | 31,18 | ChemWalker | QPMJZXUOGXCBCJ-UHFFFAOYSA-N | Organic compounds | Organoheterocyclic compounds | Pyrrolines |
| 429,3187 | 31,18 | ChemWalker | KOLAFJFESDNVMY-UHFFFAOYSA-N | Organic compounds | Lipids and lipid-like molecules | Fatty Acyls |
| 468,3896 | 31,21 | ChemWalker | ZGALAVFQYJOLRQ-UHFFFAOYSA-N | Organic compounds | Organoheterocyclic compounds | Quinolidines |
| 473,3451 | 31,21 | ChemWalker | ZCVSSDYMQMAXFL-UHFFFAOYSA-N | Organic compounds | Organic acids and derivatives | Carboxylic acids and derivatives |
| 512,4156 | 31,24 | SIRIUS | GRJWVVQBQJBYRR-QHCPKHFHSA-N | Organic compounds | Organic acids and derivatives | Carboxylic acids and derivatives |
| 517,3712 | 31,24 | ChemWalker | GXJRZJDDDHEKMP-UHFFFAOYSA-N | Organic compounds | Organic acids and derivatives | Hydroxy acids and derivatives |
| 556,4418 | 31,27 | ChemWalker | PWYYOLVACHXFGC-UHFFFAOYSA-N | Organic compounds | Organic oxygen compounds | Organooxygen compounds |
| 425,2875 | 31,29 | ChemWalker | FBFXIJVFPQRQJE-UHFFFAOYSA-N | Organic compounds | Lipids and lipid-like molecules | Fatty Acyls |
| 357,2031 | 31,30 | ChemWalker | JWJLIQZDNRPDFM-UHFFFAOYSA-N | None | None | None |
| 600,4679 | 31,29 | ChemWalker | NNKZUSAFPLLGKI-UHFFFAOYSA-N | Organic compounds | Lipids and lipid-like molecules | Prenol lipids |
| 386,3992 | 31,33 | SIRIUS | DYKUZSNJDWKTFZ-BJKOFHAPSA-N | Organic compounds | Organic nitrogen compounds | Organonitrogen compounds |
| 337,2350 | 31,37 | ChemWalker | WLPLVYBAMNYYSU-UHFFFAOYSA-N | Organic compounds | Benzenoids | Benzene and substituted derivatives |
| 339,2506 | 31,45 | ChemWalker | VIEQPAAMSLJMCL-UHFFFAOYSA-N | Organic compounds | Organoheterocyclic compounds | Pyrans |
| 579,3864 | 31,44 | ChemWalker | BKVVCDHXZMHTDM-UHFFFAOYSA-N | Organic compounds | Phenylpropanoids and polyketides | Macrolides and analogues |
| 491,3344 | 31,45 | ChemWalker | NQLGGBRFCXOFNZ-UHFFFAOYSA-N | Organic compounds | Lipids and lipid-like molecules | Prenol lipids |
| 535,3606 | 31,45 | ChemWalker | LKCGYKMSZBCFDL-UHFFFAOYSA-N | Organic compounds | Organoheterocyclic compounds | Diazanaphthalenes |
| 229,2164 | 31,51 | ChemWalker | JQEQRSJZNQFUBC-UHFFFAOYSA-N | Organic compounds | Lipids and lipid-like molecules | Fatty Acyls |
| 313,1689 | 31,52 | ChemWalker | SDEMLABHEIGMCS-UHFFFAOYSA-N | Organic compounds | Organoheterocyclic compounds | Benzopyrans |
| 400,4151 | 31,66 | SIRIUS | MSIUSKXETTXQIW-LOSJGSFVSA-N | Organic compounds | Organic nitrogen compounds | Organonitrogen compounds |
| 369,3843 | 31,76 | GNPS2 | FNLMCNKPGGHKQL-FPLPWBNLSA-N | Organic compounds | Benzenoids | Phenols |
| 593,4022 | 31,79 | ChemWalker | OOJRNONYBXTOMZ-UHFFFAOYSA-N | Organic compounds | Lipids and lipid-like molecules | Prenol lipids |
| 414,4309 | 31,82 | GNPS2 | PVAMXWLZJKTXFW-UHFFFAOYSA-N | Organic compounds | Lipids and lipid-like molecules | Steroids and steroid derivatives |
| 427,3898 | 31,82 | GNPS2 | RVGGCRQPGKFZDS-AKRYRNCVSA-N | Organic compounds | Phenylpropanoids and polyketides | Coumarins and derivatives |
| 255,2322 | 31,87 | ChemWalker | KVXIRQZWCOAYRD-UHFFFAOYSA-N | Organic compounds | Lipids and lipid-like molecules | Fatty Acyls |
| 277,2140 | 31,87 | GNPS2 | DOIRQSBPFJWKBE-UHFFFAOYSA-N | Organic compounds | Benzenoids | Benzene and substituted derivatives |
| 339,1840 | 31,87 | GNPS2 | WEJZLJFKUYPPDV-UHFFFAOYSA-N | Organic compounds | Phenylpropanoids and polyketides | Isoflavonoids |
| 466,5348 | 31,87 | SIRIUS | HGKGOKPTJMTGGN-UHFFFAOYSA-N | Organic compounds | Organic nitrogen compounds | Organonitrogen compounds |
| 348,3115 | 31,91 | GNPS2 | QHZLMUACJMDIAE-UHFFFAOYSA-N | Organic compounds | Lipids and lipid-like molecules | Glycerolipids |
| 353,2666 | 31,91 | ChemWalker | PUAPJGQBSUWLAY-UHFFFAOYSA-N | Organic compounds | Lipids and lipid-like molecules | Prenol lipids |
| 415,2371 | 31,91 | ChemWalker | KHCKIJOBWGIPFP-UHFFFAOYSA-N | Organic compounds | Lipids and lipid-like molecules | Saccharolipids |
| 331,2847 | 31,91 | GNPS2 | QDKBEJZVRCVOPZ-SDNWHVSQSA-N | Organic compounds | Lipids and lipid-like molecules | Fatty Acyls |
| 480,5507 | 31,93 | SIRIUS | HGKGOKPTJMTGGN-UHFFFAOYSA-N | Organic compounds | Organic nitrogen compounds | Organonitrogen compounds |
| 494,5660 | 31,98 | GNPS2 | FBRKYRSUSJWLHH-UHFFFAOYSA-N | Organic compounds | Alkaloids and derivatives | Emetine alkaloids |
| 508,5815 | 32,01 | SIRIUS | MQDVWHVLPUVVPQ-UHFFFAOYSA-N | Organic compounds | Organic nitrogen compounds | Organonitrogen compounds |
| 640,5868 | 32,03 | ChemWalker | UFCCKRLRJHVCPX-UHFFFAOYSA-N | Organic compounds | Organic oxygen compounds | Organooxygen compounds |
| 364,3424 | 32,06 | ChemWalker | UCGOBIVMMZGWNF-UHFFFAOYSA-N | Organic compounds | Organoheterocyclic compounds | Pyridines and derivatives |
| 370,3013 | 32,06 | GNPS2 | FNLMCNKPGGHKQL-FPLPWBNLSA-N | Organic compounds | Benzenoids | Phenols |
| 522,5971 | 32,06 | ChemWalker | SWZDQOUHBYYPJD-UHFFFAOYSA-N | Organic compounds | Organic nitrogen compounds | Organonitrogen compounds |
| 369,2980 | 32,06 | ChemWalker | GLIPXQAGIQSPTQ-UHFFFAOYSA-N | Organic compounds | Lipids and lipid-like molecules | Fatty Acyls |
| 413,3234 | 32,08 | ChemWalker | ZKFUKHCEACWDKL-UHFFFAOYSA-N | Organic compounds | Organic acids and derivatives | Carboxylic acids and derivatives |
| 457,3488 | 32,09 | ChemWalker | AIUNLQUMHTUTNR-UHFFFAOYSA-N | Organic compounds | Lipids and lipid-like molecules | Fatty Acyls |
| 367,2819 | 32,09 | ChemWalker | OIEZJKMVJYGTMT-UHFFFAOYSA-N | Organic compounds | Lipids and lipid-like molecules | Fatty Acyls |
| 501,3763 | 32,10 | ChemWalker | CVJWXRNBYKUGQI-UHFFFAOYSA-N | Organic compounds | Lipids and lipid-like molecules | Fatty Acyls |
| 536,6127 | 32,10 | SIRIUS | HFUXZBPRLMUEJG-UHFFFAOYSA-N | Organic compounds | Organic nitrogen compounds | Organonitrogen compounds |
| 668,6180 | 32,12 | GNPS2 | IKMYHRZEPWIULH-JJPXTECOSA-N | Organic compounds | Phenylpropanoids and polyketides | Coumarins and derivatives |
| 379,2820 | 32,14 | ChemWalker | LWXLRBPWKGXXED-UHFFFAOYSA-N | Organic compounds | Lipids and lipid-like molecules | Glycerolipids |
| 476,3017 | 32,14 | ChemWalker | PZBMKJNJSUCGAK-UHFFFAOYSA-N | Organic compounds | Lipids and lipid-like molecules | Prenol lipids |
| 550,6283 | 32,14 | SIRIUS | MIFZOICMMBUJKC-UHFFFAOYSA-N | Organic compounds | Organic nitrogen compounds | Organonitrogen compounds |
| 343,2855 | 32,23 | ChemWalker | VOQDFJUHTSKSNX-UHFFFAOYSA-N | Organic compounds | Organoheterocyclic compounds | Dioxanes |
| 341,2038 | 32,23 | ChemWalker | MTIAAZFBFDTPDE-UHFFFAOYSA-O | Organic compounds | Organoheterocyclic compounds | Imidazopyrimidines |
| 279,2293 | 32,23 | GNPS2 | DOIRQSBPFJWKBE-UHFFFAOYSA-N | Organic compounds | Benzenoids | Benzene and substituted derivatives |
| 359,2118 | 32,23 | ChemWalker | GVLRAQXHGZVPNP-UHFFFAOYSA-N | Organic compounds | Organic acids and derivatives | Carboxylic acids and derivatives |
| 551,4345 | 32,24 | ChemWalker | KPSZFRNHUVCRCO-UHFFFAOYSA-N | Organic compounds | Lipids and lipid-like molecules | Fatty Acyls |
| 685,4348 | 32,24 | ChemWalker | VUVQHSGDUUVSJD-UHFFFAOYSA-N | Organic compounds | Lignans, neolignans and related compounds | Dibenzylbutane lignans |
| 680,4793 | 32,24 | ChemWalker | SYSBJMYYTKWCBW-UHFFFAOYSA-N | Organic compounds | Lipids and lipid-like molecules | Steroids and steroid derivatives |
| 663,4528 | 32,25 | GNPS2 | JJSVXWKQLGHFES-QGZYWIHFSA-N | Organic compounds | Lipids and lipid-like molecules | Steroids and steroid derivatives |
| 381,2976 | 32,31 | ChemWalker | IKBDZTUBAQMHJD-UHFFFAOYSA-N | Organic compounds | Lipids and lipid-like molecules | Glycerolipids |
| 423,3083 | 32,33 | ChemWalker | PAXVNVHHFVFSMM-UHFFFAOYSA-N | Organic compounds | Lipids and lipid-like molecules | Prenol lipids |
| 339,1821 | 32,35 | ChemWalker | KUKDHFMUSAZTFT-UHFFFAOYSA-N | Organic compounds | Lipids and lipid-like molecules | Fatty Acyls |
| 474,3147 | 32,35 | ChemWalker | HXJDIWPGVZIJFH-UHFFFAOYSA-M | Organic compounds | Lipids and lipid-like molecules | Steroids and steroid derivatives |
| 339,2208 | 32,35 | ChemWalker | RPWFJAMTCNSJKK-UHFFFAOYSA-N | Organic compounds | Benzenoids | Benzene and substituted derivatives |
| 367,2152 | 32,35 | ChemWalker | CUTBIBOIOVRFHC-UHFFFAOYSA-N | Organic compounds | Organic acids and derivatives | Carboxylic acids and derivatives |
| 359,3158 | 32,40 | ChemWalker | ASKIVFGGGGIGKH-UHFFFAOYSA-N | Organic compounds | Lipids and lipid-like molecules | Glycerolipids |
| 379,2821 | 32,45 | ChemWalker | DLFKFFZPKVOYJI-UHFFFAOYSA-N | Organic compounds | Organic acids and derivatives | Organic sulfuric acids and derivatives |
| 803,5418 | 32,47 | GNPS2 | RFZOTNNDUHYGNN-RQBLKUNCSA-N | Organic compounds | Lipids and lipid-like molecules | Steroids and steroid derivatives |
| 414,2698 | 32,47 | ChemWalker | ISHIUSZGAFLPLK-UHFFFAOYSA-N | Organic compounds | Organoheterocyclic compounds | Pyrrolines |
| 391,2845 | 32,47 | GNPS2 | MQIUGAXCHLFZKX-UHFFFAOYSA-N | Organic compounds | Benzenoids | Benzene and substituted derivatives |
| 413,2663 | 32,47 | ChemWalker | YTHCLWOOCGCXRD-UHFFFAOYSA-N | Organic compounds | Organic oxygen compounds | Organooxygen compounds |
| 587,3548 | 32,47 | ChemWalker | FZSYIFKMFMIEJP-UHFFFAOYSA-N | Organic compounds | Lipids and lipid-like molecules | Prenol lipids |
| 450,3580 | 32,47 | SIRIUS | KYBCRMGJISMEBY-UHFFFAOYSA-N | None | None | None |
| 371,3159 | 32,52 | ChemWalker | JHXUIUBDCUTACR-UHFFFAOYSA-N | Organic compounds | Organoheterocyclic compounds | Dioxanes |
| 388,3423 | 32,52 | GNPS2 | SGTNXGSCHXKQJX-OQAPJDJNSA-N | Organic compounds | Organic oxygen compounds | Organooxygen compounds |
| 393,2972 | 32,52 | ChemWalker | XGXUFBAHOWHDNU-UHFFFAOYSA-N | Organic compounds | Benzenoids | Benzene and substituted derivatives |
| 409,2716 | 32,52 | ChemWalker | URXNJUNXUWLMAY-UHFFFAOYSA-N | Organic compounds | Lipids and lipid-like molecules | Prenol lipids |
| 625,9305 | 32,54 | SIRIUS | QGWNVMDLQWNPRV-UHFFFAOYSA-O | None | None | None |
| 603,9175 | 32,54 | SIRIUS | RDSIBICHPKRSLJ-UHFFFAOYSA-N | Organic compounds | Benzenoids | Benzene and substituted derivatives |
| 581,9045 | 32,55 | SIRIUS | JGKLKTXPJMJEMQ-UHFFFAOYSA-N | Organic compounds | Organoheterocyclic compounds | Furans |
| 559,8915 | 32,55 | SIRIUS | XJEDNRGSPMBTAG-UHFFFAOYSA-N | Organic compounds | Benzenoids | Phenol ethers |
| 537,8783 | 32,56 | SIRIUS | JNGLPGOVUHSSAJ-UHFFFAOYSA-N | Organic compounds | Organoheterocyclic compounds | Azolidines |
| 515,8655 | 32,56 | SIRIUS | NKPUDVODBKGAAF-UHFFFAOYSA-N | Organic compounds | Organic acids and derivatives | Carboxylic acids and derivatives |
| 397,3294 | 32,59 | ChemWalker | IYVZSZAMCRNRDJ-UHFFFAOYSA-N | Organic compounds | Organic oxygen compounds | Organooxygen compounds |
| 441,3552 | 32,59 | ChemWalker | RDKSFIAFDUECDH-UHFFFAOYSA-N | Organic compounds | Lipids and lipid-like molecules | Fatty Acyls |
| 485,3808 | 32,59 | ChemWalker | SYWAOYCXRQDRLP-UHFFFAOYSA-N | Organic compounds | Lipids and lipid-like molecules | Steroids and steroid derivatives |
| 379,2822 | 32,65 | ChemWalker | YLWJMUPPJKELEC-UHFFFAOYSA-N | Organic compounds | Lipids and lipid-like molecules | Fatty Acyls |
| 371,3158 | 32,69 | ChemWalker | JHXUIUBDCUTACR-UHFFFAOYSA-N | Organic compounds | Organoheterocyclic compounds | Dioxanes |
| 393,2977 | 32,69 | GNPS2 | CFXCGWWYIDZIMU-UHFFFAOYSA-N | Organic compounds | Lipids and lipid-like molecules | Fatty Acyls |
| 369,2313 | 32,73 | ChemWalker | ROUDCKODIMKLNO-UHFFFAOYSA-N | Organic compounds | Lipids and lipid-like molecules | Fatty Acyls |
| 387,2420 | 32,73 | ChemWalker | IARVMWOCDIKRTC-UHFFFAOYSA-N | Organic compounds | Lipids and lipid-like molecules | Fatty Acyls |
| 341,2364 | 32,74 | ChemWalker | IVJBKPMOAQVSDN-UHFFFAOYSA-N | Organic compounds | Organoheterocyclic compounds | Pyrans |
| 409,3287 | 32,79 | ChemWalker | QBFWTWXSXJFPRL-UHFFFAOYSA-N | Organic compounds | Lipids and lipid-like molecules | Prenol lipids |
| 401,3417 | 32,80 | ChemWalker | IRLPWWSRHJOKPQ-UHFFFAOYSA-N | Organic compounds | Organic acids and derivatives | Carboximidic acids and derivatives |
| 385,3317 | 32,85 | ChemWalker | VOVHGIJYIIRIQO-UHFFFAOYSA-N | Organic compounds | Organoheterocyclic compounds | Lactones |
| 407,3136 | 32,85 | ChemWalker | BGBDYUJDNLVMOV-UHFFFAOYSA-N | Organic compounds | Benzenoids | Benzene and substituted derivatives |
| 419,3155 | 32,91 | ChemWalker | UEFNZITZGVWLFK-UHFFFAOYSA-N | Organic compounds | Organic oxygen compounds | Organooxygen compounds |
| 423,3446 | 32,96 | ChemWalker | WPPKZMUCJIDWFE-UHFFFAOYSA-N | Organic compounds | Organic acids and derivatives | Hydroxy acids and derivatives |
| 705,9850 | 32,98 | SIRIUS | KPVUPGRLVYMICZ-UHFFFAOYSA-N | Organic compounds | Organic oxygen compounds | Organooxygen compounds |
| 683,9721 | 32,98 | SIRIUS | QPQAKAWUIXJOMD-UHFFFAOYSA-M | Organic compounds | Organoheterocyclic compounds | Thiophenes |
| 661,9592 | 32,99 | SIRIUS | STHBGJSFUVWVOR-LDADJPATSA-N | None | None | None |
| 639,9464 | 33,0 | SIRIUS | RVUDNOHNUHIOAB-CAMWMBJASA-N | Organic compounds | Benzenoids | Benzene and substituted derivatives |
| 617,9334 | 33,00 | SIRIUS | UUEMRKBFLBJBRG-UHFFFAOYSA-M | None | None | None |
| 595,9204 | 33,01 | SIRIUS | XHASFKKPXGYVKD-UHFFFAOYSA-N | None | None | None |
| 573,9074 | 33,02 | SIRIUS | BNPFJTOHLAAJEZ-UHFFFAOYSA-N | Organic compounds | Benzenoids | Benzene and substituted derivatives |
| 554,8740 | 33,02 | SIRIUS | WRUYQLARBOHHQQ-UHFFFAOYSA-N | Organic compounds | Benzenoids | Benzene and substituted derivatives |
| 551,8943 | 33,02 | SIRIUS | LDCKIIAHWMDXBX-UHFFFAOYSA-N | Organic compounds | Organoheterocyclic compounds | Bi- and oligothiophenes |
| 529,8812 | 33,03 | SIRIUS | DFGAEXSQQAETIG-UHFFFAOYSA-N | None | None | None |
| 532,8606 | 33,03 | SIRIUS | HCYHYRDHBCPKAV-UHFFFAOYSA-N | None | None | None |
| 507,8681 | 33,04 | SIRIUS | ORLSVWJGNDUAFC-LWELHUNNSA-N | Organic compounds | Organic acids and derivatives | Carboxylic acids and derivatives |
| 485,8548 | 33,05 | SIRIUS | JGWSYNPGVZYOIN-UHFFFAOYSA-N | Organic compounds | Organoheterocyclic compounds | Thiophenes |
| 488,3328 | 33,05 | SIRIUS | PYYQRZZRTSMIJA-KNJKTWCYSA-N | Organic compounds | Benzenoids | Benzene and substituted derivatives |
| 816,6393 | 33,06 | ChemWalker | XKJMKIJXBCHUJK-UHFFFAOYSA-O | Organic compounds | Lipids and lipid-like molecules | Fatty Acyls |
| 772,6134 | 33,07 | ChemWalker | LMGTVCKIUNTOEP-UHFFFAOYSA-N | Organic compounds | Lipids and lipid-like molecules | Glycerophospholipids |
| 684,5613 | 33,09 | SIRIUS | WUPIVEAEVYWLIH-BOWRCTOHSA-N | Organic compounds | Lipids and lipid-like molecules | Prenol lipids |
| 689,5166 | 33,09 | ChemWalker | OUICMEDLCFNSEM-UHFFFAOYSA-N | Organic compounds | Lipids and lipid-like molecules | Glycerolipids |
| 728,5872 | 33,09 | ChemWalker | RURFWJXZZZCIOZ-UHFFFAOYSA-N | Organic compounds | Lipids and lipid-like molecules | Sphingolipids |
| 645,4904 | 33,10 | ChemWalker | YGFHUDRQPPYWTP-UHFFFAOYSA-N | Organic compounds | Lipids and lipid-like molecules | Prenol lipids |
| 601,4653 | 33,11 | ChemWalker | LXVXTMHMMKQUMB-UHFFFAOYSA-N | Organic compounds | Lipids and lipid-like molecules | Steroids and steroid derivatives |
| 557,4385 | 33,11 | ChemWalker | HHCSYPMWJQHCMZ-UHFFFAOYSA-N | Organic compounds | Lipids and lipid-like molecules | Prenol lipids |
| 469,3859 | 33,12 | ChemWalker | UFLNXVJKZIUIOQ-UHFFFAOYSA-N | Organic compounds | Lipids and lipid-like molecules | Fatty Acyls |
| 513,4126 | 33,12 | ChemWalker | FKRYNOKEFDLDSY-UHFFFAOYSA-N | Organic compounds | Lipids and lipid-like molecules | Steroids and steroid derivatives |
| 425,3604 | 33,12 | ChemWalker | JJWZFUFNJNGKAF-UHFFFAOYSA-L | Organic compounds | Lipids and lipid-like molecules | Fatty Acyls |
| 439,3560 | 33,19 | ChemWalker | FQQMLVSIEXRTOS-UHFFFAOYSA-N | Organic compounds | Lipids and lipid-like molecules | Prenol lipids |
| 399,3475 | 33,20 | GNPS2 | PNXFYKPSFTUVDS-UHFFFAOYSA-N | Organic compounds | Organic acids and derivatives | Carboxylic acids and derivatives |
| 421,3290 | 33,20 | ChemWalker | LZEAIEMZXDHOKL-UHFFFAOYSA-N | Organic compounds | Organic oxygen compounds | Organooxygen compounds |
| 633,5056 | 33,22 | ChemWalker | UCLAKHRTNSBSRH-UHFFFAOYSA-N | Organic compounds | Lipids and lipid-like molecules | Glycerolipids |
| 429,3727 | 33,23 | ChemWalker | MTZGAGHRRROZFS-UHFFFAOYSA-N | Organic compounds | Lipids and lipid-like molecules | Fatty Acyls |
| 437,3600 | 33,33 | ChemWalker | OGARQWOPYHKQCQ-UHFFFAOYSA-N | Organic compounds | Lipids and lipid-like molecules | Steroids and steroid derivatives |
| 536,1650 | 33,34 | GNPS2 | SKUCQDOSGKINGP-YQMRLJPGSA-N | Organic compounds | Organic oxygen compounds | Organooxygen compounds |
| 447,3465 | 33,36 | ChemWalker | UTDDXZOYFWRGBB-UHFFFAOYSA-N | Organic compounds | Organic oxygen compounds | Organooxygen compounds |
| 449,3602 | 33,36 | ChemWalker | BFWBTIXVUPMKPV-UHFFFAOYSA-N | Organic compounds | Benzenoids | Benzene and substituted derivatives |
| 435,3446 | 33,42 | ChemWalker | USFJGINJGUIFSY-UHFFFAOYSA-N | Organic compounds | Lipids and lipid-like molecules | Steroids and steroid derivatives |
| 599,4274 | 33,47 | ChemWalker | BGKZYMYPODAKRB-UHFFFAOYSA-N | Organic compounds | Lipids and lipid-like molecules | Fatty Acyls |
| 435,3449 | 33,47 | ChemWalker | USFJGINJGUIFSY-UHFFFAOYSA-N | Organic compounds | Lipids and lipid-like molecules | Steroids and steroid derivatives |
| 399,1790 | 33,50 | ChemWalker | CBXOPGGSUUGLQL-UHFFFAOYSA-N | Organic compounds | Organoheterocyclic compounds | Indoles and derivatives |
| 377,1965 | 33,51 | ChemWalker | YPQOLUGHCAWRBO-UHFFFAOYSA-N | Organic compounds | Lipids and lipid-like molecules | Prenol lipids |
| 451,3747 | 33,54 | ChemWalker | JNMZYUSSPYVEHM-UHFFFAOYSA-N | Organic compounds | Lipids and lipid-like molecules | Steroids and steroid derivatives |
| 449,3611 | 33,59 | ChemWalker | XOIOXNHDDFEMFF-UHFFFAOYSA-N | Organic compounds | Organoheterocyclic compounds | Pyrrolopyrimidines |
| 435,3453 | 33,67 | ChemWalker | ZGOVEAPLOYTARH-UHFFFAOYSA-N | Organic compounds | Organoheterocyclic compounds | Pyrans |
| 449,3608 | 33,73 | ChemWalker | QGQMCLKYJMHDEZ-UHFFFAOYSA-N | Organic compounds | Organic oxygen compounds | Organooxygen compounds |
| 437,3600 | 33,75 | ChemWalker | IPUMBTXUZKFJIY-UHFFFAOYSA-N | Organic compounds | Organic oxygen compounds | Organooxygen compounds |
| 481,4015 | 33,76 | ChemWalker | AMSCMASJCYVAIF-UHFFFAOYSA-P | Organic compounds | Phenylpropanoids and polyketides | Macrolides and analogues |
| 605,4744 | 33,78 | ChemWalker | IWOYCFRIFNMRSY-UHFFFAOYSA-N | Organic compounds | Lipids and lipid-like molecules | Fatty Acyls |
| 449,3603 | 33,83 | ChemWalker | XOIOXNHDDFEMFF-UHFFFAOYSA-N | Organic compounds | Organoheterocyclic compounds | Pyrrolopyrimidines |
| 427,3803 | 33,84 | GNPS2 | SMKIPHDXOMDNDD-UHFFFAOYSA-N | Organic compounds | Organic acids and derivatives | Carboxylic acids and derivatives |
| 435,3450 | 33,84 | ChemWalker | ZGOVEAPLOYTARH-UHFFFAOYSA-N | Organic compounds | Organoheterocyclic compounds | Pyrans |
| 610,1834 | 33,87 | GNPS2 | OVMSOCFBDVBLFW-VHLOTGQHSA-N | Organic compounds | Lipids and lipid-like molecules | Prenol lipids |
| 465,3911 | 33,99 | ChemWalker | BKDLTBPGTMXGMR-UHFFFAOYSA-N | Organic compounds | Lipids and lipid-like molecules | Steroids and steroid derivatives |
| 548,5032 | 34,01 | SIRIUS | XAAAJVUZCYVFBM-RSOPFXSKSA-N | None | None | None |
| 553,4586 | 34,01 | SIRIUS | JZVCMKSGCLGCGX-UHFFFAOYSA-N | None | None | None |
| 443,2054 | 34,06 | ChemWalker | XZUPRNIANOZBMY-UHFFFAOYSA-N | Organic compounds | Lipids and lipid-like molecules | Prenol lipids |
| 421,2226 | 34,06 | ChemWalker | IZZZOEMEYJMVAL-UHFFFAOYSA-N | Organic compounds | Organoheterocyclic compounds | Furofurans |
| 441,3940 | 34,08 | ChemWalker | NVQGQHXFWUBOAU-UHFFFAOYSA-N | Organic compounds | Lipids and lipid-like molecules | Glycerolipids |
| 463,3759 | 34,08 | ChemWalker | VQOOTGXXVONQEH-UHFFFAOYSA-N | Organic compounds | Organic oxygen compounds | Organooxygen compounds |
| 465,2475 | 34,39 | ChemWalker | GLONBVCUAVPJFV-UHFFFAOYSA-N | Organic compounds | Lipids and lipid-like molecules | Steroids and steroid derivatives |
| 449,3723 | 34,40 | ChemWalker | YDZHZRCGUOPLTM-UHFFFAOYSA-N | Organic compounds | Lipids and lipid-like molecules | Prenol lipids |
| 684,2019 | 34,41 | GNPS2 | LLQBCHODNVGKSF-RVXDDWIJSA-N | Organic compounds | Organic oxygen compounds | Organooxygen compounds |
| 213,0526 | 34,63 | ChemWalker | BNLRKUSVMCIOGU-UHFFFAOYSA-N | Organic compounds | Organoheterocyclic compounds | Benzopyrans |
| 203,0228 | 34,8 | SIRIUS | DHOKASQYAFMGCQ-UHFFFAOYSA-N | Organic compounds | Organic acids and derivatives | Organic phosphonic acids and derivatives |
| 207,0177 | 34,79 | ChemWalker | NZQAQAUWFHMVEM-UHFFFAOYSA-M | Organic compounds | Phenylpropanoids and polyketides | Coumarins and derivatives |
| 232,9283 | 34,79 | SIRIUS | YQSJWKDRDGUZAA-UHFFFAOYSA-N | Organic compounds | Organoheterocyclic compounds | Diazines |
| 288,9212 | 34,79 | SIRIUS | YXCAIHHYOXKVFO-UHFFFAOYSA-N | Organic compounds | Organic acids and derivatives | Organic phosphonic acids and derivatives |
| 214,9174 | 34,80 | ChemWalker | YSHVYMPDKIBRIL-UHFFFAOYSA-N | Organic compounds | Organic 1,3-dipolar compounds | Allyl-type 1,3-dipolar organic compounds |
| 274,9060 | 34,80 | SIRIUS | DTGXOQKFBUUEQP-UHFFFAOYSA-N | Organic compounds | Organic acids and derivatives | Organic sulfonic acids and derivatives |
| 207,0177 | 34,80 | ChemWalker | JDAPQMINSZDTBM-UHFFFAOYSA-N | Organic compounds | Organoheterocyclic compounds | Triazines |
| 270,9104 | 34,80 | SIRIUS | QYUJBOJJEGIBCJ-UHFFFAOYSA-N | Organic compounds | Organic acids and derivatives | Organic sulfuric acids and derivatives |
| 238,8840 | 34,81 | SIRIUS | LNDYWKZPMGIUCR-UHFFFAOYSA-N | None | None | None |
| 256,8950 | 34,81 | SIRIUS | BVWCYPZMDYXDNO-UHFFFAOYSA-N | Organic compounds | Organic acids and derivatives | Organic thiophosphoric acids and derivatives |
| 240,8810 | 34,81 | ChemWalker | FRPHJINKMDMSPH-UHFFFAOYSA-N | Organic compounds | Benzenoids | Benzene and substituted derivatives |
| 203,0228 | 34,81 | SIRIUS | DHOKASQYAFMGCQ-UHFFFAOYSA-N | Organic compounds | Organic acids and derivatives | Organic phosphonic acids and derivatives |
| 221,0333 | 34,83 | ChemWalker | HEUOHXSHPVULTJ-UHFFFAOYSA-N | Organic compounds | Hydrocarbons | Unsaturated hydrocarbons |
| 294,8766 | 34,84 | SIRIUS | ZZYCPTONQZEZMR-UHFFFAOYSA-N | Organic compounds | Organic acids and derivatives | Carboxylic acids and derivatives |
| 318,8833 | 34,85 | SIRIUS | NBSLYSBFFUXFHL-UHFFFAOYSA-N | Organic compounds | Organoheterocyclic compounds | Thiophenes |
| 298,8718 | 34,85 | SIRIUS | PSBZFFLWXFWOAJ-UHFFFAOYSA-N | None | None | None |
| 312,8879 | 34,85 | SIRIUS | DVYJQHHXUDEMDD-UHFFFAOYSA-N | Organic compounds | Organoheterocyclic compounds | Pyridines and derivatives |
| 314,8849 | 34,85 | SIRIUS | KBOKWHCLOASPNC-UHFFFAOYSA-N | None | None | None |
| 316,8848 | 34,85 | SIRIUS | IVGWTJLQZWWWQJ-UHFFFAOYSA-N | None | None | None |
| 264,8777 | 34,86 | SIRIUS | JMGGDZWLZSAYRG-UHFFFAOYSA-N | Organic compounds | Organoheterocyclic compounds | Azoles |
| 212,9738 | 34,89 | SIRIUS | RPATYFZKHKXVFH-UHFFFAOYSA-N | Organic compounds | Organoheterocyclic compounds | Azoles |
| 200,9739 | 34,88 | ChemWalker | NOKPBJYHPHHWAN-UHFFFAOYSA-M | Organic compounds | Organic acids and derivatives | Carboxylic acids and derivatives |
| 214,9891 | 34,88 | SIRIUS | NZRSSORLUUMQEM-UHFFFAOYSA-N | Organic compounds | Organosulfur compounds | Sulfonyls |
| 224,9407 | 34,90 | SIRIUS | AHENLAPXTCWKIR-UHFFFAOYSA-N | Organic compounds | Organic acids and derivatives | Organic sulfonic acids and derivatives |
| 218,9848 | 34,90 | ChemWalker | JPOGVWCIWYTWAR-UHFFFAOYSA-N | Organic compounds | Organoheterocyclic compounds | Diazines |
| 204,9683 | 34,90 | ChemWalker | JIQXVIJARQLCOY-UHFFFAOYSA-N | Organic compounds | Benzenoids | Phenol ethers |
| 238,9558 | 34,90 | SIRIUS | XVMKDSWCXTYZLQ-UHFFFAOYSA-M | None | None | None |
| 232,9632 | 34,91 | SIRIUS | QYBFVDGIJIWFPD-UHFFFAOYSA-N | Organic compounds | Organic acids and derivatives | Carboxylic acids and derivatives |
| 242,9518 | 34,92 | SIRIUS | SEFNCZCTPWQJKT-UHFFFAOYSA-N | None | None | None |
