## Supplementary figures and images for "Characterization of a marine bacteria through a novel metabologenomics approach"

### SuppFig2_cinerubin.tiff

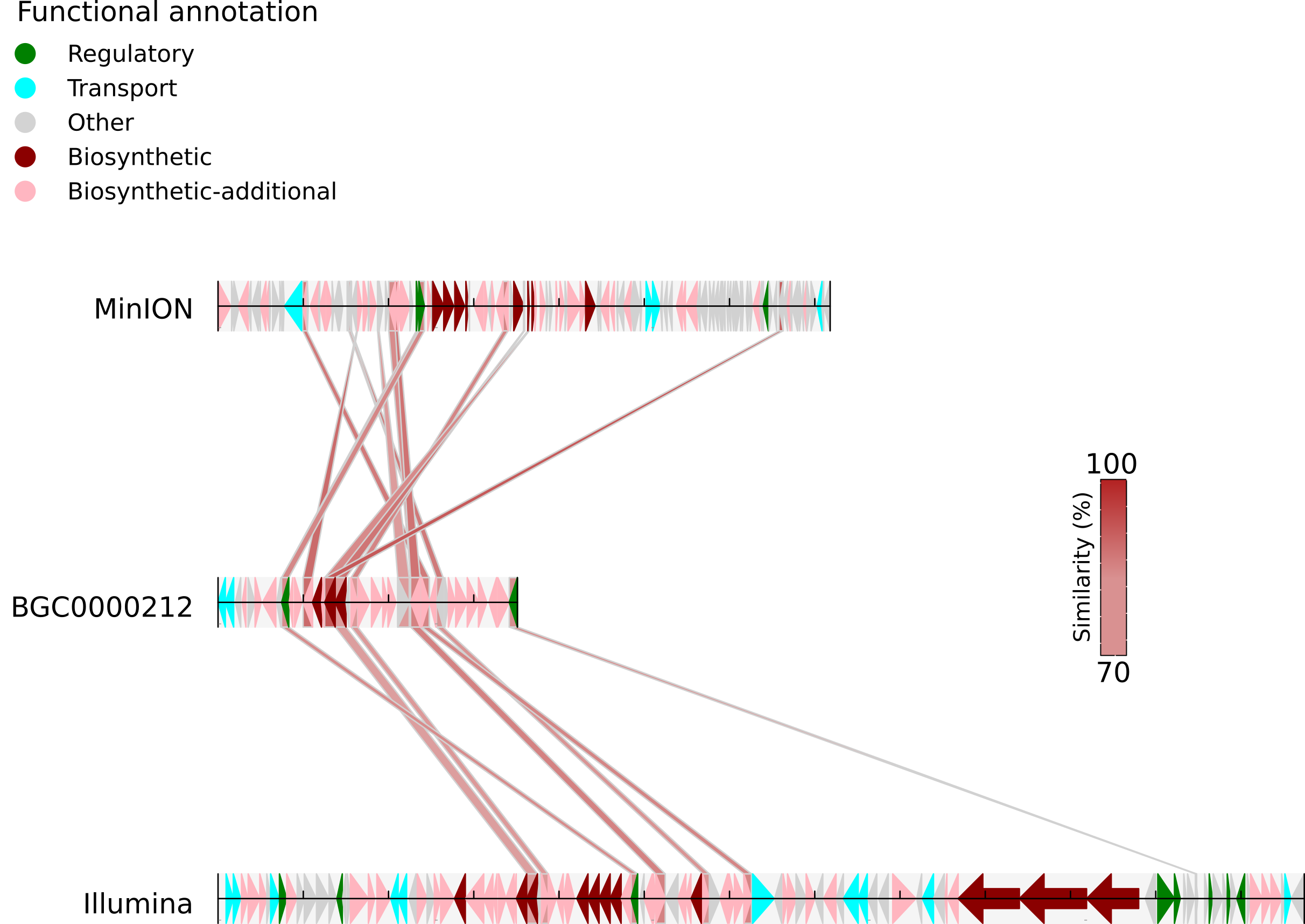

### SuppFig3_quinilodomicin.tiff

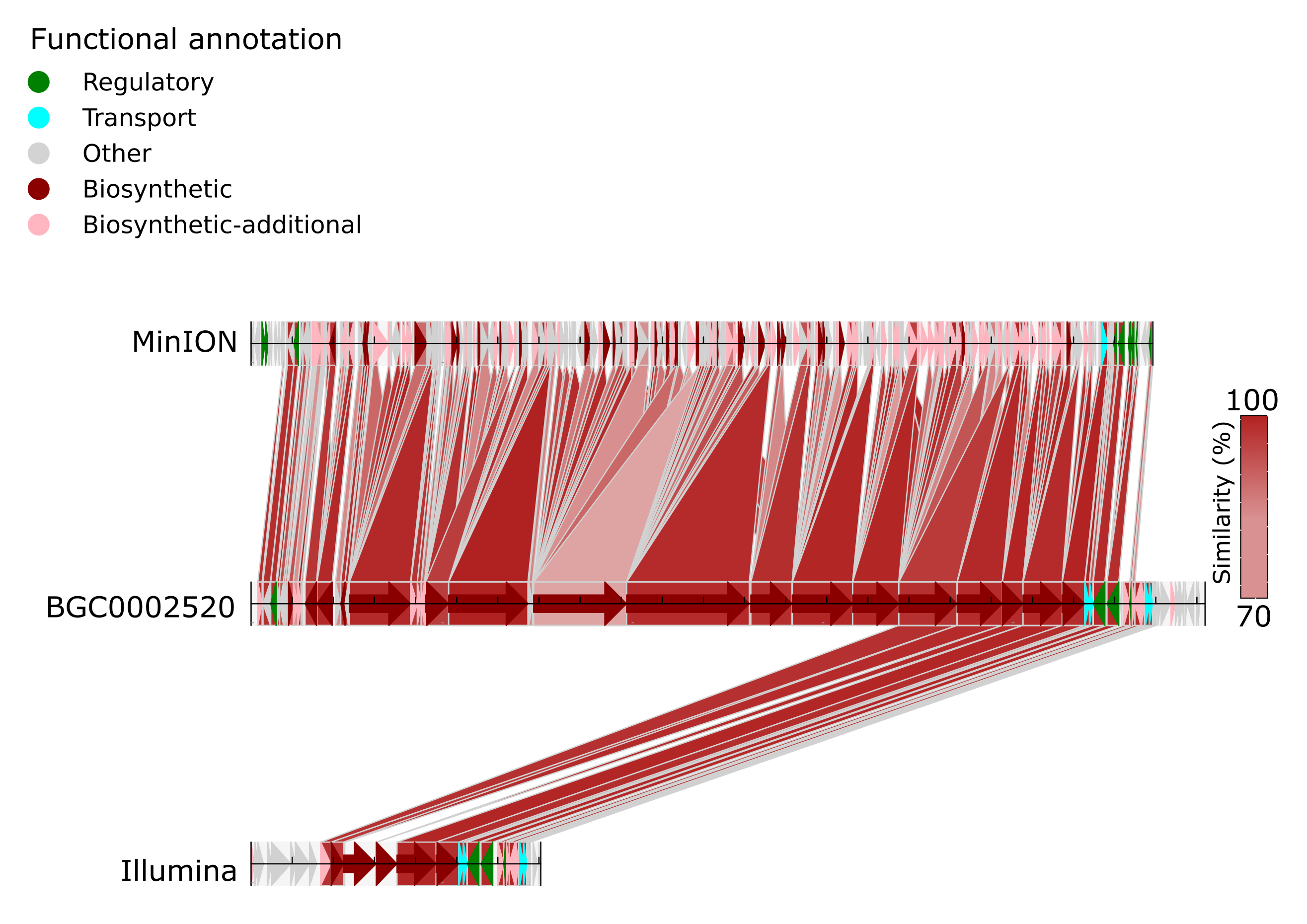

### SuppFig4_KEGG_pathway.tiff

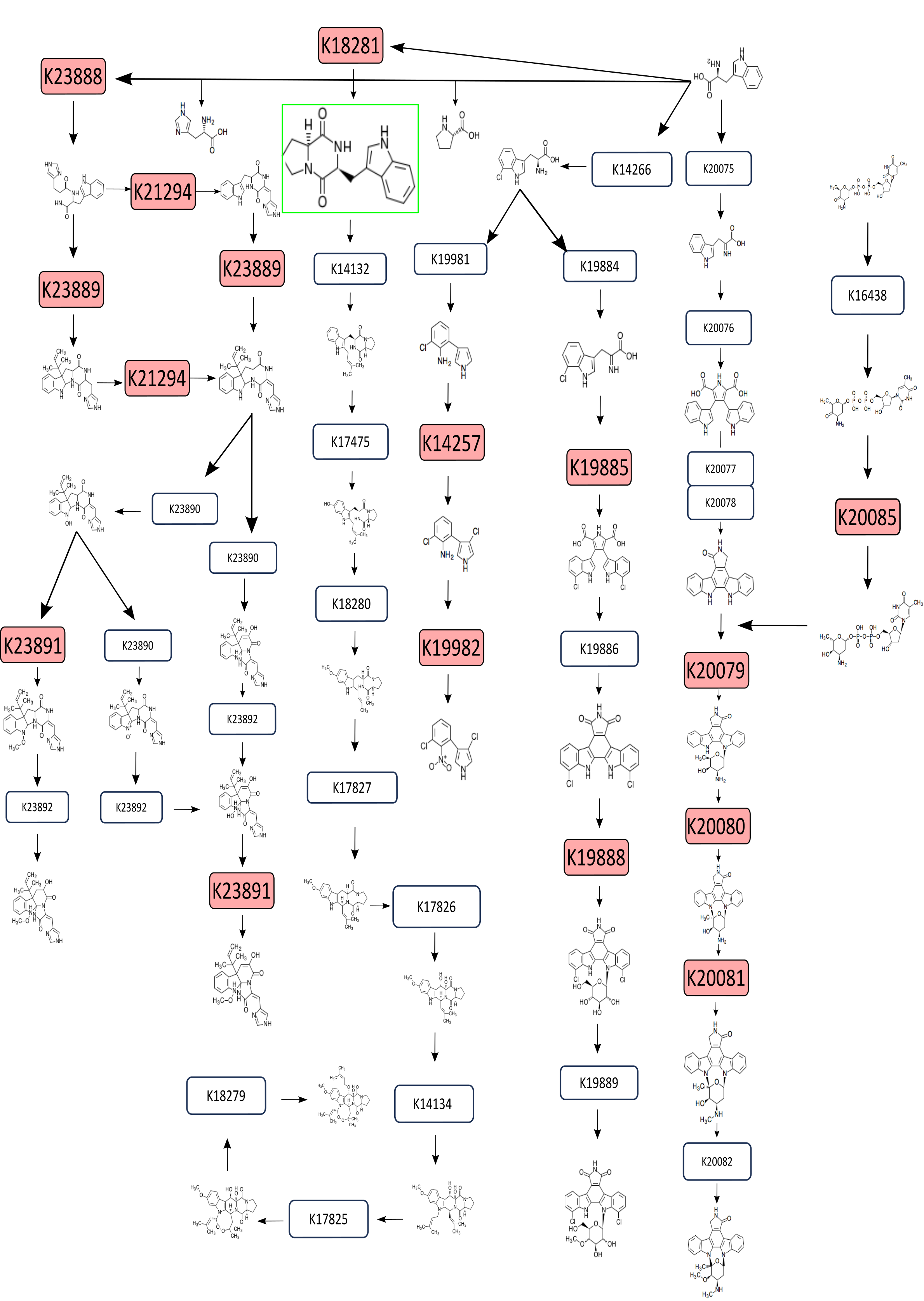
